# Functional insights from the GC-poor genomes of two aphid parasitoids, *Aphidius ervi* and *Lysiphlebus fabarum*

**DOI:** 10.1101/841288

**Authors:** Alice B. Dennis, Gabriel I. Ballesteros, Stéphanie Robin, Lukas Schrader, Jens Bast, Jan Berghöfer, Leo Beukeboom, Maya Belghazi, Anthony Bretaudeau, Jan Büllesbach, Elizabeth Cash, Dominique Colinet, Zoé Dumas, Patrizia Falabella, Jean-Luc Gatti, Elzemiek Geuverink, Joshua D. Gibson, Corinne Hertäg, Stefanie Hartmann, Emmanuelle Jacquin-Joly, Mark Lammers, Blas I. Lavandero, Ina Lindenbaum, Lauriane Massardier-Galata, Camille Meslin, Nicolas Montagné, Nina Pak, Marylène Poirié, Rosanna Salvia, Chris R. Smith, Denis Tagu, Sophie Tares, Heiko Vogel, Tanja Schwander, Jean-Christophe Simon, Christian C. Figueroa, Christoph Vorburger, Fabrice Legeai, Jürgen Gadau

**Author notes:** Joint first authors (Alice B. Dennis, Gabriel I. Ballesteros).

## Abstract

**Background:** Parasitoid wasps have fascinating life cycles and play an important role in trophic networks, yet little is known about their genome content and function. Parasitoids that infect aphids are an important group with the potential for biocontrol, and infecting aphids requires overcoming both aphid defenses and their defensive endosymbionts.

**Results:** We present the *de novo* genome assemblies, detailed annotation, and comparative analysis of two closely related parasitoid wasps that target pest aphids: *Aphidius ervi* and *Lysiphlebus fabarum* (Hymenoptera: Braconidae: Aphidiinae). The genomes are small (139 and 141 Mbp), highly syntenic, and the most AT-rich reported thus far for any arthropod (GC content: 25.8% and 23.8%). This nucleotide bias is accompanied by skewed codon usage, and is stronger in genes with adult-biased expression. AT-richness may be the consequence of reduced genome size, a near absence of DNA methylation, and age-specific energy demands. We identify expansions of F-box/Leucine-rich-repeat proteins, suggesting that diversification in this gene family may be associated with their broad host range or with countering defenses from aphids’ endosymbionts. The absence of some immune genes (Toll and Imd pathways) resembles similar losses in their aphid hosts, highlighting the potential impact of symbiosis on both aphids and their parasitoids.

**Conclusions:** These findings are of fundamental interest for insect evolution and beyond. This will provide a strong foundation for further functional studies including coevolution with respect to their hosts, the basis of successful infection, and biocontrol. Both genomes are available at https://bipaa.genouest.org.

## Background

Parasites are ubiquitously present across all of life (Poulin 2007; Windsor 1998). Their negative impact on host fitness can impose strong selection on hosts to resist, tolerate, or escape potential parasites. Parasitoids are a special group of parasites whose successful reproduction is fatal to the host (Godfray 1994; Quicke 2014). The overwhelming majority of parasitoid insects are hymenopterans that parasitize other terrestrial arthropods, and they are estimated to comprise up to 75% of the species-rich insect order Hymenoptera (Forbes *et al*. 2018; Godfray 1994; Heraty 2009; Pennacchio & Strand 2006). Parasitoid wasps target virtually all insects and developmental stages (eggs, larvae, pupae, and adults), including other parasitoids (Chen & van Achterberg 2018; Godfray 1994; Müller *et al*. 2004; Poelman *et al*. 2012). Parasitoid radiations appear to have coincided with those of their hosts (Peters *et al*. 2017), and there is ample evidence that host-parasitoid relationships impose strong reciprocal selection, promoting a dynamic process of antagonistic coevolution (Dupas *et al*. 2003; Kraaijeveld *et al*. 1998; Vorburger & Perlman 2018).

Parasitoids of aphids play an economically important role in biological pest control (Boivin *et al*. 2012; Heimpel & Mills 2017), and aphid-parasitoid interactions are an excellent model to study antagonistic coevolution, specialization, and speciation (Henter & Via 1995; Herzog *et al*. 2007). While parasitoids that target aphids have evolved convergently several times, their largest radiation is found in the braconid subfamily Aphidiinae, which contains at least 400 described species across 50 genera (Chen & van Achterberg 2018; Shi & Chen 2005). As koinobiont parasitoids, their development progresses initially in still living, feeding, and developing hosts, and ends with the aphids’ death and the emergence of adult parasitoids. Parasitoids increase their success with a variety of strategies, including host choice (Chau & Mackauer 2000; Łukasik *et al*. 2013), altering larval development timing (Martinez *et al*. 2016), injecting venom during stinging and oviposition, and developing special cells called teratocytes (Burke & Strand 2014; Colinet *et al*. 2014; Falabella *et al*. 2003; Poirié *et al*. 2014; Strand 2014). In response to strong selection imposed by parasitoids, aphids have evolved numerous defenses, including behavioral strategies (Gross 1993), immune defenses (Schmitz *et al*. 2012), and symbioses with heritable endosymbiotic bacteria whose integrated phages can produce toxins to hinder parasitoid success (Oliver *et al*. 2010; Oliver & Higashi 2018; Vorburger & Perlman 2018).

The parasitoid wasps *Lysiphlebus fabarum* and *Aphidius ervi* (*Braconidae*: Aphidiinae) are closely related endoparasitoids (Figure 1). In the wild both species are found infecting a wide range of aphid species although their host ranges differ, with *A. ervi* more specialized on aphids in the Macrosiphini tribe and *L. fabarum* on the Aphidini tribe (Kavallieratos *et al*. 2004; Monticelli *et al*. 2019). In both taxa, there is evidence that parasitoid success is hindered by the presence of defensive symbionts in the aphid haemocoel, including the bacteria *Hamiltonella*, *Regiella*, and *Serratia* (Oliver *et al*. 2003; Vorburger *et al*. 2010). Studies employing experimental evolution in both species have shown that wild-caught populations can counter-adapt to cope with aphids and the defenses of their endosymbionts, and that the coevolutionary relationships between parasitoids and the aphids’ symbionts likely fuel diversification of both parasitoids and their hosts (Dennis *et al*. 2017; Dion *et al*. 2011; Rouchet & Vorburger 2014). While a number of parasitoid taxa are known to inject viruses and virus-like particles into their hosts, there is thus far no evidence that this occurs in parasitoids that target aphids; emerging studies have identified abundant RNA viruses in *L. fabarum* (Lüthi *et al*. submitted; Obbard *et al*. in revision), but whether this impacts their ability to parasitize is not yet fully understood.

**Figure 1.**
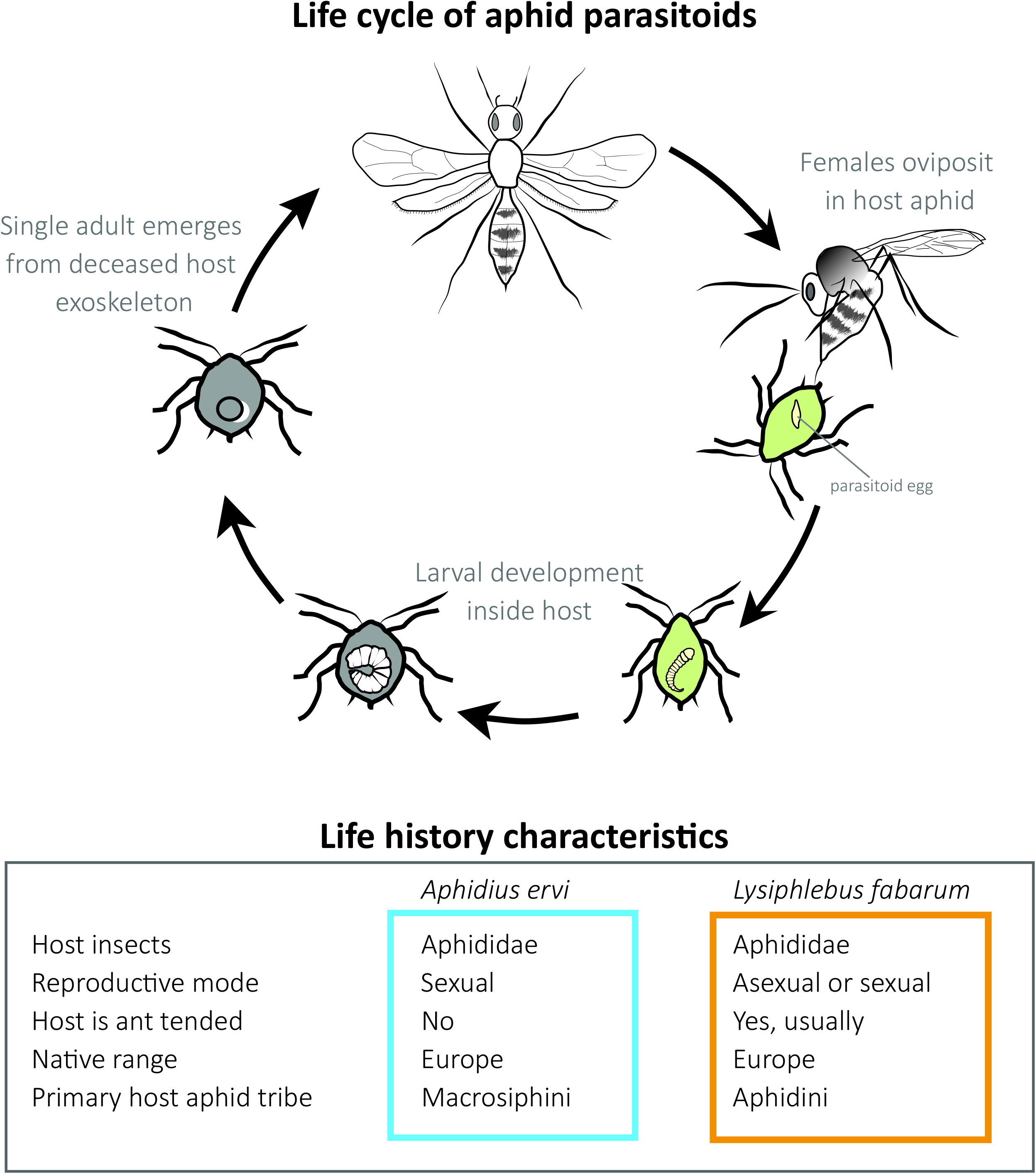
Aphid parasitoid life cycle: Generalized life cycle of *Aphidius ervi* and *Lysiphlebus fabarum*, two different parasitoid wasps that target aphid hosts.

These two closely related parasitoids differ in several important life history traits, and are expected to have experienced different selective regimes as a result. *Aphidius ervi* is has successfully been introduced widely (Nearctic, Neotropics) as a biological control agent (far more than *L. fabarum*). Studies on both native and introduced populations of *A. ervi* have shown ongoing evolutionary processes with regard to host preferences, gene flow, and other life history components (Henry *et al*. 2008; Hufbauer *et al*. 2004; Zepeda-Paulo *et al*. 2015; Zepeda-Paulo *et al*. 2013). *A. ervi* is known to reproduce only sexually, whereas *L. fabarum* is capable of both sexual and asexual reproduction. In fact, wild *L. fabarum* populations are more commonly composed of asexually reproducing (thelytokous) individuals (Sandrock *et al*. 2011). In asexual populations, diploid *L. fabarum* females produce diploid female offspring via central fusion automixis (Belshaw & Quicke 2003). While they are genetically differentiated, sexual and asexual populations appear to maintain gene flow and thus both reproductive modes and genome-wide heterozygosity are maintained in the species as a whole (Mateo Leach *et al*. 2009; Sandrock *et al*. 2011; Sandrock & Vorburger 2011). *Aphidius. ervi* and *L. fabarum* are also expected to have experienced different selective regimes with regard to their cuticular hydrocarbon profiles and chemosensory perception. *Lysiphlebus* target aphid species that are ant-tended, and ants are known to prevent parasitoid attacks on “their” aphids (Rasekh *et al*. 2010). To counter ant defenses, *L. fabarum* has evolved the ability to mimic the cuticular hydrocarbon profile of the aphid hosts (Liepert & Dettner 1993, 1996). With this, they are able to circumvent ant defenses and access this challenging ecological niche, from which they also benefit nutritionally; they are the only parasitoid species thus far documented to behaviorally encourage aphid honeydew production and consume this high-sugar reward (Rasekh *et al*. 2010; Völkl 1992; Völkl 1997).

We present here the genomes of *A. ervi* and *L. fabarum*, assembled *de novo* using a hybrid sequencing approach. The two genomes are highly syntenic and strongly biased towards AT nucleotides. We have examined GC content in the context of host environment, nutrient limitation, and gene expression. By comparing these two genomes we identify key functional specificities in genes underlying venom composition, oxidative phosphorylation, cuticular hydrocarbon composition, and chemosensory perception. In both species, we identify losses in key immune genes and an apparent lack of key DNA methylation machinery. These are functionally important traits associated with success infecting aphids and the evolution of related traits across all of Hymenoptera.

## Results and Discussion

### Two de novo genome assemblies

The genome assemblies for *A. ervi* and *L. fabarum* were constructed using hybrid approaches that incorporated high-coverage short read (Illumina) and long-read (Pac Bio) sequencing, but were assembled with different parameters (Supplementary Tables 1, 2). This produced two high quality genome assemblies (*A. ervi* N50 = 581kb, *L. fabarum* N50 = 216kb) with similar total lengths (*A. ervi*: 139MB, *L. fabarum*: 141MB) but different ranges of scaffold-sizes (Table 1, Supplementary Table 3). These assembly lengths are within previous estimates of 110-180Mbp for braconids, including *A. ervi* (Ardila-Garcia *et al*. 2010; Hanrahan & Johnston 2011). Both assemblies are available in NCBI (SAMN13190903-4) and can be accessed via the BioInformatics Platform for Agroecosystem Arthropods (BIPAA, https://bipaa.genouest.org), which contains the full annotation reports, predicted genes, and can be searched via both keywords and blast.

**Table 1:**
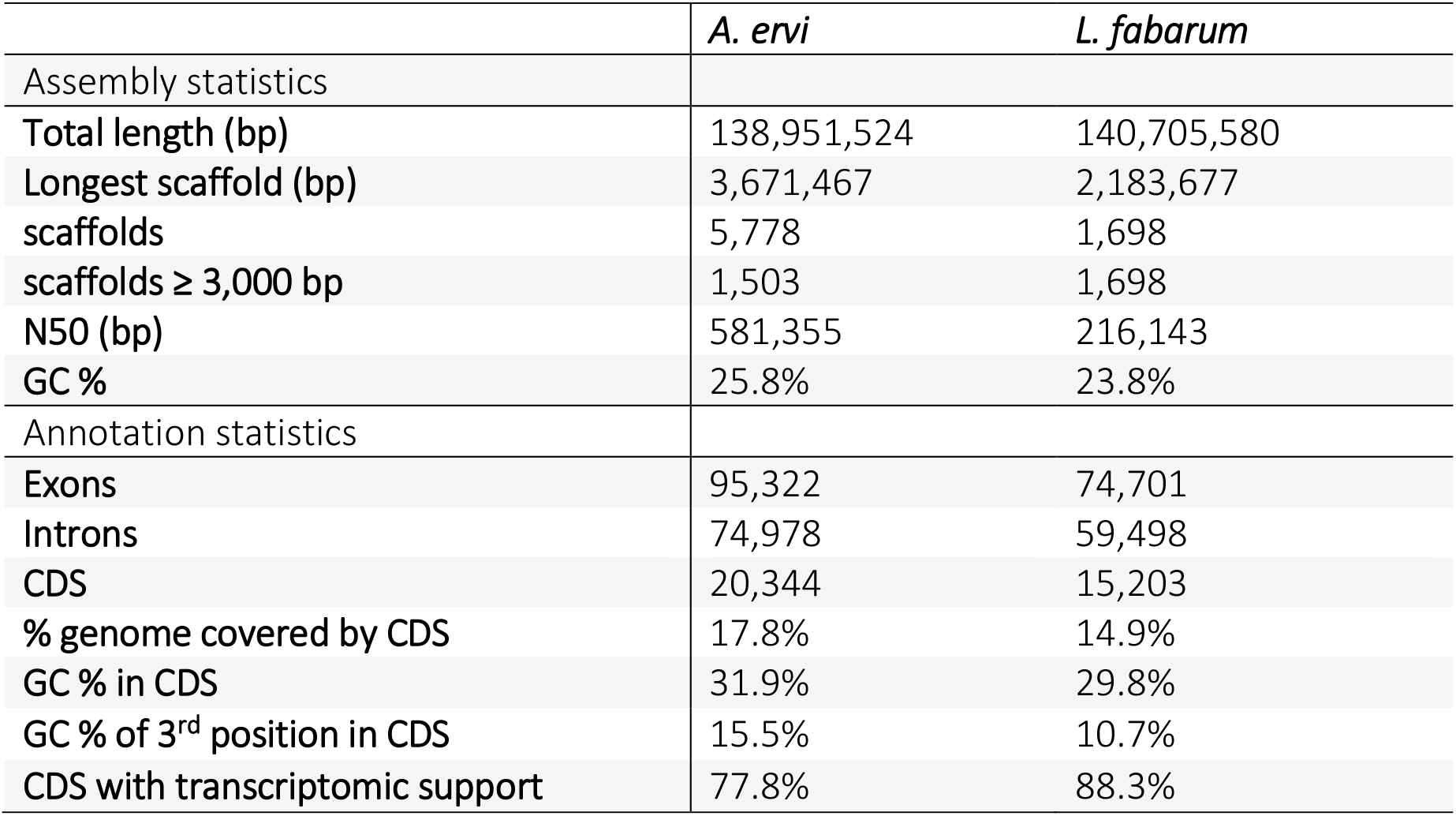
Assembly and draft annotation statistics.

|  | *A. ervi* | *L. fabarum* |
| --- | --- | --- |
| Assembly statistics |  |  |
| Total length (bp) | 138,951,524 | 140,705,580 |
| Longest scaffold (bp) | 3,671,467 | 2,183,677 |
| scaffolds | 5,778 | 1,698 |
| scaffolds ≥ 3,000 bp | 1,503 | 1,698 |
| N50 (bp) | 581,355 | 216,143 |
| GC % | 25.8% | 23.8% |
| Annotation statistics |  |  |
| Exons | 95,322 | 74,701 |
| Introns | 74,978 | 59,498 |
| CDS | 20,344 | 15,203 |
| % genome covered by CDS | 17.8% | 14.9% |
| GC % in CDS | 31.9% | 29.8% |
| GC % of 3<sup>rd</sup> position in CDS | 15.5% | 10.7% |
| CDS with transcriptomic support | 77.8% | 88.3% |

We constructed linkage groups for the *L. fabarum* scaffolds using phased SNPs from the haploid (male) sons of a single female wasp from a sexually reproducing population. This placed the 297 largest scaffolds (>50% of the nucleotides, Supplementary Table 5, Supplementary Figure 1, Additional File 1) into the expected six chromosomes (Belshaw & Quicke 2003). With this largely contiguous assembly, we show that the two genomes are highly syntenic, with >60k links in alignments made by NUCmer (Kurtz *et al*. 2004) and >350 large syntenic blocks that match the six *L. fabarum* chromosomes to 28 *A. ervi* scaffolds (Supplementary Figures 2 and 3).

Within the two assemblies, we used the Maker2 annotation pipeline to predict coding genes (CDS) for the two genomes, and these were functionally annotated against the NCBI *nr* database (NCBI), matches to gene ontology (GO) terms, and predictions for known protein motifs, signal peptides, and transmembrane domains (Supplemental Table 6). In *A. ervi* there were 20,344 predicted genes comprising 27.8Mbp, while in *L. fabarum* there were 15,203 genes across 21.9 Mbp (Table 1). These numbers are on par with those predicted in other hymenopteran genomes (Table 2), and comparisons among taxa suggest that the lower number of predicted genes in *L. fabarum* are more likely due to their loss than to a gene gain in *A. ervi*. However, it is important to recognize that predictive annotation is imperfect and any missing genes should be specifically screened with more rigorous methods. In both species, there was high transcriptomic support for the predicted genes (77.8% in *A. ervi* and 88.3% in *L. fabarum*). The two genome annotations appear to be largely complete; at the nucleotide level, we could match 94.8% (*A. ervi*) and 76.3% (*L. fabarum*) of the 1,658 core orthologous BUSCO genes for Insecta in both species (Supplementary Table 4). Within the predicted genes, protein-level matches to the BUSCO genes were improved in *L. fabarum* (95.9%) and slightly lower for *A. ervi* (93.7%). These numbers suggest that low GC content did not negatively impact gene prediction (Supplementary Table 4).

**Table 2:** Assembly summary statistics compared to other parasitoid genomes. All species are from the family Braconidae, except for N. vitripennis (Pteromalidae). Protein counts from the NCBI genome deposition.

| Parasitoid species | Assembly | Total Length (Mbp) | Scaffold Count | Scaffold N50 (bp) | Predicted genes (CDS) | GC (%) | NCBI BioProject |
| --- | --- | --- | --- | --- | --- | --- | --- |
| *Aphidius ervi* | A. ervi_v3 | 139.0 | 5,778 | 581,355 | 20,344 | 25.8 | *This paper* |
| *Lysiphlebus fabarum* | L. fabarum_v1 | 140.7 | 1,698 | 216,143 | 15,203 | 23.8 | *This paper* |
| *Fopius arisanus* | ASM80636v1 | 153.6 | 1,042 | 51,867 | 18,906 | 39.4 | PRJNA258104 (Geib *et al.* 2017) |
| *Diachasma alloeum* | Dall1.0 | 388.8 | 3,968 | 44,932 | 19,692 | 39.1 | PRJNA284396 (Tvedte *et al.* 2019) |
| *Microplitis demolitor* | Mdem 2 | 241.2 | 1,794 | 27,508 | 18,586 | 33.1 | PRJNA251518 (Burke *et al.* 2018) |
| *Cotesia vestalis* | ASM95615v1 | 186.1 | 9,156 | 46,055 | - | 30.4 | PRJNA271135 |
| *Macrocentrus cingulum* | MCINOGS1.0 | 127.9 | 12,056 | 65,089 | 11,993 | 35.6 | PRJNA361069 (Yin *et al.* 2018) |
| *Nasonia vitripennis* | Nvit_2.1 | 295.7 | 6,169 | 18,840 | 24,891 | 40.6 | PRJNA13660 (Werren *et al.* 2010) |

A survey of transposable Elements (TEs) identified a similar overall number of putative TE elements in the two assemblies (*A. ervi*: 67,695 and *L. fabarum*: 60,306, Supplementary Table 7). Despite this similarity, the overall genomic coverage by TEs is larger in *L. fabarum* (41%, 58 Mbp) than in A. *ervi* (22%, 31 Mbp) and they differ in the TE classes that they contain (Supplementary Table 7, Supplementary Figures 4, 5). The spread of reported TE coverage in arthropods is quite large, even among *Drosophila* species (ca. 2.7% - 25%, Drosophila 12 Genomes *et al*. 2007). Within parasitoids, reported TE content also varies, and relatively low coverage in the parasitoid *Macrocentrus cingulum* in comparison to *Nasonia vitripennis* (24.9% vs 40.6% Yin *et al*. 2018) was attributed the smaller genome size of *M. cinculum* (127.9Mbp and 295.7Mbp, respectively, Table 3). However, the variation we observe here suggests that differences in predicted TE content may be evolutionary quite labile, even within closely related species with the same genome size.

**Table 3:**
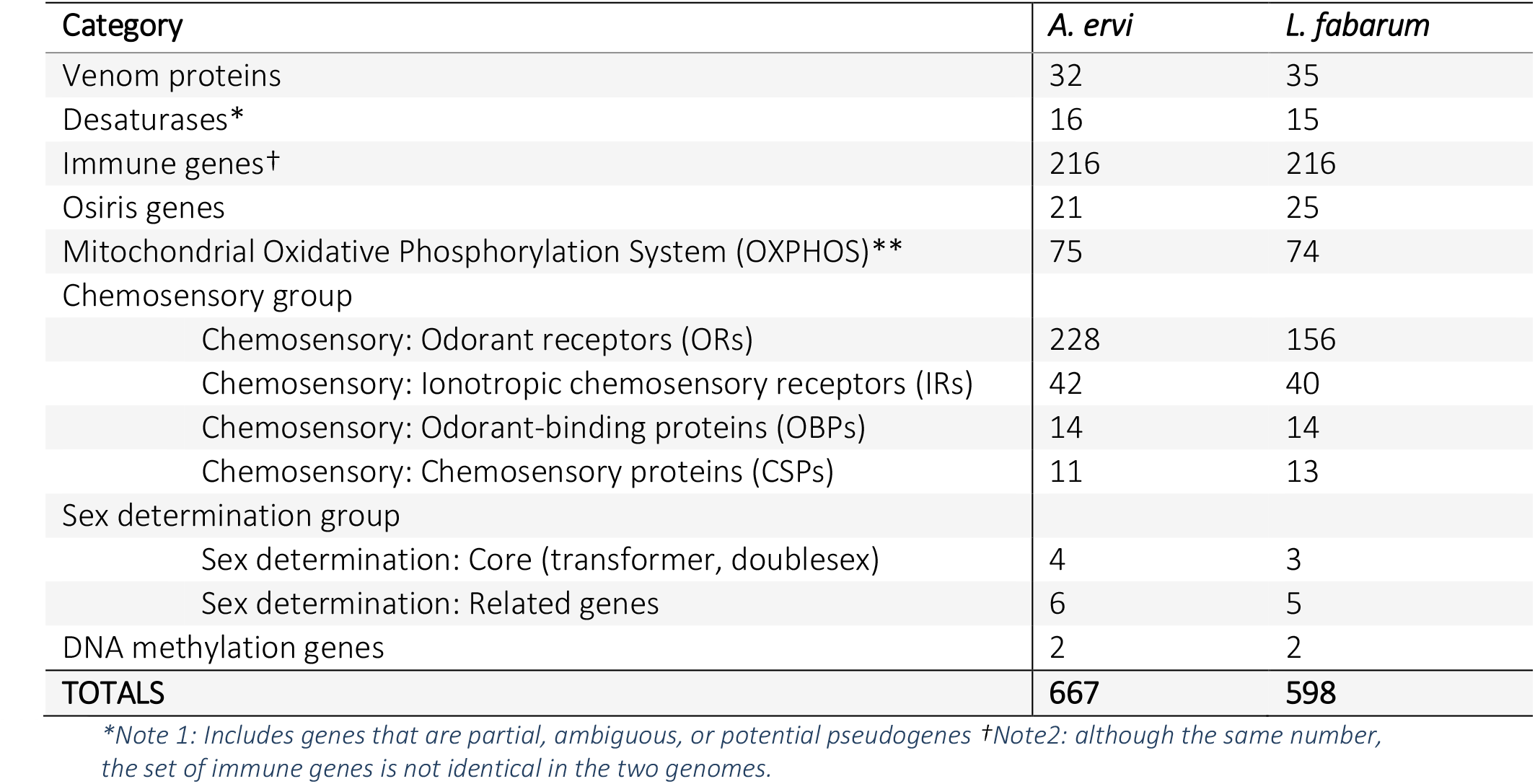
Summary of manual curations of select gene families in the two parasitoid genomes.

| Category | *A. ervi* | *L. fabarum* |
| --- | --- | --- |
| Venom proteins | 32 | 35 |
| Desaturases* | 16 | 15 |
| Immune genes† | 216 | 216 |
| Osiris genes | 21 | 25 |
| Mitochondrial Oxidative Phosphorylation System (OXPHOS)** | 75 | 74 |
| Chemosensory group |  |  |
| Chemosensory: Odorant receptors (ORs) | 228 | 156 |
| Chemosensory: Ionotropic chemosensory receptors (IRs) | 42 | 40 |
| Chemosensory: Odorant-binding proteins (OBPs) | 14 | 14 |
| Chemosensory: Chemosensory proteins (CSPs) | 11 | 13 |
| Sex determination group |  |  |
| Sex determination: Core (transformer, doublesex) | 4 | 3 |
| Sex determination: Related genes | 6 | 5 |
| DNA methylation genes | 2 | 2 |
| **TOTALS** | **667** | **598** |

\*Note 1: Includes genes that are partial, ambiguous, or potential pseudogenes †Note2: although the same number, the set of immune genes is not identical in the two genomes.

### GC content

The *L. fabarum* and *A. ervi* genomes are the most GC-poor of insect genomes sequenced to date (GC content: 25.8% and 23.8% for *A. ervi* and *L. fabarum*, respectively, Table 3, Supplementary Figure 6). This nucleotide bias is accompanied by strong codon bias in the predicted genes, meaning that within the possible codons for each amino acid, the two genomes are almost universally skewed towards the codon(s) with the lowest GC content (measured as Relative Synonymous Codon Usage, RSCU, Figure 2). These patterns are much more extreme than RSCU found in other hymenopterans, which are known to prefer codons that end in –A or –U (Behura & Severson 2013). This codon bias has functional consequences; work in other taxa has shown that codon usage is tied to both expression efficiency and mRNA stability (Barahimipour *et al*. 2015).

**Figure 2.**
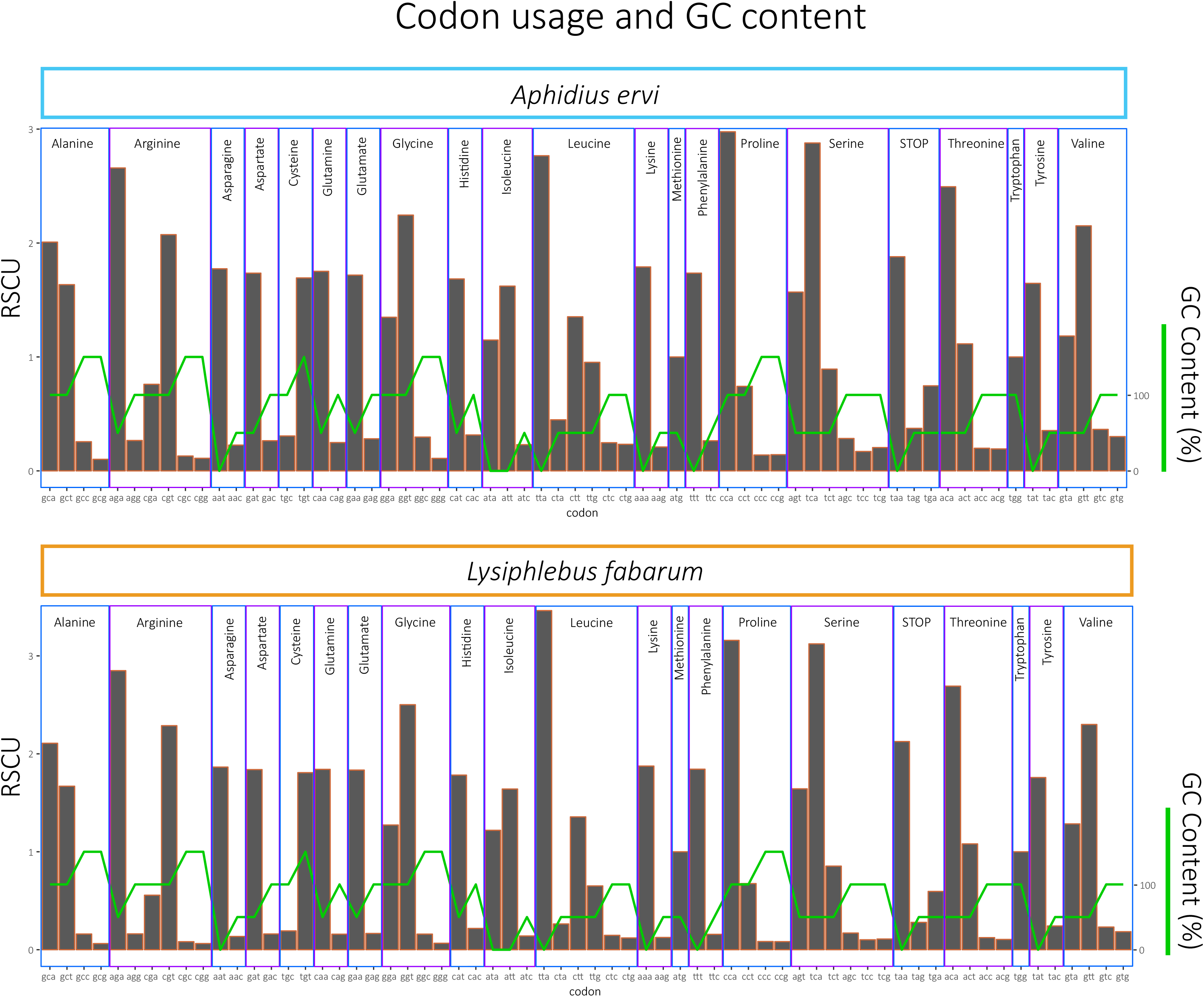
Codon usage in predicted genes: Proportions of all possible codons, as used in the predicted genes in *A. ervi* (top) and *L. fabarum* (bottom). Codon usage was measured as relative synonymous codon usage (RSCU), which scales usage to the number of possible codons for each amino acid (RSCU). Codons are listed at the bottom and are grouped by the amino acid that they encode. The green line depicts GC contend (%) of the codon.

Low GC content could be a consequence of the relatively small size of these genomes. Genome size and GC content are positively correlated in a diverse set of taxa including bacteria (Almpanis *et al*. 2018; McCutcheon *et al*. 2009), plants (Šmarda *et al*. 2014; Veleba *et al*. 2016), and vertebrates (Vinogradov 1998). This widespread pattern may be driven by GC-rich repetitive elements that are more abundant in larger genomes, stronger selection on thermal stability in larger genomes, or thermal stability associated with the environment (Šmarda *et al*. 2014; Vinogradov 1998). The apparent lack of DNA methylation in this system may also contribute to low GC content (see below and Bewick *et al*. 2017). Methylation is a stabilizing factor with regard to GC content (Mugal *et al*. 2015), so its absence could relax selection on GC content and allow it to decline. However, neither the absence of methylation nor codon bias are unique to these taxa, suggesting that some additional selective factors or genetic drift may have further shaped the composition of these two genomes.

We used two approaches to investigate whether environmental constraints could drive extremely low GC content, but found no evidence for such constraints. There is reason to expect that environment could contribute to the low GC content of these genomes; in taxa including bacteria (Foerstner *et al*. 2005) and plants (Šmarda *et al*. 2014) the environment has been shown to influence GC content via limitation in elements including nitrogen. These two wasps parasitize aphids exclusively, and aphids themselves have relatively low genome-wide GC content. This includes the pea aphid (*Acyrtosiphon pisum*), which is a frequent host of *A. ervi* and also has notably low GC content (29.8%, Li *et al*. 2019). This is not limited to *A. pisum*, with other aphid genomes’ GC content ranging between 26.8% - 30% (Additional File 2), perhaps related to their high-sugar, low-nitrogen, sap diet. One way to explore the restrictions imposed by nutrient limitation is to look at the expressed genes, since selective pressure should be higher for genes that are more highly expressed (Ran & Higgs 2010; Seward & Kelly 2016). For our first test, we explored potential constraints in the most highly expressed genes in both genomes. In both species, the most highly expressed 5% of genes had higher GC content and higher nitrogen content, although the higher number of nitrogen molecules in G’s and C’s means that these two measures cannot be entirely disentangled (Additional File 3, Supplementary Figure 7). This is in line with observations across many taxa, and with the idea that GC-rich mRNA has increased expression via its stability and secondary structure (Kudla *et al*. 2009; Plotkin & Kudla 2011). For a second approach to examining constraints, we compared codon usage between our genomes and taxa associated with this parasitoid-host-endosymbiont system (Supplementary Table 8). We found no evidence of similarity in codon usage (scaled as RSCU) nor in nitrogen content (scaled per amino acid) between parasitoids and host aphids, the primary endosymbionts *Buchnera* nor, with the secondary endosymbiont *Hamiltonella* (Supplementary Figures 8-10). Together, these tests do not support environmental constraints as the driver of low GC content in these two genomes.

In contrast, we did find evidence for reduced GC content in genes expressed at different parasitoid life-history stages. We found higher GC content in larvae-biased genes in *L. fabarum* (Figure 3). This was true when we compared the 10% most highly expressed genes in adults (32.6% GC) and larvae (33.2%, p=1.2e-116, Figure 3, Additional File 3), and this pattern holds even more strongly for genes that are differentially expressed between adults (upregulated in adults: 28.7% GC) and larvae (upregulated in larvae: 30.7% GC, p=2.2e-80. Note that the most highly expressed genes overlap partially with those that are differentially expressed, Additional File 3). At the same time, we found no evidence that nitrogen content differs in either of these comparisons (Figure 3). While the magnitude of these differences is not very large, subtle differences in gene content are hypothesized to be the result of selection in other systems (Acquisti *et al*. 2009). It seems plausible that GC content differences among genes expressed at different life history stages could be selected in a process analogous to the small changes in gene expression that are linked to large phenotypic differences within and between species (Romero *et al*. 2012). One explanation for lower GC content in adult-biased genes could be differences in energy demands and availability of resource across life stages. Given the extreme codon bias in these genomes (Figure 2), using codons that match this bias is expected to be more efficient and accurate, resulting in lower energy consumption and faster turnover (Chaney & Clark 2015; Galtier *et al*. 2018; Kudla *et al*. 2006; Rao *et al*. 2013). Expressing AT-rich genes is slightly more energy-efficient in itself, and this could favor otherwise neutral mutations from GC to AT (Rocha & Danchin 2002). There is good motivation for adults to have a greater demand for energy efficiency. Adult parasitoids usually feed on carbohydrate rich but protein and lipid poor resources like nectar, while performing costly tasks including flying, mating, and laying eggs. Meanwhile, parasitoid larvae are feeding on their aphid host’s tissue, and likely benefit further from nutrients coming from the aphids’ endosymbionts, while their only task is to grow as fast as possible (Cheng *et al*. 2011; Miao *et al*. 2004; Pennacchio *et al*. 1999).

**Figure 3.**
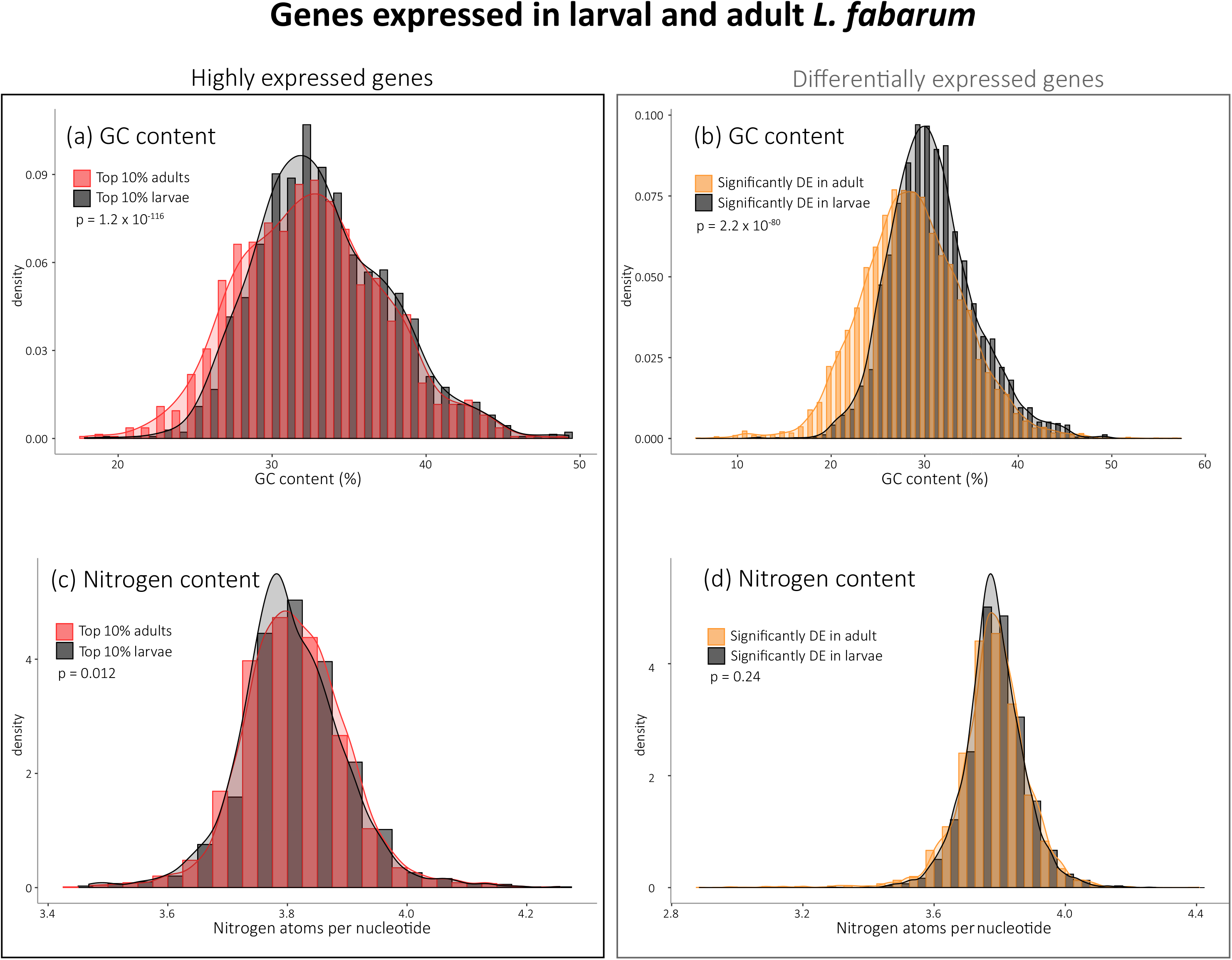
GC and nitrogen content of expressed genes: We observe significant differences (p-values from two-sided t-test) in the GC content between adult and larval *L. fabarum* in: (A) the most highly expressed 10% of the genes and (B) genes that are differentially expressed between adults and larvae. In contrast, there is no difference in the nitrogen content of the same set of genes (C, D).

This supports the idea that selection at the level of gene expression is shaping the GC content of these genomes. Nonetheless, further work should more explicitly test both nutrient limitation and how selective pressures differ across life-history stages. While we do not have the power to test for GC-biased gene conversion with two taxa, the even lower third position GC content (15.5% and 10.7%, Table 1) suggests that this should be tested in relation to other parasitoids (Galtier *et al*. 2018). Further explanations to be considered include effective population size, translational efficiency, and mutational bias (Behura & Severson 2013; Bentele *et al*. 2013; Galtier *et al*. 2018). Altogether, these patterns raise important questions about how codon biases impact genome content, and whether synonymous mutations are always functionally neutral (Plotkin & Kudla 2011; Powell & Moriyama 1997).

### Orphan genes in the assembly

To examine genes that may underlie novel functional adaptation, we identified sequences that are unique within the predicted genes in the *A. ervi* and *L. fabarum* genomes. We defined orphan genes as predicted genes with transcriptomic support and with no identifiable homology based on searches against the NCBI *nr*, *nt*, and Swissprot databases. With this, we identified 2,568 (*A. ervi*, Additional File 4) and 968 (*L. fabarum*, Additional File 5) putative orphans (Supplementary Table 9). The evolutionary origin of these orphan genes is not known (Gold *et al*. 2018; Van Oss & Carvunis 2019), but their retention or evolution could be important to understanding specific functions or traits in these taxa. The higher number of orphan genes in *A. ervi* partially explains the absolute difference in the number of annotated genes between both taxa.

### Gene family expansions

To examine gene families that may have undergone expansions in association with functional divergence and specialization, we identified groups of orthologous genes that have increased and decreased in size in the two genomes, relative to one another. We identified these species-specific gene-family expansions using the OMA standalone package (Altenhoff *et al*. 2018). OMA predicted 8,817 OMA groups (strict 1:1 orthologs) and 8,578 HOGs (Hierarchical Ortholog Groups, Additional File 6). Putative gene-family expansions would be found in the predicted HOGs, because they are calculated to allow for >1 member per species. Among these, there were more groups in which *A. ervi* possessed more genes than *L. fabarum* (865 groups with more genes in *A. ervi*, 223 with more in *L. fabarum*, Supplementary Figure 11, Additional File 6). To examine only the largest gene-family expansions, we looked further at the HOGs containing >20 genes (10 HOG groups, Supplementary Figure 12). Strikingly, the four largest expansions were more abundant in *A. ervi* and were all identified as F-box proteins/ Leucine-rich-repeat proteins (*LRR*, total: 232 genes in *A. ervi* and 68 in *L. fabarum*, Supplementary Figure 12, Additional File 6). This signature of expansion does not appear to be due to fragmentation in the *A. ervi* assembly: the size of scaffolds containing *LRR*s is on average larger in *A. ervi* than in *L. fabarum* (Welch two-sampled t-test, p=0.001, Supplementary Figure 13).

The *LRR*s are a broad class of proteins associated with protein-protein interactions, including putative venom components in these parasitoids (Colinet *et al*. 2014). *LRRs* belong to a larger category of leucine rich repeat pattern recognition receptor proteins, which are an important component of innate immunity and cell-surface recognition of bacterial intruders and include toll-like receptors in insects (Soanes & Talbot 2010; Takeda & Akira 2005). While the functions of these proteins are diverse, expansion in F-box/*LRR* proteins has been shown to have specific function in immunity in parasitic insects. In the Hessian fly (*Mayetiola destructor*), fly-encoded F-box/*LRR* proteins bind with plant-encoded proteins to form a complex that blocks the plant’s immune defenses against the parasitic fly (Zhao *et al*. 2015). Thus, we hypothesize that this class of proteins has expanded in these parasitoids in relation to recognizing the diverse bacterial defenses of their aphid hosts. Under this hypothesis, we argue that expansion of F-box/*LRR* proteins contributes to the broad host recognition in both species, and that their greater abundance in *A. ervi* may be associated with a recent arms race with respect to the immune defenses and protective endosymbionts of their host aphids.

The six largest gene families that were expanded in *L. fabarum*, relative to *A. ervi,* were less consistently annotated. Interestingly, they contained two different histone proteins: Histone H2B and H2A (Supplementary Figure 12). All eukaryotic genomes examined to date contain multiple histone genes for the same histone variants found in humans (e.g. 22 genes for H2B or 16 genes for H2A in humans, Singh *et al*. 2018), and it has recently been suggested that these histone variants are not functionally equivalent but rather play a role in chromatin regulation (Singh *et al*. 2018). Hence, these variants could also play a role in several *L. fabarum* specific traits, including the switch from sexual to asexual reproduction (thelytoky); in mammals, sex determination has been linked to regulation via histone modification (Kuroki *et al*. 2013).

### Venom proteins

Venom injected at oviposition is crucial for successful reproduction in most parasitoid wasp species (Moreau & Asgari 2015; Poirié *et al*. 2014). The venom of *A. ervi* was previously analyzed using a combined transcriptomic and proteomic approach (Colinet *et al*. 2014), and we applied similar methods here to compare the venom composition in *L. fabarum*. The venom gland in *L. fabarum* is morphologically different from *A. ervi* (Supplementary Figure 14). A total of 35 *L. fabarum* proteins were identified as putative venom proteins using 1D gel electrophoresis and mass spectrometry, combined with transcriptomic and the genome data (Supplementary Figure 15, Additional File 7, Dennis *et al*. 2017). These putative venom proteins were identified based on predicted secretion (for complete sequences) and the absence of a match to typical cellular proteins (e.g. actin, myosin). To match the analysis between the two taxa, the previous *A. ervi* venom data (Colinet *et al*. 2014) was analyzed using the same criteria as *L. fabarum*. This identified 32 putative venom proteins in *A. ervi* (Additional File 7).

Although these two species differ in their host range (Kavallieratos *et al*. 2004), comparison of venom proteins between species revealed that more than 50% of the proteins are shared between species (Figure 4A and Additional File 7), corresponding to more than 70% of the putative function categories that were predicted (Figure 4B and Additional File 7). Among venom proteins shared between both parasitoids, a gamma glutamyl transpeptidase (GGT1) is the most abundant protein in the venom of both *A. ervi* (Colinet *et al*. 2014) and *L. fabarum* (Additional File 7). This protein has been suggested to be involved in the castration of the aphid host after parasitism (Falabella *et al*. 2007). As previously reported for *A. ervi* (Colinet *et al*. 2014), a second GGT venom protein (GGT2) containing mutations in the active site was also found in the venom of *L. fabarum* (Supplementary Figure 16, 17). Phylogenetic analysis (Figure 5) revealed that the *A. ervi* and *L. fabarum* GGT venom proteins occur in a single clade in which GGT1 venom proteins group separately from GGT2 venom proteins, thus suggesting that they originated from a duplication that occurred prior to the split from their most recent common ancestor. As previously shown for *A. ervi*, the GGT venom proteins of *A. ervi* and *L. fabarum* are found in one of the three clades described for the non-venomous hymenopteran GGT proteins (clade “A”, Figure 5 and Colinet *et al*. 2014). Within this clade, venomous and non-venomous GGT proteins had a similar exon structure, except for exon 1 that corresponds to the signal peptide only present in venomous GGT proteins (Supplementary Figure 17). *Aphidius ervi* and *L. fabarum* venomous GGT proteins thus probably result from a single imperfect duplication of the non-venomous GGT gene belonging to clade A in their common ancestor, followed by recruitment of the signal peptide coding sequence. This first imperfect duplication event would then have been followed by a second duplication of the newly recruited venomous GGT gene before the separation of both species.

**Figure 4.**
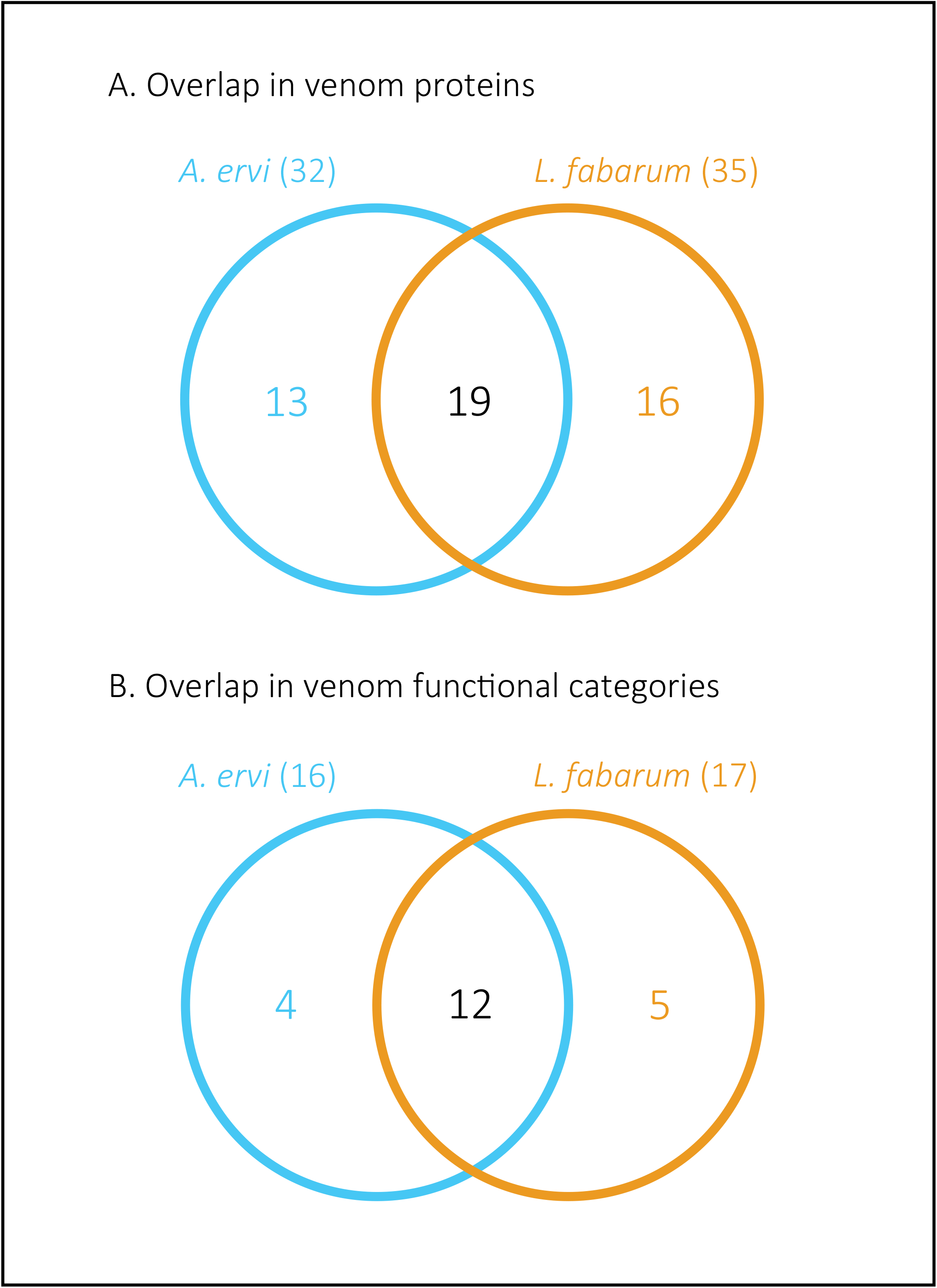
Overlap in Venom proteins between *A. ervi* and *L. fabarum*: Overlap in venom proteins (A) and venom protein putative function (B) between *A. ervi* and *L. fabarum*

**Figure 5:**
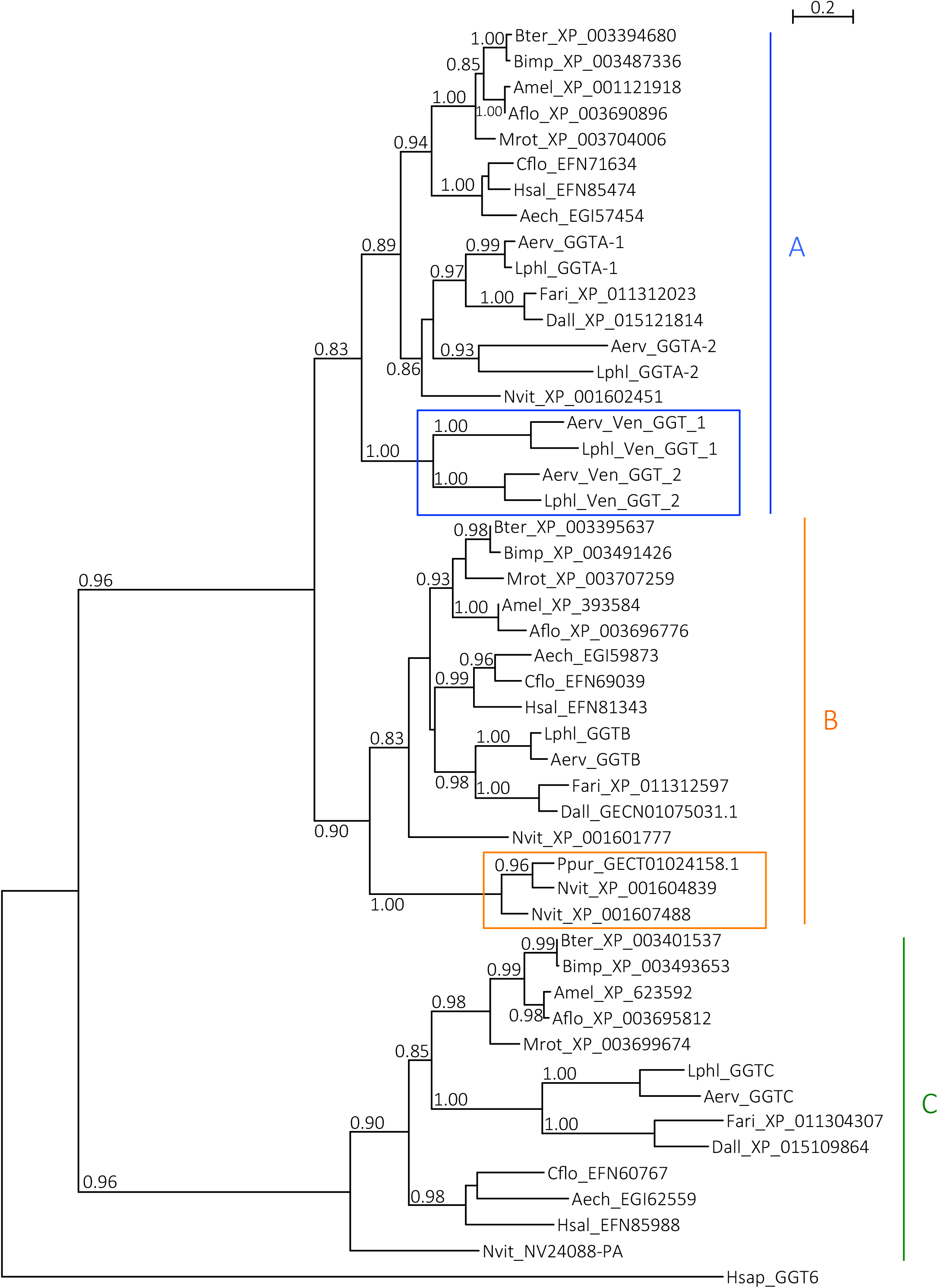
Phylogeny of hymenopteran GGT sequences. *A. ervi/L. fabarum* and N. vitripennis/P. puparum venom GGT sequences are marked with blue and orange rectangles respectively. Letters A, B and C indicate the major clades observed for hymenopteran GGT sequences. Numbers at corresponding nodes are aLRT values. Only aLRT support values greater than 0.8 are shown. The outgroup is human GGT6 sequence.

The presence of truncated *LRR* proteins was previously reported in venom of *A. ervi* (Colinet et al. 2014) and other Braconidae (Mathé-Hubert et al. 2016) that likely interfere with the host immune response. Several *LRR* proteins were found in the venom of *L. fabarum* as well, however these results should be interpreted with caution since the sequences were incomplete and the presence of a signal peptide could not be confirmed (Additional File 7). Moreover, these putative venom proteins were only identified from transcriptomic data of the venom apparatus and we could not find any corresponding annotated gene in the genome. This supports the idea that gene-family expansions in putative F-box/*LRR* proteins (discussed above) are not related to venom production.

Approximately 50% of the identified venom proteins were unique to either *A. ervi* or *L. fabarum,* and these could be related to their differing host ranges (Additional File 7). However, most of these proteins had no predicted function, making it difficult to hypothesize their possible role in parasitism success. Among the venom proteins with a predicted function, an apolipophorin was found in the venom of *L. fabarum* but not in *A. ervi*. Apolipohorin is an insect-specific apolipoprotein involved in lipid transport and innate immunity that is not commonly found in venoms. Among parasitoid wasps, apolipophorin has been described in the venom of the ichneumonid *Hyposoter didymator* (Dorémus *et al*. 2013) and the encyrtid *Diversinervus elegans* (Liu *et al*. 2017), but its function is yet to be deciphered. Apolipophorin is also present in low abundance in honeybee venom where it could have antibacterial activity (Kim & Jin 2015; Van Vaerenbergh *et al*. 2014). Lastly, we could not find *L. fabarum* homologs for any of the three secreted cystein-rich toxin-like peptides that are highly expressed in the *A. ervi* venom apparatus (Additional File 7). However, this may not be definitive since the search for similarities in the genome is complicated by the small size of these toxin-like sequences.

#### Key gene families

We manually annotated more than 1,000 genes (667 for *A. ervi* and 598 for *L. fabarum*; Table 3) using Apollo, hosted on the BIPAA website (Dunn *et al*. 2019; https://bipaa.genouest.org; Lee *et al*. 2013) to confirm and improve the results of the machine annotation. This is especially important for large gene families, which are usually poorly annotated by automatic prediction (Robertson *et al*. 2018); since such gene families potentially underlie key adaptive differences between the two parasitoids, accurate annotation is needed.

### Desaturases

Desaturases are an important gene family that introduce carbon-carbon double bonds in fatty acyl chains in insects (Los & Murata 1998; Sperling *et al*. 2003). While these function broadly across taxa, a subset of these genes (specifically acyl-CoA desaturases) have been implicated in insect chemical recognition for roles including alkene production and modification of fatty acids (Helmkampf *et al*. 2015). This gene family is particularly interesting because it has been shown that *Lysiphlebus cardui*, a close relative of *L. fabarum,* have no unsaturated cuticular hydrocarbons, just as is seen in its aphid host. This allows the parasitoid to go undetected in aphid colonies that are ant-tended and therefore better parasitize them (Liepert & Dettner 1996). We confirmed that the same is true for *L. fabarum*; its CHC profile is dominated by saturated hydrocarbons (alkanes), contains only trace alkenes, and is completely lacking dienes (Supplementary Figure 18, 20). In contrast, *A. ervi* females produce a large amount of unsaturated hydrocarbons, with a significant amount of alkenes and alkadiens in their CHC profiles (app. 70% of the CHC profile are alkenes/alkadienes, Supplementary Figure 19, 20).

The loss of one annotated desaturase gene in *L. fabarum* compared to *A. ervi* (Table 3) might explain these differences in the composition of their CHC profiles, especially their apparent inability to synthesize dienes. We also note there is little evidence that members of this gene family are clustered in the genome (just three and two desaturase genes in the same scaffolds of *A. ervi* and *L. fabarum*, respectively). Further investigations should verify this loss in *L. fabarum*, identify the ortholog of the missing copy in *A. ervi*, and test if this potential lost desaturase gene in *L. fabarum* is involved in the generation of unsaturated CHCs in *A. ervi*. This would determine if this loss is a key adaptation for mimicry of their aphid hosts’ cuticular hydrocarbon profiles in *L. fabarum*.

### Immune genes

We searched for immune genes in the two genomes based on a list of 367 immunity related genes, collected primarily from the *Drosophila* literature (Additional File 8). Using blast-based searches, 204 of these genes (59%) were found and annotated in both species. Six were present in only the *A. ervi* genome and six in only the *L. fabarum* genome. We compared these with the immune genes used to define the main *Drosophila* immune pathways (Toll, Imd, and JAK-STAT, Supplementary Table 10) and conserved in a large number of insect species (Buchon *et al*. 2014; Charroux & Royet 2010; Lemaitre & Hoffman 2007). Among these genes there are several well characterized pathways. The *D. melanogaster* Toll pathway is essential for the response to fungi and Gram-positive bacteria (Valanne *et al*. 2011). It was initially identified as a developmental pathway acting via the nuclear factor kappa B (NF-κB). The Imd/NF-kappa-B pathway is pivotal in the humoral and epithelial immune response to Gram-negative bacteria. Signaling through *imd* (a death domain protein) ultimately activates the transcription of specific antimicrobial peptides (AMPs, Myllymäki *et al*. 2014). The JAK-STAT pathway is involved in the humoral and cellular immune response (Morin-Poulard *et al*. 2013). It is activated after a cytokine-like protein called unpaired (*upd*) binds to its receptor Domeless (Dome). Activated JAK phosphorylates STAT molecules that translocate into the nucleus, where they bind the promoters of target genes.

In the genome of both wasps, many genes encoding proteins of the Imd and Toll pathways were absent, such as upstream GNBPs (Gram Negative Binding Proteins) and PGRPs (Peptidoglycan Recognition Proteins) and downstream AMPs (Supplementary Table 10, Supplementary Figure 21, Additional File 8). While none of these genes were found in *L. fabarum*, one PGRP related to PGRP-SD, involved in the response to Gram-positive bacteria (Bischoff *et al*. 2004), and one *defensin*-related gene were found in *A. ervi*. The *imd* gene was also absent in in both wasps; this is noteworthy because *imd* has been present in other hymenopteran genomes analyzed to date. Strikingly, all of the Imd pathway genes, including GNBP- and PGRP-encoding genes, *imd*, *FADD*, *Dredd* and *Relish* are lacking in aphid genomes (*A. pisum*, *A. gossypii* and *D. noxia*, via AphidBase (Legeai et al. 2010) and Gerardo et al (2010)), and *imd* is absent in *A. glycines*, *M. persicae, M. cerisae, R. padi* genomes, some of which are hosts for *A. ervi* and *L. fabarum* (Kavallieratos *et al*. 2004). The lack of an Imd pathway in aphids is suggested to be an adaptation to tolerate the obligate bacterial symbiont, *Buchnera aphidicola*, as well as their facultative endosymbionts that are gram-negative gamma-proteobacteria (e.g. *Hamiltonella defensa*). These facultative symbionts exhibit defensive activities against microbial pathogens and insect parasitoids (Guo *et al*. 2017; Leclair *et al*. 2016; Oliver *et al*. 2010; Scarborough *et al*. 2005) and may at least partially compensate for the host aphids innate immune functions. Recent data also suggest that cross-talk occurs between the Imd and Toll pathways to target wider and overlapping arrays of microbes (Nishide *et al*. 2019). Whether a similar cross-talk occurs in these two Aphidiidae (*A. ervi* and *L. fabarum*) needs further study.

Overall, our results suggest convergent evolution of loss in immunity genes, and possibly function, between these parasitoids and their aphid hosts. One reason might be that during the early stages of development, parasitoids need host symbionts to supply their basic nutrients, and thus an immune response from the parasitoid larvae might impair this function. Alternatively, but not exclusively, mounting an immune response against bacteria by the parasitoid larvae may be energetically costly and divert resources from its development. This idea of energy conservation would be especially relevant if the GC-loss discussed above is a mechanism to conserve resources. In both cases, the immune response will be costly for the parasitoid. Further work is needed to address whether other unrelated aphid parasitoids are lacking *imd*, upstream activators, and downstream effectors of the immune pathways (a preliminary blast search suggests that *imd* is present in the Aphelinidae *Aphelinus abdominalis*). This impaired immunity might lead to a decrease in both wasps’ responses to pathogenic bacteria, or they may use other defensive components to fight bacterial infections (perhaps some in common with aphids) that await to be discovered. For example, in *L. fabarum*, recent transcriptomic work has shown that detoxifying genes may be a key component of parasitoid success (Dennis *et al*. in revision), and these could play a role in immunity.

### Osiris genes

The Osiris genes are an insect-specific gene family that underwent multiple tandem duplications early in insect evolution. These genes are essential for proper embryogenesis (Smoyer *et al*. 2003) and pupation (Andrade López *et al*. 2017; Schmitt-Engel *et al*. 2015), and are also tied to immune and toxin-related responses (e.g. Andrade López *et al*. 2017; Greenwood *et al*. 2017) and developmental polyphenism (Smith *et al*. 2018; Vilcinskas & Vogel 2016).

We found 21 and 25 putative Osiris genes in the *A. ervi* and *L. fabarum* genomes, respectively (Supplementary Tables 11, 12). In insects with well assembled genomes, there is a consistent synteny of approximately 20 Osiris genes; this cluster usually occurs in a ∼150kbp stretch and gene synteny is conserved in all known Hymenoptera genomes (Supplementary Figure 22). The Osiris cluster is largely devoid of non-Osiris genes in most of the Hymenoptera, but the assemblies of *A. ervi* and *L. fabarum* suggest that if the cluster is actually syntenic in these species, there are interspersed non-Osiris genes (those are black boxes in Supplementary Figures 23 and 24).

In support of their role in defense (especially metabolism of xenobiotics and immunity), these genes were much more highly expressed in larvae than in adults (Supplementary Table 12). We hypothesize that their upregulation in larvae is an adaptive response to living within a host. Because of the available transcriptomic data, we could only make this comparison in *L. fabarum*. Here, 19 of the 26 annotated Osiris genes were significantly differentially expressed in larvae over adults (Supplementary Table 12, Additional File 9). In both species, transcription in adults was very low, with fewer than 10 raw reads per cDNA library sequenced, and most often less than one read per library.

### OXPHOS

In most eukaryotes, mitochondria provide the majority of cellular energy (in the form of adenosine triphosphate, ATP) through the oxidative phosphorylation (OXPHOS) pathway. OXPHOS genes are an essential component of energy production, and have increased in Hymenoptera relative to other insect orders (Li *et al*. 2017). We identified 69 out of 71 core OXPHOS genes in both genomes, and identified five putative duplication events that are apparently not assembly errors (Supplementary Table 13, Additional File 10). The gene sets of *A. ervi* and *L. fabarum* contained the same genes and the same genes were duplicated in each, implying duplication events occurred prior to the split from their most recent common ancestor. One of these duplicated genes appears to be duplicated again in *A. ervi*, or the other copy has been lost in *L. fabarum*.

### Chemosensory genes

Genes underlying chemosensory reception play important roles in parasitoid mate and host localization (Comeault *et al*. 2017; Nouhaud *et al*. 2018). Several classes of chemosensory genes were annotated separately (Table 4): odorant receptors (ORs) are known to detect volatile molecules, odorant-binding proteins (OBPs) and chemosensory proteins (CSPs) are possible carriers of chemical molecules to sensory neurons, and ionotropic receptors (IRs) are involved in both odorant and gustatory molecule reception. With these manual annotations, further studies can now be made with respect to life history characters including reproductive mode, specialization on aphid hosts, and mimicry.

**Table 4:** Summary of annotation of putative DNA methylation genes.

| Species | Gene | Scaffold | e-value (*Nasonia*) |
| --- | --- | --- | --- |
| *A. ervi* | EEF1AKMT1 homolog | scaffold94 | 1.00E-66 |
| *L. fabarum* |  | tig00000449 | 5.00E-63 |
| *A. ervi* | DNA methyltransferase 3 | scaffold45 | 5.00E-138 |
| *L. fabarum* |  | tig00002022 | 9.00E-117 |
| *A. ervi* | DNA methyltransferase 1 | *no homolog detected* |  |
| *L. fabarum* |  | *no homolog detected* |  |

#### Chemosensory: Soluble proteins (OBPs and CSPs)

Hymenoptera have a wide range of known OBP genes, with up to 90 in *N. vitripenis* (Vieira *et al*. 2012). However, the numbers of these genes appear to be similar across parasitic wasps, with 14 in both species studied here and 15 recently described in *D. alloeum* (Tvedte *et al*. 2019). Similarly, CSP numbers are in the same range within parasitic wasps (11 and 13 copies here, Table 4). Interestingly, two CSP sequences (one in *A. ervi* and one in *L. fabarum*) did not have the conserved cysteine motif, characteristic of this gene family. So although they were annotated here, further work should investigate if and how these genes function.

#### Chemosensory: Odorant receptors (ORs)

In total, we annotated 228 putative ORs in *A. ervi* and 156 in *L. fabarum* (Table 4). This is within the range of OR numbers annotated in other hymenopteran parasitoids, including: 79 in *M. cingulum* (Ahmed *et al*. 2016), 225 in *N. vitripennis* (Robertson *et al*. 2010), and 187 in *D. alloeum* (Tvedte *et al*. 2019). Interestingly, we annotated a larger set of ORs in *A. ervi* than in *L. fabarum*. One explanation is that *A. ervi* generally has more annotated genes than *L. fabarum*, and whatever broad pattern underlies the reduction in the gene repertoire of *L. fabarum* also affected OR genes. One functional explanations for a lower number of OR genes in *L. fabarum* is that the *A. ervi* strain sequenced of was derived from several field strains that parasitized different hosts on different host plants, and the ability to parasitize a broader host range could select for more OR genes (Monticelli *et al*. 2019).

#### Chemosensory: Ionotropic chemosensory receptors (IRs)

In total, we annotated 38 putative IRs in *A. ervi* and 37 in *L. fabarum* (Table 4). Three putative co-receptors (IR 8a, IR 25a and IR 76b) were annotated both species, one of which (IR 76b) was duplicated in *A. ervi*. This bring the total for the IR functional group to 42 and 40 genes for *A. ervi* and *L. fabarum*, respectively. This is within the range of IRs known from other parasitoid wasps such as *Aphidius gifuensis* (23 IRs identified in antennal transcriptome, Braconidae, Kang *et al*. 2017), *D. alloeum* (51 IRs, Braconidae, Tvedte *et al*. 2019) and *N. vitripennis* (47 IRs, Pteromalidae, Robertson *et al*. 2010). A phylogenetic analysis of these genes showed a deeply rooted expansion in the IR genes (Supplementary Figure 25). Thus, in contrast to the expansion usually observed in hymenopteran ORs compared to other insect orders, IRs have not undergone major expansions in parasitic wasps, which is generally the case for a majority of insects with the exception of Blattodea (Harrison *et al*. 2018)

### Sex determination

The core sex determination genes (*transformer*, *doublesex*) are conserved in both species (Supplementary Table 14, Additional File 11). Notably, *A. ervi* possesses a putative *transformer* duplication. This scaffold carrying the duplication (scaffold2824) is only fragmentary, but a *transformer* duplicate has also been detected in the transcriptome of a member of the *A. colemani* species complex, suggesting a conserved presence within the genus (Peters *et al*. 2017). In *A. ervi*, *transformer* appears to have an internal repeat of the CAM-domain, as is seen in the genus *Asobara* (Geuverink *et al*. 2018). In contrast, there is no evidence of duplication in sex determination genes in *L. fabarum*. This supports the idea that complementary sex determination (CSD) in sexually reproducing *L. fabarum* populations is based on up-stream cues that differ from those known in other CSD species (Matthey-Doret *et al*. 2019), whereas the CSD locus known from other hymenopterans locus is a paralog of transformer (Heimpel & de Boer 2007).

In addition to the core sex determination genes, we identified homologs of several genes related to sex determination (Supplementary Table 15). We identified *fruitless* in both genomes, which is associated with sex-specific behavior in taxa including *Drosophila* (Yamamoto 2008). Both genomes also have homologs of *sex-lethal* which is the main determinant of sex in *Drosophila* (Bell *et al*. 1988). *Drosophila* has two homologs of this gene, and the single version in Hymenoptera may have more in common with the non-sex-lethal copy, called *sister-of-sex-lethal*. We identified homologs of the gene *CWC22*, including a duplication in *A. ervi*; this duplication is interesting because a duplicated copy of *CWC22* is the primary signal of sex determination in the house fly *Musca domestica* (Sharma *et al*. 2017). Lastly, there was a duplication of *RBP1* in both genomes. The duplication of *RBP1* is not restricted to these species, nor is the duplications of *CWC22*, which appears sporadically in Braconidae. Together, these annotations add to our growing knowledge of duplications of these genes, and provide possibilities for further examinations of the role of duplications and specialization in association with sex determination.

### DNA Methylation genes

DNA methyltransferase genes are thought to be responsible for the generation and maintenance of DNA methylation. In general, DNA methyltransferase 3 (*DNMT3*) introduces *de novo* DNA methylation sites and DNA methyltransferase 1 (*DNMT1*) maintains and is essential for DNA methylation (Jeltsch & Jurkowska 2014; Provataris *et al*. 2018). A third gene, *EEF1AKMT1* (formerly known as *DNMT2*), was once thought to act to methylate DNA but is now understood to methylate tRNA (Provataris *et al*. 2018). In both *A. ervi* and *L. fabarum*, we successfully identified homologs *DNMT3* and *EEF1AKMT1*. In contrast, *DNMT1* was not detected in either species (Table 4, Supplementary Table 16). This adds to growing evidence that these genes are not conserved across family Braconidae, as *DNMT1* appears to be absent in several other braconid genera, including *Asobara tabida*, *A. japonica*, *Cotesia sp*., and *F. arisanus* (Bewick *et al*. 2017; Geuverink 2017). However, *DNMT1* is present in some braconids, including *M. demolitor*, and outside of Braconidae these genes are otherwise strongly conserved across insects. In contrast, DNMT3, present here, is more often lost in insects (Provataris *et al*. 2018).

This absence of *DNMT1* helps explains previous estimates of very low DNA methylation in *A. ervi* (0.5%, Bewick *et al*. 2017). We confirmed these low levels of methylation in *A. ervi* by mapping this previously generated bisulfite sequencing data (Bewick *et al*. 2017) to our genome assembly. We aligned >80% of their data (total 94.5Mbp, 625,765 reads). The sequence coverage of this mapped data was low: only 63,554 methylation-available cytosines were covered and only 1,216 were represented by two or more mapped reads. Nonetheless, of these mapped cytosines, the vast majority (63,409) were never methylated, just 143 sites were always methylated, and two were variably methylated. Methylation-available cytosine classes were roughly equally distributed among three cytosine classes (CG: 0.154%, CHG: 0,179%, and CHH: 0.201%). This methylation rate is less than the 0.5% estimated by Bewick (2017) and confirms a near absence of DNA methylation in *A. ervi*. Given the parallel absence of DNMT1 in *L. fabarum*, it seems likely that both species sequenced here may have very low levels of DNA methylation, and that this is not a significant mechanism in these species.

This stark reduction in DNA methylation is interesting, given that epigenetic mechanisms are likely important to insect defenses, including possible responses to host endosymbionts (Huang *et al*. 2019; Vilcinskas 2016, 2017). As with the immune pathways discussed above, this could reflect a loss that is adaptive to developing within endosymbiont-protected hosts. It is also interesting that while one epigenetic mechanism seems to be absent in both *A. ervi* and *L. fabarum*, we see an increase in histone variants in *L. fabarum* (based on the OMA analysis of gene family expansion), and these histones could function in gene regulation. However, whether there is a functional or causal link between these two observations is yet to be tested.

## Conclusions

These two genomes have provided insight into adaptive evolution in parasitoids that infect aphids. Both genomes are extremely GC-poor, and their extreme codon bias provides an excellent system for examining the chemical biases and selective forces that may overshadow molecular evolution in eukaryotes. We have also highlighted several groups of genes that are key to functional evolution across insects, including venom, sex determination, response to bacterial infection (F-box/*LRR* proteins), and near absence of DNA methylation. Moreover, the absence of certain immune genes (e.g. from the Imd and Toll pathways) in these two species is similar to losses in host aphids, and raises intriguing questions related to the effects of aphids’ symbiosis on both aphid and parasitoid genomics.

Parasitoid wasps provide an excellent model for studying applied and basic biological questions, including host range (specialist vs generalist), reproductive mode (sexual vs asexual), antagonistic coevolution, genome evolution, and epigenetic regulation, to mention just a few. Our new genomic resources will open the way for a broad set of future research, including work to understand host specialization, adaptive changes associated with climate, and the potential loss of diapause in *A. ervi* (Tougeron *et al*. 2019; Tougeron *et al*. 2017). Lastly, the genomes of these two non-social Hymenoptera provide a valuable comparison for understanding processes specific to social insects with complex caste structure, and are a first but essential step to better understand the genetic architecture and evolution of traits that are important for a parasitic life style and their use in biological control.

## Methods

*\*More complete methods are available in the Supplementary Material*

### Insect collection and origin

#### Aphidius ervi

*Aphidius ervi* samples used for whole-genome sequencing came from two different, sexually reproducing, isofemale lines established from parasitized aphids (recognizable as mummies) from fields of cereals and legumes in two different geographic zones in Chile: Region de Los Rios (S 39° 51’, W 73° 7’) and Region del Maule (S 35° 24’, W 71° 40’). Mummies (parasitized aphids) of *Sitobion avenae* aphids were sampled on wheat (*Triticum aestivum* L.) while mummies of *Acyrtosiphon pisum* aphids were sampled on *Pisum sativum* L. (pea aphid race). Aphid mummies were isolated in petri dishes until adult parasitoids emerged. These two parasitoid lineages were separated in two cages with hosts *ad libitum* and were propagated for approximately 75 generations under controlled conditions as described elsewhere (Ballesteros *et al*. 2017; Sepúlveda *et al*. 2016). A further reduction of genetic variation was accomplished by establishing two isofemale *A. ervi* lines, which were maintained as described previously and propagated for approximately 10 generations before adult parasitoids (male and female) were collected live and stored in 1.5 ml centrifuge tubes containing ethanol (95%) at −20°C.

*Aphidius ervi* samples used for CHC analysis (below) were purchased from Katz Biotech AG (Baruth, Germany). Species identification was confirmed with COI barcoding following Hebert *et al*. (2003). Wasps sacrificed for CHC analysis were sampled from the first generation reared in the lab on *Acyrtosiphon pisum* strain LL01 (Peccoud *et al*. 2009), which were mass-reared on *Vicia faba* cv. *Dreifach Weisse*.

#### Lysiphlebus fabarum

*Lysiphlebus fabarum* samples used for whole-genome sequencing came from a single, asexually reproducing, isofemale line (IL-07-64). This lineage was first collected in September 2007 from Wildberg, Zürich, Switzerland as mummies of the aphid *Aphis fabae fabae*, collected from the host plant *Chenopodium album*. In the lab, parasitoids were reared on *A. f. fabae* raised on broad bean plants (*Vicia faba*) under controlled conditions [16 h light: 8 h dark, 20°C] until sampling in September 2013, or approximately 150 generations. Every lab generation was founded by ca. 10 individuals that were transferred to fresh host plants containing wasp-naïve aphids. Approximately 700 individuals were collected for whole-genome sequencing from a single generation in December 2013 and flash frozen at −80°C. To avoid sequencing non-wasp DNA, samples were sorted over dry ice to remove any contaminating host aphid or plant material.

For linkage group construction, separate *L. fabarum* collections were made from a sexually reproducing lineage. Here, we collected all sons produced by a single virgin female, sampled from the control lineage in a recently employed evolution experiment (H-lineage; Dennis *et al*. 2017). Wasps were stored on ethanol until RAD-seq library construction. Lastly, a third population was sampled for the proteomic analysis of the venom-apparatus (below); these females came from the genetically-diverse starting population used to found the evolution experiment of Dennis *et al*. (2017), and were sampled in December 2014.

### DNA extraction and library preparation

#### Aphidius ervi

DNA was extracted from adult haploid males of *A. ervi* in seven sub-samples (ca. 120 males each), reared in *S. avenae*. Total DNA was extracted using the DNEasy Plant Mini Kit (QIAGEN) following the manufacturer’s instructions. DNA was quantified by spectrophotometry (Epoch Microplate Spectrophotometer, Biotek) and fluorometry (Qubit 3.0; Qubit DNA High sensitivity Assay Kit, Invitrogen), and quality was assessed using 1% agarose gel electrophoresis. DNA samples were sent on dry ice to MACROGEN (Seoul, South Korea) and were used to produce Illumina paired-end (PE) and mate-pair (MP) libraries for sequencing. A PE library was constructed from one of the seven sub-samples (120 individuals, 1μg DNA) sheared by ultrasonication (Covaris) company, average sheared insert size: 350bp). The remaining DNA samples were pooled (6 samples, 720 individuals) and used for MP sequencing (3kb, 5kb and 8kb insert sizes), which were prepared with the Nextera mate-pair protocol (Illumina). All libraries were sequenced using an Illumina HiSeq 2000 sequencer (MACROGEN).

Long read PacBio (Pacific Biosciences) RS II sequencing was performed from a single DNA extraction of 270 *A. ervi* females, reared on *A. pisum*. Genomic DNA was extracted using the Wizard genomic DNA purification kit (Promega) according to manufacturer instructions and quantified spectrophotometrically using a NanoDrop 2000 (Thermo Scientific). Input DNA was mechanically sheared to an average size distribution of 10Kb (Covaris gTube, Kbiosciences) and the resulting library was size selected on a Blue Pippin Size Selection System (Cat #BLU0001, Sage Science) to enrich fragments > 8Kb. Quality and quantity were checked on Bioanalyzer (Agilent Technologies) and Qubit, respectively. Four SMRT RSII cells with P6 chemistry were sequenced at GenoScreen, France.

#### Lysiphlebus fabarum

DNA was extracted from adult female *L. fabarum* in 10 sub-samples (50-100 wasps each) using the QIAmp DNA mini Kit (Qiagen) according to the manufacturer’s instructions, with the inclusion of an overnight tissue digestion at 56°C. Extracted DNA was then pooled and used to produce Illumina PE and MP, and PacBio libraries. The PE library was prepared using the Illumina Paired-End DNA protocol; the average fragment size was 180 base pair (bp). The MP library (5kb insert) was generated with the Nextera mate-pair protocol (Illumina). Both libraries were sequenced on the Illumina MiSeq in Paired-End mode at the University of Zürich.

Long-read libraries for PacBio RS II sequencing were produced using the DNA Template Prep Kit 2.0 (Pacific Biosciences). Input DNA was mechanically sheared to an average size distribution of 10Kb (Covaris gTube, Kbiosciences) and the resulting library was size selected on a Blue Pippin Size Selection System (Sage Science) machine to enrich fragments > 8Kb; quality and quantity were checked on the Bioanalyzer and Qubit, respectively. Ten SMRT Cells were sequenced at the University of Zürich.

### Genome assembly

#### Aphidius ervi

Library quality was checked with FastQC ver. 0.11.3 (Andrews *et al*. 2010). Paired-end libraries were processed with Trimmomatic ver. 0.35 (Bolger et al., 2014) for trimming Illumina adapters/primers, low quality bases (Q <25, 4bp window) and discarding sequences shorter than 50bp or without its mate-pair. In the case of Mate-Pair libraries, removal of improperly oriented read-pairs and removal of Nextera adapters was performed using NextClip (Leggett *et al*. 2014). Filtered PE and MP libraries were used for genome assembly with Platanus ver. 1.2.1 with default parameters (Kajitani *et al*. 2014), gap closing was performed with GapCloser (Luo *et al*. 2012). Scaffolding with PacBio reads was performed using a modified version of SSPACE-LR v1.1 (Boetzer & Pirovano 2014), with the maximum link option set by –a 250. Finally, the gaps of this last version were filled with the Illumina reads using GapCloser.

#### Lysiphlebus fabarum

Library quality was also checked with FastQC (Andrews *et al*. 2010). Illumina reads were filtered using Trimmomatic to remove low quality sequences (Q<25, 4bp window), to trim all Illumina primers, and to discard any sequence shorter than 50bp or without its mate-pair. NextClip was used to remove all improperly oriented read pairs.

Raw PacBio reads were error-corrected using the quality filtered Illumina data with the program Proovread (Hackl *et al*. 2014). These error-corrected reads were then used for *de novo* assembly in the program *canu* v1.0 (Koren et al. 2017). Since our PacBio reads were expected to have approximately 30X coverage (based on the presumed size of 128MB), *Canu* was run with the recommended settings for low coverage data (corMhapSensitivity=high corMinCoverage=2 errorRate=0.035), and with the specification that the genome is approximately 128Mbp. The resulting assembly was polished using Pilon (Walker *et al*. 2014) to correct for both single nucleotide and small indel errors, using mapping of both the MP and PE data, generated with bwa-mem (Li & Durbin 2009).

### Linkage map construction: *L. fabarum*

For linkage map construction, we followed the methodology described in Wang *et al*. (2013) and Purcell *et al*. (2014). In brief, we genotyped 124 haploid male offspring from one sexual female using ddRADseq. Whole-body DNA was high-salt extracted (Aljanabi & Martinez 1997), digested with the *EcoRI* and *MseI* restriction enzymes, and ligated with individual barcodes (Parchman *et al*. 2012; Peterson *et al*. 2012). Barcoded samples were purified and amplified with Illumina indexed primers by PCR (Peterson *et al*. 2012) and quality-checked on an agarose gel.

Pooled samples were sequenced on the Illumina HiSeq2500. Raw single-end libraries were quality filtered and de-multiplexed using the process_radtags routine within Stacks v1.28 with default parameters (Catchen *et al*. 2011), and further filtered for possible adapter contamination using custom scripts. Genotyping was performed by mapping all samples against the *L. fabarum* draft genome assembly using bowtie2 (Langmead & Salzberg 2012) with rg-id, sensitive and end-to-end options. Genotypes were extracted using samtools mpileup (Li *et al*. 2009) and bcftools (haploid option, Li 2011). We filtered the resulting genotypes for a quality score >20 and removed loci with >20% missing data and/or a minor allele frequency <15% using VCFtools v0.1.12b (Danecek *et al*. 2011). After filtering, 1,319 biallelic SNPs in 90 offspring remained.

For constructing linkage groups, we followed Gadau (2009) to account for the unknown phase of the maternal genotype. In short, we duplicated the haploid male genotypes and reversed the phase for one duplicated set and removed one of the mirror linkage group sets after mapping. We generated the map using MSTmap (Wu *et al*. 2008) on the data with following parameters: population_type DH, distance_function kosambi, no_map_dist 15.0, no_map_size 2, missing_threshold 1.00, and the cut_off_p_value 1e-6. The cut-off p-value was adjusted to create a linkage map of five linkage groups, however the biggest group had a gap of >70 cM, indicating a false fusion of two groups, which we split in two groups. This result corresponded to the six chromosomes previously described for *L. fabarum* (Belshaw & Quicke 2003), these were visualized with AllMaps (Tang *et al*. 2015). Initial mapping showed that 14 SNPs at one end of tig0000000 mapped to Chromosome1, while the majority of the contig (>150,000 bp) mapped to Chromosome 2. Thus, these SNPs were removed from the linkage maps, and it is advised that subsequent drafts of the *L. fabarum* genome should split this contig around position 153,900.

### Genome completeness and synteny

Completeness of the two assemblies was assessed by identifying Benchmarking Universal Single-Copy Orthologs (BUSCOs) using the BUSCO v3.0.2 pipeline in genome mode (Simão *et al*. 2015). We identified single copy orthologs based on the Arthropoda_db9 (1,066 genes, training species: *Nasonia vitripennis*).

Synteny between the two genomes was assessed using the NUCmer aligner, which is part of the MUMmer v3.23 package (Kurtz *et al*. 2004). For this, we used the *L. fabarum* chromosomes as the reference, and included the scaffolds not incorporated into chromosomes (total 1,407 pieces). The *A. ervi* assembly was mapped to this using the default settings of NUCmer.

### Predictive gene annotation

For both assembled genomes, gene predictions were generated using MAKER2 (Holt & Yandell 2011). Within MAKER2, predictive training was performed in a three step process. A first set of genes was predicted by similarity to known proteins or contigs from RNAseq in the same species (described below). This gene set was used thereafter for training both Augustus (Keller *et al*. 2011) and SNAP (Korf 2004), in two steps, with the results of the first training re-used to train the software in the second round. Transcriptomic evidence was provided separately for each species. For *A. ervi*, six separate *de novo* transcriptome assemblies from Trinity (Grabherr *et al*. 2011) were constructed, one each for the adults reared on different hosts (NCBI PRJNA377544, Ballesteros *et al*. 2017). For each transcript, we only included variants based on filtering with RSEM v 1.2.21 using the option –fpkm_cutoff 1.0, --isopct_cutoff=15.00. This resulted in 452,783 transcripts. For *L. fabarum*, we utilized a joint transcriptome, built using RNAseq data (NCBI PRJNA290156) collected from adults (Dennis *et al*. 2017) and 4-5 day old larvae (Dennis *et al*. in review). Peptide evidence came from the Hymenoptera genomes database (http://hymenopteragenome.org, *Acromyrmex echiniator* v3.8, *Apis mellifera* v3.2, *Nasonia vitripennis* v1.2), from the BioInformatics Platform of Agroecosystems Arthropod database (https://bipaa.genouest.org, *Hyposoter didymator* v1.0), and *Drosphila melanogaster* (http://flybase.org, v6.13), and SwissProt (October 2016) databases. Summary statistics were generated with GAG (Hall *et al*. 2014). Transcriptomic support for the predicted genes was estimated by mapping available transcriptomic data (same as above) to the respective genomes using STAR (Dobin *et al*. 2013) in the “quantMode”.

### Functional annotation

The putative functions of the proteins predicted by the above pipeline were identified based on blastp (v2.5.0) matches against Genbank *nr* (non-redundant GenBank CDS translations+PDB+SwissProt+PIR+PRF) release 12/2016 and interproscan v5 against Interpro (1.21.2017). GO terms associations were collected from blast *nr* and interproscan results with blast2GO (v2.2). Finally, transmembrane domains were identified with Hidden Markov Models (HMM) in tmhmm v2.0c, and peptide signals with signalP (euk v4.1, Emanuelsson *et al*. 2007; Nielsen 2017).

### Transposable elements

Transposable elements (TE) were predicted using the REPET pipeline (Flutre *et al*. 2011), combining *de novo* and homology-based annotations. *De novo* prediction of TEs was restricted to scaffolds larger than the scaffold N50 for each species. Within these, repetitive elements were identified using a blast-based alignment of each genome to itself followed by clustering with Recon (Bao & Eddy 2002), Grouper (Quesneville *et al*. 2005) and Piler (Edgar & Myers 2005). For each cluster, a consensus sequence was generated by multiple alignment of all clustered elements with MAP (Huang 1994). The resulting consensus was then scanned for conserved structural features or homology to nucleotide and amino acid sequences from known TEs (RepBase 20.05, Bao *et al*. 2015; Jurka 1998) using BLASTER (tblastx, blastx, Flutre *et al*. 2011) or HMM profiles of repetitive elements (Pfam database 27.0) using hmmer3 (Mistry *et al*. 2013). Based on identified features, repeats were classified using Wicker’s TE classification as implemented in the PASTEclassifier (Hoede *et al*. 2014). The resulting *de novo* TE library for the genome was then filtered to retain only the elements with at least one perfect match in the genome. Subsequently, all TEs in the genomes were annotated with REPET’s TE annotation pipeline. Reference TE sequences were aligned to the genome using BLASTER, Repeat Masker (Smit *et al*. 2013-2015) and CENSOR (Kohany *et al*. 2006). The resulting HSPs were filtered using an empirical statistical filter implemented in REPET (Flutre *et al*. 2011) and combined using MATCHER (Quesneville *et al*. 2005). Short repeats were identified using TRF (Benson 1999) and Mreps (Kolpakov *et al*. 2003). Elements in genomic sequences with homology with known repbase elements (RepBase 20.05) were identified with BLASTER (blastx, tblastx) and curated by MATCHER. Finally, redundant TEs and spurious SSR annotations were filtered and separate annotations for the same TE locus were combined using REPET’s “long join procedure”.

### GC content and codon usage

We examined several measures of nucleotide composition, at both the nucleotide and protein level. Whole genome GC content was calculated by totaling the numbers of A, C, T, and G in the entire assembly. In the predicted coding sequences, this was also calculated separately for each predicted gene and third position GC composition was calculated separately in the predicted coding sequences. In all cases, this was done with the sscu package in R (Sun 2016). Relative Synonymous Codon Usage (RSCU) was extracted from the entire CDS using the seqinR package in R (Charif & Lobry 2007), and visualized with a PCA (R packages factoextra, reshape, and ggplot2, Kassambara & Mundt 2016; Wickham 2007, 2009). To examine GC content in coding genes of other insects, we downloaded the 118 available CDS in the RefSeq database of NCBI (date: October 2018) and again calculated per-gene GC content.

To examine the GC content of life-stage biased transcripts, we compared GC content in the genes that are significantly (FDR < 0.05) differentially expressed between previously generated transcriptomes from adult (Dennis *et al*. 2017) and larval (Dennis *et al*. in revision) *L. fabarum*, as well in the 10% most highly expressed genes in adults and larvae.

### Orphan genes

We identified orphan genes as those for which we could not find orthologs in any other sequenced genomes. To do this, we first used OrthoFinder (Emms & Kelly 2015) to generate clusters of orthologous and paralogous genes among the predicted genes (CDS) from the genomes of *A. ervi* and *L. fabarum*, as well as five other sequenced parasitoids (*Diachasma alloeum*, *Fopius arisanus*, *Macrocentrus cingulum*, *Microplitis demolitor* and *Nasonia vitripennis*). OrthoFinder produces a set of genes that were not assigned to any orthogroup. We identified species specific genes, which we are calling orphan genes, by removing all genes that had hits to any other genes in the *nt*, *nr*, and *swissprot* NCBI database (June 2019). Within these putative orphans, we only retained those with transcriptomic support.

### Gene family expansions

We examined gene families that have expanded and contracted in *A. ervi* and *L. fabarum* relative to one another using the OMA standalone package (v2.2.0, default values, Altenhoff *et al*. 2018). OMA was used to compute orthologs (OMA groups) and Hierarchical Orthologous Groups (HOGs) for the predicted proteins of *L. fabarum* and *A. ervi*: 15,203 and 20,344, respectively. While OMA groups consist of strict 1:1 orthologs between OGS1 and OGS3, HOGs contain all orthologs and paralogs of a given predicted gene family. HOGs were parsed with a custom Perl script to identify all gene families in which one of the wasp species contained more members than the other. We focused on only the groups that contained more than 20 genes (ten groups, Supplementary Figure 12). These were identified by blastx against the *nr* database in NCBI.

### Venom proteins

The *L. fabarum* venom proteomic analysis was performed from 10 extracted venom glands (Supplementary Figure 14). The 16 most visible bands in 1D gel electrophoresis were cut, digested with trypsin and analyzed by mass spectrometry. All raw data files generated by mass spectrometry were processed to generate mgf files and searched against: (i) the *L. fabarum* proteome predicted from the genome (*L. fabarum* annotation v1.0 proteins) and (ii) the *L. fabarum de novo* transcriptome (Dennis *et al*. 2017) using the MASCOT software v2.3 (Perkins *et al*. 1999). The mass spectrometry proteomics data have been deposited to the ProteomeXchange Consortium (http://proteomecentral.proteomexchange.org) via the PRIDE partner repository (Hanrahan & Johnston 2011), with the ID PXD015758.

Sequence annotation was performed based on blast similarity searches. Signal peptide prediction was performed with SignalP (Emanuelsson *et al*. 2007; Nielsen 2017). Searches for protein domains was performed with PfamScan (Finn *et al*. 2013) and venom protein genes were identified using the blast tools in Apollo (Dunn *et al*. 2019; Lee *et al*. 2013). Multiple amino acid sequence alignments were made with MUSCLE (Edgar 2004a, b). Phylogenetic analysis was performed using maximum likelihood (ML) with PhyML 3.0 (Guindon *et al*. 2010). SMS was used to select the best-fit model of amino acid substitution for ML phylogeny (Lefort *et al*. 2017).

### Manual gene curation

The two genome assemblies were manually curated for a number of gene families of interest. This improved their structural and functional annotation for more in-depth analysis. Manual curation, performed in Apollo included the inspection of stop/start codons, duplications (both true and erroneous), transcriptomic support, and concordance with the predicted gene models.

#### Desaturases

Desaturase genes in both genomes were automatically identified and annotated with GeMoMa (Keilwagen *et al*. 2016) using desaturase gene annotations from *Diachasma alloeum*, *Fopius arisanus*, and *Microplitis demolitor*, retrieved from NCBI’s protein database as queries (retrieved May 2017). Additionally, all desaturase genes were manually inspected.

To measure the production of desaturases in *A. ervi*, wasps were freeze-killed and stored separately by sex at - 20 ℃. For CHC extraction, single individuals were covered with 50 μl of MS pure hexane (UniSolv) in 2 ml GC vials (Agilent Technologies,) and swirled for 10 minutes on a Thermo-shaker (IKA KS 130 Basic, Staufen). The hexane extracts where then transferred to a fresh conical 250 μl GC insert (Agilent Technologies), where the hexane was completely evaporated under a constant flow of CO_2_. The dried extract was then resuspended in 5 μl of a hexane solution containing 7.5 ng/μl of n-dodecane (EMD Millipore Corp.) as an internal standard. 3 μl of the extract were then injected into a GC-QQQ Triple Quad (GC: 7890B, Triple Quad: 7010B, Agilent) with a PAL Autosampler system operating in electron impact ionization mode. The split/splitless injector was operated at 300 °C in Pulsed splitless mode at 20 psi until 0.75 min with the Purge Flow to Split Vent set at 50 mL/min at 0.9 min. Separation of compounds was performed on a 30 m x 0.25 mm ID x 0.25 μm HP-1 Dimethylpolysiloxane column (Agilent) with a temperature program starting from 60 °C, held for 2 min, and increasing by 50 °C per min to 200 °C, held for 1 min, followed by an increase of 8 °C per min to 250 °C, held again for 1 min, and finally 4 °C per min to 320 °C, held for 10 min. Post Run was set to 325 °C for 5 min. Helium served as carrier gas with a constant flow of 1.2 ml per min and a pressure of 10.42 psi. Initially CHC peaks were identified and the chromatogram was generated using the Qualitative Analysis Navigator of the MassHunter Workstation Software (vB.08.00 / Build 8.0.8208.0, Agilent). CHC quantification was performed using the Quantitative Analysis MassHunter Workstation Software (vB.09.00 / Build 9.0.647.0, Agilent). Peaks were quantified using their diagnostic (or the neighboring most abundant) ion as quantifier and several characteristic ions in their mass spectra as qualifiers to allow for unambiguous detection by the quantification software. The pre-defined integrator Agile 2 was used for the peak integration algorithm to allow for maximum flexibility. All peaks were then additionally checked for correct integration and quantification, and, where necessary, re-integrated manually. Percentages were based on the respective averages of four individual female CHC extracts.

### Immune genes

The list of immune genes to be searched against the *A. ervi* and *L. fabarum* genomes was established based on *Drosophila melanogaster* lists from the Lemaitre laboratory (lemaitrelab.epfl.ch/fr/ressources, adapted from De Gregorio *et al*. 2001; De Gregorio *et al*. 2002) and from the interactive fly web site (www.sdbonline.org/sites/fly/aignfam/immune.htm and Buchon *et al*. 2014). Each *D. melanogaster* protein sequence was used in blast similarity searches against the two predicted wasp proteomes. The best match was retained, and its protein sequence was used to perform a new blast search using the NCBI non-redundant protein sequence database to confirm the similarity with the *D. melanogaster* sequence. When both results were concordant, the retained sequence was then searched for in *Nasonia vitripennis* and *Apis mellifera* proteomes to identify homologous genes in these species.

### Osiris genes

Osiris gene orthologs were determined with a two-part approach: candidate gene categorization followed by phylogenetic clustering. Candidate Osiris genes were generated using HMM (with hmmer v3.1b2, Wheeler & Eddy 2013) and local alignment searching (blast, Altschul *et al*. 1990). A custom HMM was derived using all 24 well annotated and curated Osiris genes of *Drosophila melanogaster*. Next, an HMM search was performed on the *A. ervi* and *L. fabarum* proteomes, extracting all protein models with P < 0.05. Similarly, all *D. melanogaster* Osiris orthologs were searched in the annotated proteomes of *A. ervi* and *L. fabarum* using protein BLAST (e < 0.05). The top BLAST hit for each ortholog was then searched within each parasitoid genome for additional paralogs (e < 0.001). All unique candidates from the above approaches were then aligned using MAFFT (Katoh & Standley 2013), and an approximate maximum-likelihood phylogeny was constructed using FastTree (Price *et al*. 2009) via the CIPRES science gateway of Xsede (Miller *et al*. 2015). The species used were: the fruit fly (*D. melanogaste*r), the tobacco hornworm moth (*Manduca sexta*), the silkworm moth (*Bombyx mori*), the flour beetle (*Tribolium castaneum*), the jewel wasp (*Nasonia vitripennis*), the honeybee (*Apis mellifera*), the buff tail bumble bee (*Bombus terrestris*), the red harvester ant (*Pogonomyrmex barbatus*), the Florida carpenter ant (*Camponotus floridanus*), and Jerdon’s jumping ant (*Harpegnathos saltator*).

### OXPHOS

Genes involved in the oxidative phosphorylation pathway (OXPHOS) were identified in several steps. Initial matches were obtained using the nuclear-encoded OXPHOS proteins from *Nasonia vitripennis* (Gibson *et al*. 2010; J. D. Gibson unpublished) and *Drosophila melanogaster* (downloaded from www.mitocomp.uniba.it: Porcelli *et al*. 2007). These two protein sets were used as queries to search the protein models predicted for *A. ervi* and *L. fabarum* (blastp, Altschul *et al*. 1997). Here, preference was given to matches to *N. vitripennis*. Next, genes from the *N. vitripennis* and *D. melanogaster* reference set that did not have a match in the predicted proteins were used as queries to search the genome-assembly (blastn), in case they were not in the predicted gene models. Gene models for all matches were then built up manually, based on concurrent evidence from the matches in both *A. ervi* and *L. fabarum* and their available expression evidence. The resulting protein models were aligned to one another and to *N. vitripennis* using MAFFT (Katoh & Standley 2013) to identify missing or extraneous sections. These results were used as queries to search the *N. vitripennis* proteins to ensure that all matches are reciprocal-best-blast-hits. Gene naming was assigned based on the existing *N. vitripennis* nomenclature. Potential duplicates were flagged based on blast-matches back to *N. vitripennis* (Additional Data 10).

### Olfactory genes

#### Odorant-binding proteins (OBPs) and chemosensory Proteins (CSPs)

To identify OBPs based on homology to known sequences, we retrieved 60 OBP amino acid sequences from other Braconidae (namely *Fopius arisanus and Microplitis demolitor)* from GenBank. To this, we added seven OBPs found in a previous transcriptome of *A. ervi* (Patrizia Falabella, unpublished, EBI SRI Accessions: ERS3933807-ERS3933809). To identify CSPs, we used CSP amino acid sequences from more Hymenoptera species (*Apis mellifera*, *Nasonia vitripennis*, *Fopius arisanus and Microplitis demolitor)*. These sets were used as query against *A. ervi* and *L. fabarum* genomes using tblastn (e-value cutoff 10e-3 for OBPs and 10e-2 for CSPs). Genomic scaffolds that presented a hit with at least one of the query sequences were selected. To identify precise intron/exon boundaries, the Braconidae OBP and CSP amino acid sequences were then aligned on these scaffolds with Scipio (Keller *et al*. 2008) and Exonerate (Slater & Birney 2005). These alignments were used to generate gene models in Apollo. Gene models were manually curated based on homology with other Hymenoptera OBP and CSP genes and on RNAseq data, when available. Lastly, the deduced amino acid sequences of *A. ervi* and *L. fabarum* OBP and CSP candidates were then used as query for another tblastn search against the genomes in an iterative process to identify any additional OBPs. Since both OBPs and CSPs are secreted proteins, the occurrence of a signal peptide was verified using SignalP (Emanuelsson *et al*. 2007; Nielsen 2017).

#### Odorant receptors (ORs)

ORs were annotated using available OR gene models from *Diachasma alloeum*, *Fopius arisanus*, and *Microplitis demolitor* retrieved from NCBIs protein database (retrieved May 2017). Preliminary OR genes models for *A.ervi* and *L. fabarum* were predicted with exonerate (v2.4.0), GeMoMa (v1.4, Keilwagen 2016), and combined with EVidence Modeler (v.1.1.1, Haas *et al*. 2008). These preliminary models were subsequently screened for the 7tm_6 protein domain (with PfamScan v1.5) and manually curated in WebApollo2.

In an iterative approach, we annotated the IRs using known IR sequences from *Apis melifera*, *Drosophila melanogaster*, *Microplitis demolitor and Nasonia vitripennis* as queries to identify IRs in the genomes of *A. ervi* and *L. fabarum*. The hymenopteran IR sequences served as input for the prediction of initial gene model with Exonerate (Slater & Birney 2005) and GeMoMa (Keilwagen *et al*. 2016). Then, we inspected and edited homologous gene models from each tool in the Apollo genome browser to adjust for proper splice sites, start and stop codons in agreement with spliced RNA-Seq reads. After a first round of prediction, we repeated the whole process and provided the amino acid sequences of curated IR genes as queries for another round of predictions to identify any remaining paralogous IRs.

Multiple sequence alignments of the IRs were computed with hmmalign (Eddy 1998) using a custom IR HMM to guide the alignments (Harrison *et al*. 2018). Gene trees were generated with FastTree v2 (Price *et al*. 2010) using the pseudocount option and further parameters for the reconstruction of an exhaustive, accurate tree (options: -pseudo -spr 4 -mlacc 2 -slownni). Resulting trees were visualized with iTOL v4 (Letunic & Bork 2019), well supported IR clusters and expansions were highlighted by color (branch support > 0.9).

#### Sex Determination

Ortholog searches were performed with tblastn (Altschul *et al*. 1997) against the genomic scaffolds. Hits with an e-value smaller than 1e-20 were assessed, apart from *transformer* and *doublesex* where any hit was surveyed. Doublesex, Transformer-2 and Transformer peptide sequences of *Asobara tabida* (NCBI accessions MF074326-MF074334) were used as queries for the core sex determination genes. This braconid species is the closest relative whose sex determination mechanism has been examined (Geuverink *et al*., 2018). The putative *transformerB* sequence of *A. ervi* was blasted for verification against the transcriptome of *Aphidius colemani* (Peters *et al*. 2017) and a highly conserved fragment was detected (GBVE01021531). Peptide sequences of sex determination related genes to use as queries were taken from *Nasonia vitripennis*: Fruitless (NP_001157594), Sex-Lethal homolog (XP_016836645), pre-mRNA-splicing factor *CWC22* homolog (XP_001601117) and RNA-binding protein 1-like (XP_008202465). Hidden Markov models were not used as gene models because the ensuing peptide predictions did not contain all putative homologs (e.g. *transformerB* in *A. ervi*) due to fragmentation of the scaffolds containing the candidate genes.

#### DNA methylation genes

The genomes were searched with tblastn (Altschul *et al*. 1997) for the presence of potential DNA methyltransferase genes using peptide sequences from *Apis mellifera* and *N. vitripennis* as queries. These species differ in their copy number of *DNMT1*, with two copies (NP_001164522, XP_006562865) in the honeybee *A. mellifera* (Wang *et al*. 2006) and three copies (NP_001164521, XP_008217946, XP_001607336) in the wasp *N. vitripennis (Werren et al. 2010)*. *DNMT2*, currently characterized as EEF1AKMT1 (EEF1A Lysine Methyltransferase 1), has become redundant in the list of DNA methyltransferase genes as it methylates tRNA instead, but was surveyed here as a positive control (*N. vitripennis* NP_001123319, *A. mellifera* XP_003251471). *DNMT3* peptide sequences from *N. vitripennis* (XP_001599223) and from *A. mellifera* (NP_001177350) were used as queries for this gene. Low levels of methylation were confirmed by mapping the whole genome bisulfite sequencing data generated by Bewick *et al*. (2017) back to the *A. ervi* genome assembly.

## List of abbreviations

A, T, C, G, and U: Adenine, Thymine, Cytosine, Guanine, and Uracile, nucleotides
bp: Base Pair
BIPAA: BioInformatics Platform for Agroecosystem Arthropods (bipaa.genouest.org)
BUSCO: Benchmarking Universal Single-Copy Orthologs
CDS: Predicted Coding Sequence
CSD: Complementary Sex Determination
CHC: Cuticular Hydrocarbons
*DNMT*: DNA Methyltransferase genes
CSP: Chemosensory Protein
GO: Gene Ontology
HMM: Hidden Markov Model
HOG: Hierarchical Ortholog Group
IR: Ionotropic Receptor
*LRR*: Leucine Rich Repeat Proteins
Mbp: Mega Base Pairs, or 1,000,000bp
MP: Mate-pair sequence data
NCBI: National Center for Biotechnology Information
N50: A measure of genome completeness. The length of the scaffold containing the middle nucleotide
OXPHOS: Oxidative Phosphorylation
OBP: Odorant-binding Protein
OR: Odorant Receptor
PE: Paired-end sequence data
RSCU: Relative Synonymous Codon Usage
TE: Transposable Element

## Availability of data and materials

Both genomes are available from the NCBI Genome database (PRJNA587428, *A. ervi*: SAMN13190903, *L. fabarum*: SAMN13190904). The assemblies, predicted genes, and annotations are also available at https://bipaa.genouest.org. Raw Illumina and PacBio sequence data used to construct genomes is available in NCBI SRA for both *A. ervi* (SAMN12878248) and *L. fabarum* (accessions SAMN10617865, SAMN10617866, SAMN10617867), and is further detailed in Supplementary Tables 1 and 2. Venom protein data are available via ProteomeXchange with identifier PXD015758.

## Acknowledgements

Thanks to the many people who helped with collections and insect rearing, especially Paula Rodriguez (EAWAG), Laury Arthaud and Christian Rebuf (ESIM, INRA-ISA), Francisca Zepeda-Paulo (UACH), Sebastian Ortiz-Martinez, Cinthya Villegas, and Daniela Sepúlveda (UTALCA). Lucia Briones (UTALCA) and Dominique Cazes (ESIM, INRA-ISA) provided help in with *A. ervi* DNA extractions. Paul Saffert (Uni Potsdam) provided valuable discussion leading to the codon usage analysis. David Pratella (ESIM, INRA-ISA) provided help on the annotation of immune genes.

*Aphidius ervi* sequencing was funded by FONDECYT grant 1130483 and Iniciativa Científica Milenio (ICM) NC120027 (both to Christian Figueroa and Blas Lavandero, Universidad de Talca, Chile), INRA (AIP “séquençage” INRA Rennes, France), and funding from ESIM team (Marylène Poirié, INRA-ISA Sophia Antipolis, France) and BGI (funding to Denis Tagu, INRA Rennes, France). The ESIM team is supported by the French Government (National Research Agency, ANR) through the “Investments for the Future” LABEX SIGNALIFE: program reference # ANR-11-LABX-0028-01. Mark Lammers would like to thank Panagiotis Provataris (ZFMK, Bonn, Germany) for bringing to our attention the existent small whole genome bisulfite sequencing data set for *Aphidius ervi*.

*Lysiphlebus fabarum* data were generated in collaboration with the Genetic Diversity Centre (GDC, with particular thanks to Stefan Zoller and Jean-Claude Walser), ETH Zurich, and utilized the ETH Scientific Computing Cluster (Euler). Orthologs were computed on the University of Potsdam’s High Performance Computing Cluster Orson2, managed by the ZIM. *Lysiphlebus fabarum* sequencing was funded by an SNSF professorship to Christoph Vorburger (grant nrs. PP00P3_123376 and PP00P3_146341). TS acknowledges SNSF grant nr PP00P3_170627. JG and LS acknowledge DFG grant SPP 1819 Rapid evolutionary adaptation (GA 661/4-1, SCHR 1554/3-1).

DNA sequences from the *Myzus* genomes used in comparative analysis of codon usage were downloaded from AphidBase. Funding for *Myzus persicae* clone G006 genomic sequencing was provided by USDA-NIFA award 2010-65105-20558. Funding for *M. persicae* clone O genomic sequencing was provided by The Genome Analyses Centre (TGAC) Capacity and Capability Challenge program (project CCC-15 and BB/J004553/1), from the Biotechnology and Biological Sciences Research Council (BBSRC), and the John Innes Foundation.

## List of additional data files

**Additional Data 1:** details of genetic positions used to construct linkage groups for *L. fabarum*.

**Additional Data 2:** Genbank numbers and taxa information for genome (CDS) graphed in Supplemental Figure 6.

**Additional Data 3:** file detailing (a) the most highly expressed genes in both taxa and (b) differential expression between adult and larval *L. fabarum*.

**Additional Data 4:** fasta file of orphan genes for *A. ervi*

**Additional Data 5:** fasta file of orphan genes for *L. fabarum*

**Additional Data 6:** Summary of OMA output, including details of *LRR* genes

**Additional Data 7:** Annotation of venom genes in *L. fabarum* and *A. ervi*

**Additional Data 8:** Details of immune gene annotation

**Additional Data 9:** Expression details of Osiris genes in *L. fabarum* and *A. ervi*

**Additional Data 10:** Details of annotated OXPHOS genes, including duplications in the assembly

**Additional Data 11:** Details of sex determination gene annotations

