## Supplemental Materials for "Functional insights from the GC-poor genomes of two aphid parasitoids, *Aphidius ervi* and *Lysiphlebus fabarum*"

Dennis *et al*

#### Contents (hyperlinked)

1. Supplementary results and figures
    a. Assemblies
    b. Gene predictions
    c. Transposable Elements
    d. GC content analysis
    e. Gene family evolution
    f. Venom apparatus
    g. Community annotation of individual gene families
2. Extended methods
3. References for Supplementary materials

### Supplemental results and figures

#### Assemblies

In both species, *de novo* assemblies were constructed with a combination of short and long read sequencing. Summary of data used to build assemblies is in Supplementary Tables 1 and 2.

*Supplementary Table 1: Summary of sequencing data* Aphidius ervi. *Coverage based on filtered data and predicted genome size (140MB)(Ardila-Garcia et al. 2010). Hiseq v2000. N/A denotes that PacBio libraries are constructed without inserts.*

| *Library type* | Host Strain | *Platform* | *Filtered reads* | *Read Length (bp)* | *Coverage* | *Insert size* | *SRA Accession* |
|---|---|---|---|---|---|---|---|
| ***Paired End*** | *Sitobion avenae* | Illumina HiSeq | 60,828,295 | 100 | 86x | 350bp | SRR10311786 |
| ***Mate Paired 3kb*** | *Sitobion avenae* | Illumina HiSeq | 14,959,974 | 100 | 21x | 3kb | SRR9326705 |
| ***Mate Paired 5kb*** | *Sitobion avenae* | Illumina HiSeq | 15,122,455 | 100 | 22x | 5kb | SRR9326706 |
| ***Mate Paired 8kb*** | *Sitobion avenae* | Illumina HiSeq | 15,138,323 | 100 | 22x | 8kb | SRR9326707 |
| ***Long Read Fragments*** | *Acyrthosiphon pisum* | PacBio RSII | 622,140 | 6074 (mean) | 27x | N/A | SRR10208676 |

*Supplementary Table 2: Summary of sequencing data* Lysiphlebus fabarum. *Coverage based on filtered data and predicted genome size (128MB) (Belshaw & Quicke 2003). MiSeq reagents v3. N/A denotes that PacBio libraries are constructed without inserts.*

| *Library Type* | *Platform* | *Filtered reads* | *Read Length (bp)* | *Coverage* | *Insert Size (bp)* | *SRA Accession* |
|---|---|---|---|---|---|---|
| ***Paired End*** | Illumina MiSeq | 20,313,003 | 300 | 32x | 169 +/-10 | SAMN10617865 |
| ***Mate Pair 5kb*** | Illumina MiSeq | 5,361,287 | 300 | 25x | 3782 +/- 404 | SAMN10617866 |
| ***Long Read Fragments*** | PacBio RS | 617,092 | 3658 | 42x | N/A | SAMN10617867 |

#### Assemblies

The genome assemblies for *A. ervi* and *L. fabarum* were both constructed using hybrid approaches that incorporated high-coverage short read (Illumina) and long-read (Pac Bio) sequencing (Supplementary Tables 1, 2). This produced two high quality genome assemblies (N50 = 581kb and 216kb for *A. ervi* and *L. fabarum,* respectively). The assemblies have similar total lengths (139MB and 141MB), but different ranges of scaffold-sizes (Table 1, Supplementary Table 3). Both genomes can be accessed via the BioInformatics Platform for Agroecosystem Arthropods (BIPAA), which contains the full annotation report and can be searched via both keywords and blast (bipaa.genouest.org).

The difference in scaffold size-distribution between the two species is likely the result of the different assembly strategies for the two species: while *A. ervi* was assembled using the Illumina short-reads and then scaffolded with the long-read PacBio data, *L. fabarum* was assembled directly from PacBio sequences that had been error-corrected using the Illumina data. This resulted in fewer small scaffolds for the *L. fabarum* assembly, but both genomes have similar number of long scaffolds (>3,000bp, *A.ervi*: 1,503 and *L. fabarum*: 1,698; Table 1). The assembly strategies differed between the two assemblies to optimize the available data. In *A. ervi*, short read MP libraries with multiple insert sizes permitted an assembly strategy that utilized the different insert sizes to produce the best assembly. In *L. fabarum*, MP libraries were only produced with ca. 5kb inserts and these insert sizes were not reliable. Therefore, the *L. fabarum* assembly used this data to error correct the long-reads (PacBio), and this produced the best assembly for this species. Both assembled genomes appear to be largely complete, with 97.6% (*A. ervi*) and 85.1% (*L. fabarum*) of the 1,658 core orthologous BUSCO genes for Insecta (insect_odb9) present in both species (Table 2); differences in the number of duplicated BUSCO genes is possibly due to the different assembly strategies.

We constructed linkage groups for the *L. fabarum* scaffolds using phased SNPs from the haploid (male) son of a single female wasp. This placed the 297 largest scaffolds (>50% of the nucleotides, Supplementary table 4, Supplementary Figure 1), into the expected six chromosomes (Belshaw & Quicke 2003). With this largely contiguous assembly (the six chromosomes plus unincorporated pieces: 1,407 scaffolds altogether), we can show that the two genomes are highly syntenic, with >60k links in alignments made by NUCmer (Kurtz *et al.* 2004) and >350 large syntenic blocks that match the six *L. fabarum* chromosomes to 28 *A. ervi* scaffolds (Supplementary Figures 2 and 3).

*Supplementary Table 3: Detailed report of genome assemblies and gene predictions.*

| | | *A. ervi* | *L. fabarum* |
|---|---|---|---|
| **Overview** | n scaffolds | 5,778 | 1,698 |
| | Total length | 138,951,524 | 140,705,580 |
| | Longest scaffold | 3,671,467 | 2,183,677 |
| | n scaffolds ≥ 1000 bp | 5,777 | 1,698 |
| | n scaffolds ≥ 3000 bp | 1,503 | 1,698 |
| | N50 | 581,355 | 216,143 |
| | n “N”’s | 131,907 (0.09%) | 0 |
| | GC % | 23.7% | 23.8% |
| **Exons** | n Exons | 95,322 | 74,701 |
| | Longest exon | 13,754 | 11,848 |
| | Mean exon length | 311 | 317 |
| **Introns** | n Introns | 74,978 | 59,498 |
| | Longest intron | 27,991 | 19,865 |
| | Mean intron length | 395 | 383 |
| **Genes** | n Genes | 20,226 | 15,170 |
| | Longest gene | 96,195 | 65,965 |
| | Mean gene length | 2,919 | 3,052 |
| | % genome covered by genes | 42.5% | 32.9% |
| **CDS** | n CDS | 20,344 | 15,203 |
| | Longest CDS | 43,731 | 27,132 |
| | Mean CDS length | 1,216 | 1,381 |
| | % genome covered by CDS | 17.8% | 14.9% |
| | GC% in CDS | 31.9% | 29.8% |

*Supplementary Table 4: Summary of BUSCO statistics for the two species based on matches to the insecta_odb9 database of1,658 total BUSCO groups*

| BUSCO statistics | | |
|---|---|---|
| *Whole genome assembly (nucleotide)* | | |
| | ***A. ervi*** | ***L. fabarum*** |
| Complete, single-copy | 1,571 (94.8%) | 1,265 (76.3%) |
| Complete, duplicated | 46 (2.8%) | 146 (8.8%) |
| *Total Complete* | *1,617 (97.3%)* | *1,411 (85.1%)* |
| Fragmented | 13 (0.8%) | 12 (0.7%) |
| Missing | 28 (1.6%) | 235 (14.2%) |
| Predicted genes (CDS, protein level) | | |
| Complete, single-copy | 1,504 (90.7%) | 1,404 (84.7%) |
| Complete, duplicated | 50 (3.0%) | 185 (11.2%) |
| *Total Complete* | *1,554 (93.7%)* | *1,589 (95.9%)* |
| Fragmented | 32 (1.9%) | 14 (0.8%) |
| Missing | 72 (4.4%) | 55 (3.3%) |

### <u>Linkage groups in *L. fabarum*</u>

*Supplementary Table 5: Summary statistics of linkage groups built for the* L. fabarum *genome. Linkage groups based on 1,319 biallelic SNPs.*

| Linkage groups | *n* | bp |
|---|---|---|
| Linkage groups | 6 | 3.7 – 17 Mbp |
| Incorporated scaffolds | 297 | 75,424,286 |
| Unincorporated | 1,401 | 65,310,394 |

*Supplementary Figure 1: Linkage mapping of* L. fabarum *scaffolds (colored in black and white). Based on 1,319 biallelic SNPS from 90 haploid males, all from a single mother.*

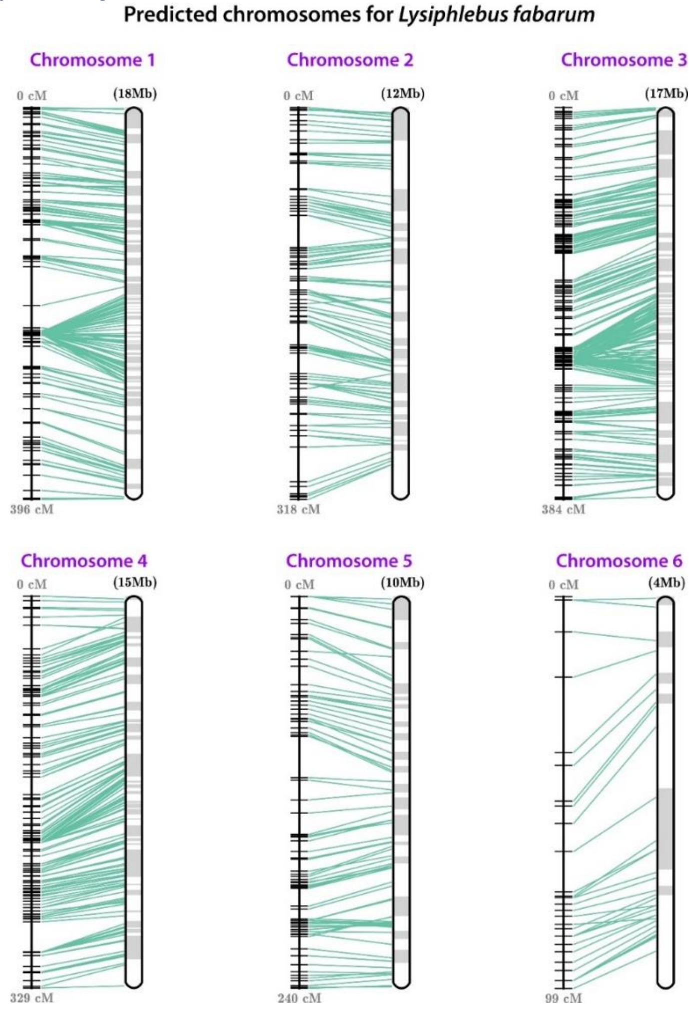

*Gene predictions*

The Maker2 annotation pipeline predicted different numbers of coding genes (CDS) for the two genomes: in *A. ervi* there were 20,344 predicted genes comprising 27.8Mbp, while in *L. fabarum* there were 15,203 genes across 21.9 Mbp. These numbers are on par with those predicted in other hymenopteran genomes (Table 3) and comparisons with these taxa suggest that the lower number of predicted genes in *L. fabarum* are more likely due to its loss than a gain in genes by *A. ervi*. In parallel with the difference in predicted genes, the predicted number of introns and exons was also higher in *A. ervi* (Table 1). In both species, there was high transcriptomic support for the predicted genes, although this was higher for *L. fabarum* (88.3% of genes) than *A. ervi* (77.8%). This difference likely reflects the available transcriptomic data rather than differences in the success of gene predictions. The resulting peptides were further functionally annotated against the NCBI *nr* database (NCBI National Center for Biotechnology Information), matched to gene ontology (GO) terms, and predictions for known protein motifs, signal peptides, and transmembrane domains (Supplemental Table 5). We also re-ran BUSCO to measure the presence of known, core, orthologous genes within the predicted proteins. At the protein level, BUSCO matches from the predicted genes were increased for *L. fabarum* (*A. ervi*: 93.7%, *L. fabarum*: 95.9%) over the nucleotide-level search of the whole genome (*A. ervi*: 97.6%, *L. fabarum*: 85.1%, Supplementary Table 2).

*Supplementary Table 6: Summary of functional annotation of peptides.*

| Proteins with: | *A. ervi* | *L. fabarum* |
|---|---|---|
| Blast match to *nr* | 13, 582 | 12,832 |
| GO annotations | 5,870 | 5,965 |
| Interproscan domains | 14,901 | 12,792 |
| Signal peptides | 1,731 | 1,474 |
| Transmembrane domains | 3,957 | 3,254 |

#### Synteny between the two genomes

We identified syntenic regions between the two genomes by mapping with NUCmer, which is part of the MUMer package (Kurtz *et al.* 2004). NUCmer identified 67,557 matches among all scaffolds in the two genomes (Supplementary Figure 2). To further examine large synteny blocks within this, we created a plot using only the six predicted *L. fabarum* chromosomes and the *A. ervi* scaffolds longer than 1MBp. The matches between these were filtered to retain only instances with at least three consecutive matches of >250bp. This produced 358 connections (each of which represent multiple NUCmer matches), and linked 28 *A. ervi* scaffolds to the six *L. fabarum* chromosomes (Supplemental Figure 3).

*Supplementary Figure 2: Whole-genome alignment generated by NUCmer. Forward matches are shown in red and reverse matches in blue.*

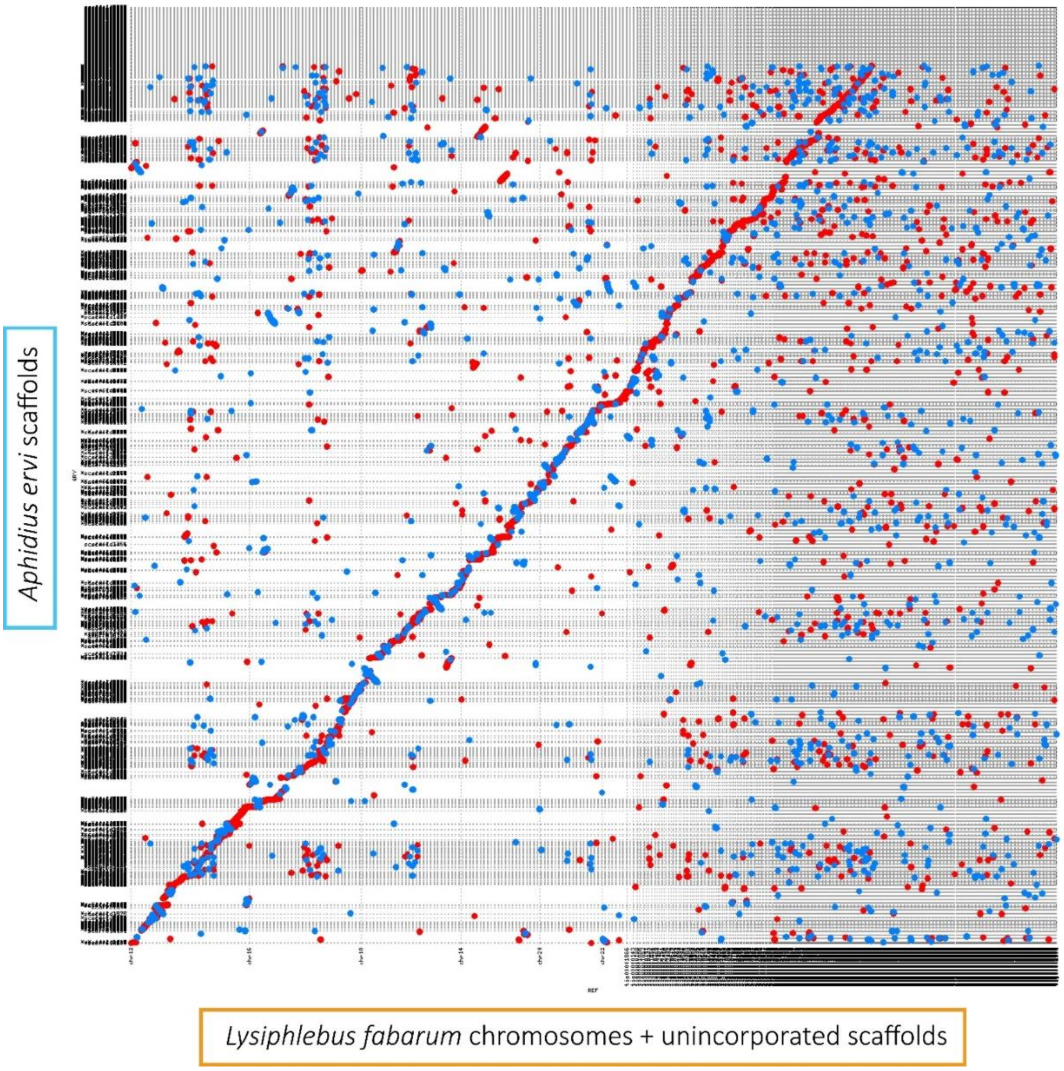

*Supplementary Figure 3: Syntenic region between the six predicted chromosomes in the* L. fabarum *genome and scaffolds in the* A. ervi *assembly. This alignment only included* A. ervi *scaffolds > 1Mbp and instances of at least three consecutive matches with a minimum of 250bp.*

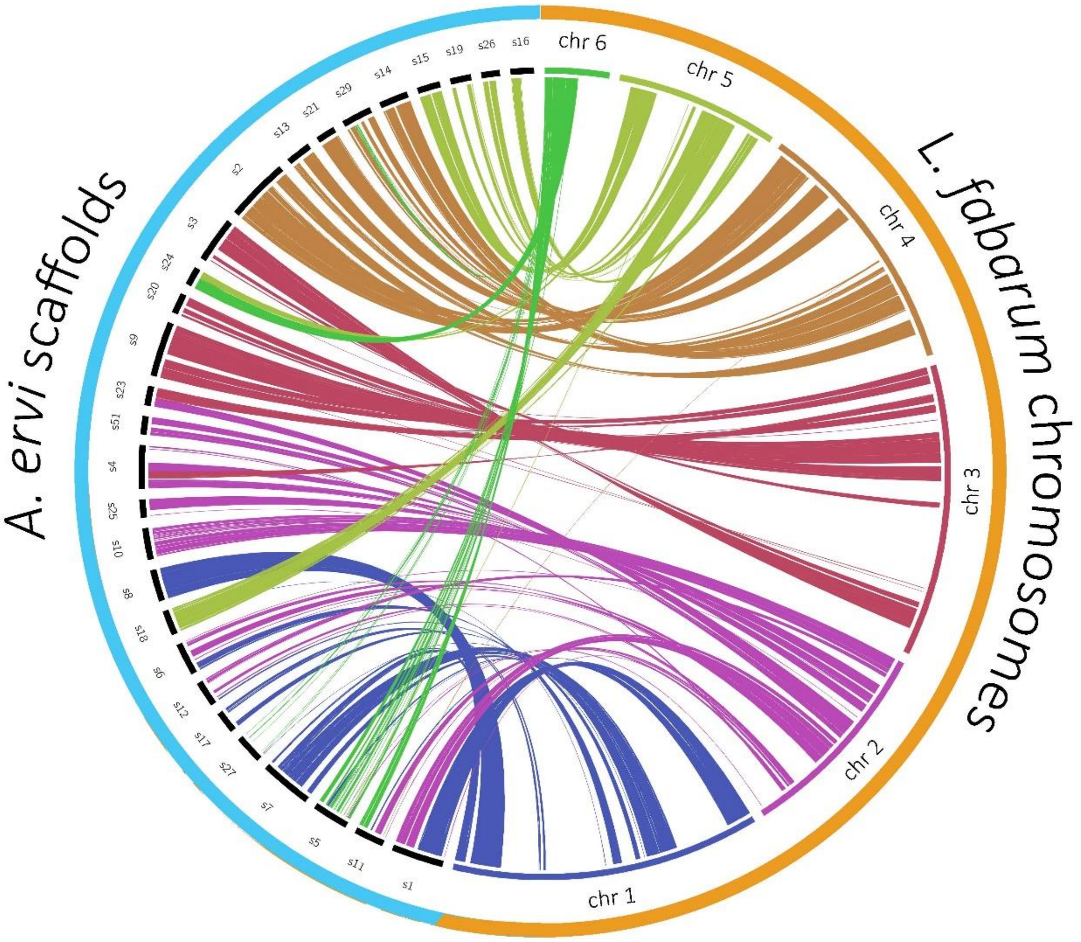

### Transposable Elements (TEs)

Predictive annotation of Transposable Elements (TEs) identified a similar overall number of putative TE elements in the two genomes (*A. ervi*: 67,695 and *L. fabarum*: 60,306, Supplementary table 7). Despite this similarity, the overall genomic coverage by TEs is larger in *L. fabarum* (41%, 58 Mbp) than in A. *ervi* (22%, 31 Mbp). The spread of reported TE coverage in arthropods is quite large, even among *Drosophila* species (ca. 2.7% - 25%, Drosophila 12 Genomes *et al.* 2007). Within parasitoids, reported TE content also varies, and relatively low coverage in the parasitoid *Macrocentrus cingulum* (ca. 18%, Yin *et al.* 2018) in comparison to *Nasonia vitripennis* was attributed to the differences in their genome sizes (127.9Mbp and 295.7Mbp, respectively, Table 3). However, the variation we observe here suggests that differences in predicted TE content may be evolutionary quite labile, even within closely related species with the same genome size.

There is also a difference in the overall types of elements composing the TE's in the two genomes, and in their size distributions (Supplemental Figures 4 and 5, Supplementary table 7). This difference could contribute to the lower GC content in *A. ervi* relative to *L. fabarum*. However, this should also be examined in light of the different assembly strategies in these two genomes. Specifically, that the long-read based assembly of the *L. fabarum* genome may have better reconstructed the large elements than the short-read based assembly in *A. ervi*.

*Supplementary Table 7: Summary of TE annotations. nb= number.*

| | *A. ervi* | *L. fabarum* |
|---|---|---|
| number of sequences | 297 | 1101 |
| number of matched sequences | 297 | 1100 |
| cumulative coverage | 31,015,099 bp | 58,003,749 bp |
| coverage percentage | 22.32% | 41.22% |
| total nb of TE fragments | 67,695 | 60,306 |
| total nb full-length fragments | 3,740 (5.52%) | 5,513 (9.14%) |
| total nb of TE copies | 64,653 | 49,579 |
| total nb full-length copies | 3,869 (5.98%) | 5,698 (11.49%) |
| families with full-length fragments | 297 (100.00%) | 1,096 (99.55%) |
| with only one full-length fragment | 23 | 307 |
| with only two full-length fragments | 33 | 377 |
| with only three full-length fragments | 27 | 129 |
| with more than three full-length fragments | 214 | 283 |
| families with full-length copies | 297 (100.00%) | 1,094 (99.36%) |
| with only one full-length copy | 22 | 282 |
| with only two full-length copies | 27 | 379 |
| with only three full-length copies | 32 | 134 |
| with more than three full-length copies | 216 | 299 |
| mean of median identity of all families | 84.47 +- 8.52 | 92.71 +- 5.54 |
| mean of median length percentage of all families | 49.18 +- 28.57 | 39.99 +- 34.78 |

*Supplementary Figure 4: Distribution of TE classes in* A. ervi

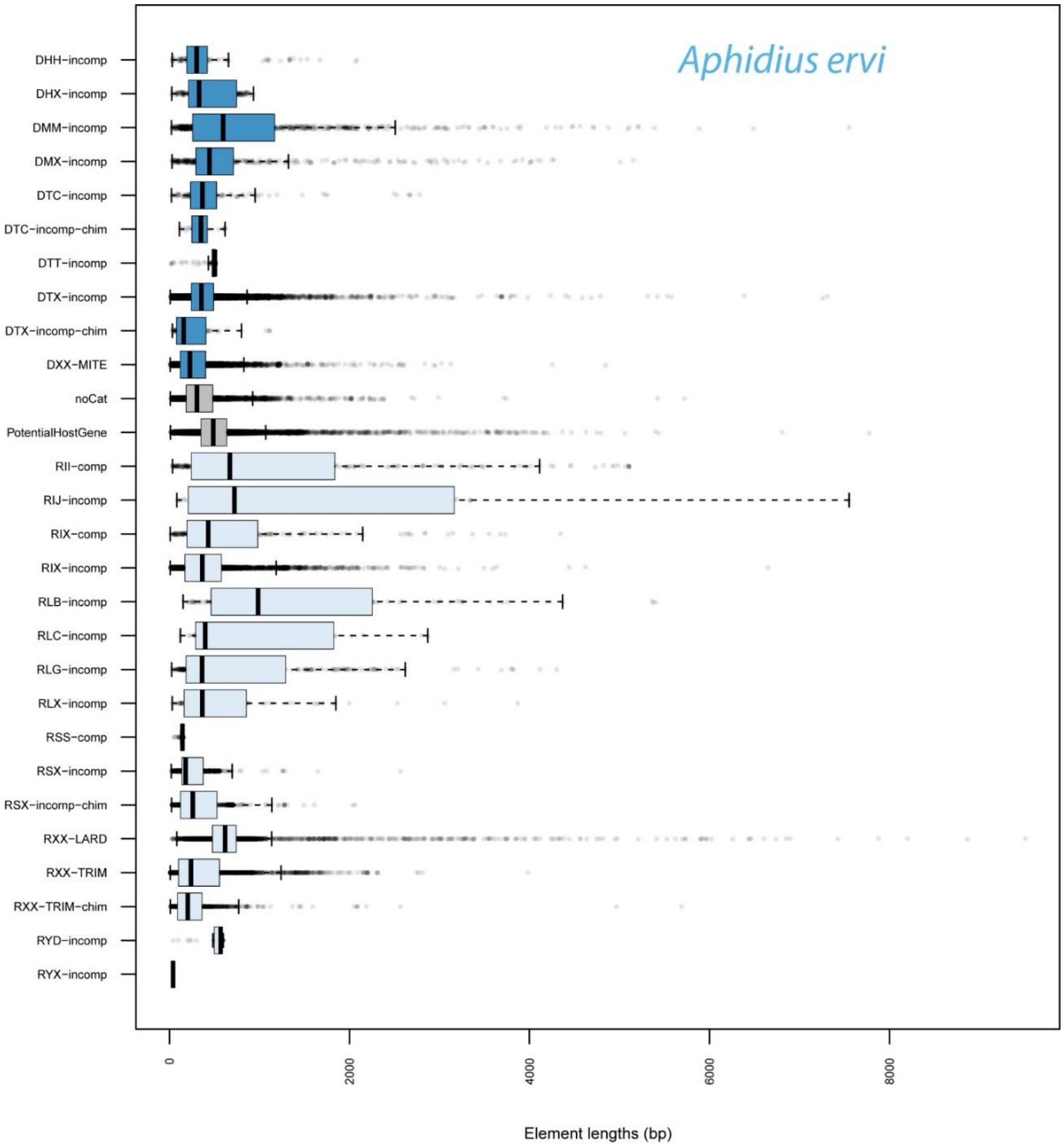

*Supplementary Figure 5: Distribution of TE classes in* L. fabarum

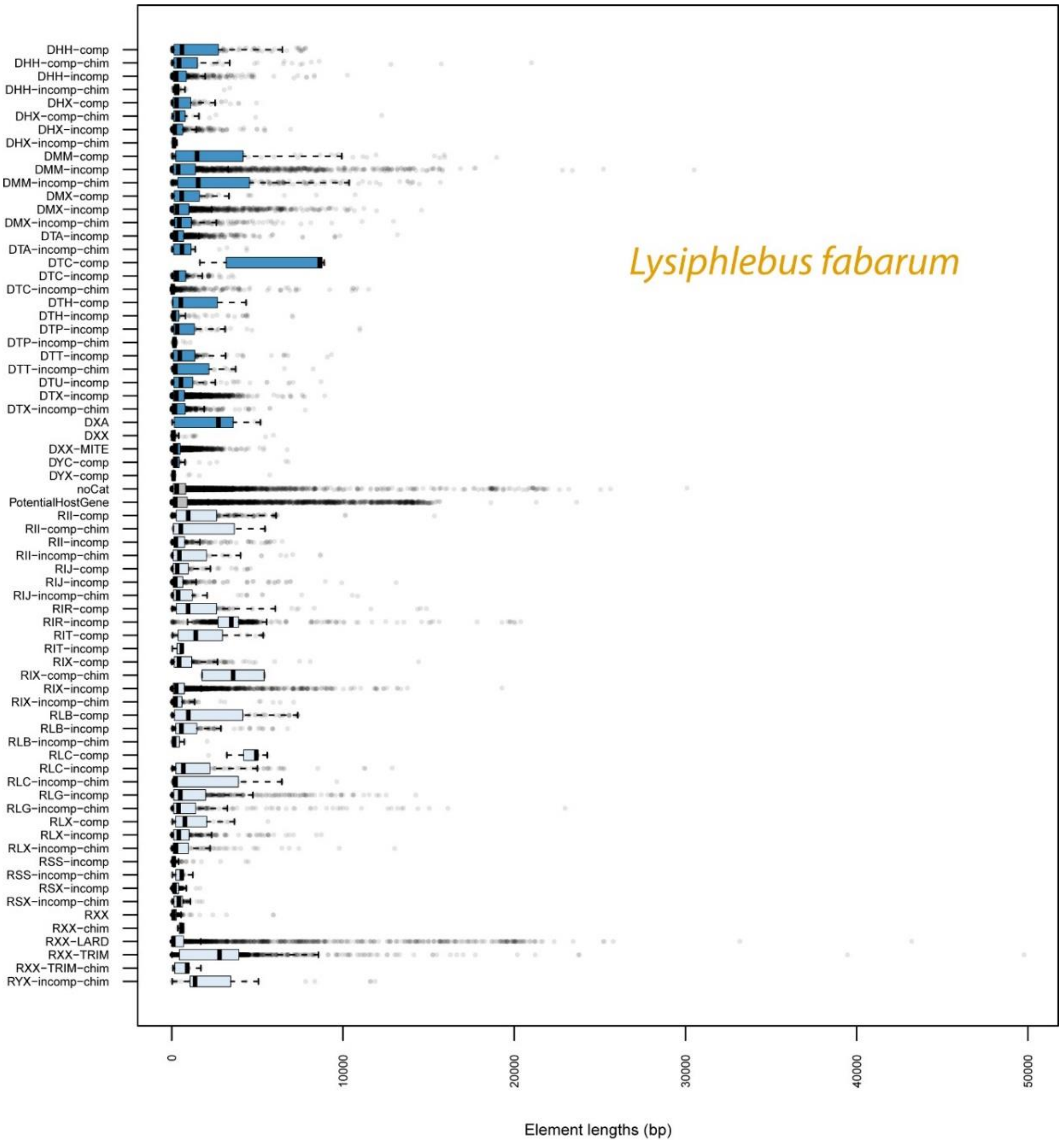

 

### GC Content – in relation to other taxa

We compared GC content within the predicted genes (CDS) of all insect taxa for which this information was available in NCBI (October 2018). These taxa are listed in Supplemental Data File 10.

*Supplementary Figure 6: Boxplot of GC content in all predicted genes. Boxes represent the upper and lower quartiles of the data and their midline depicts the median. Vertical whiskers depict 1.5 the interquartile range. Outliers are shown as black dots. Taxa represented are all Insecta available in the RefSeq RNA database downloaded from NCBI in October 2018, plus the two genomes presented in this paper (far right, colored).*

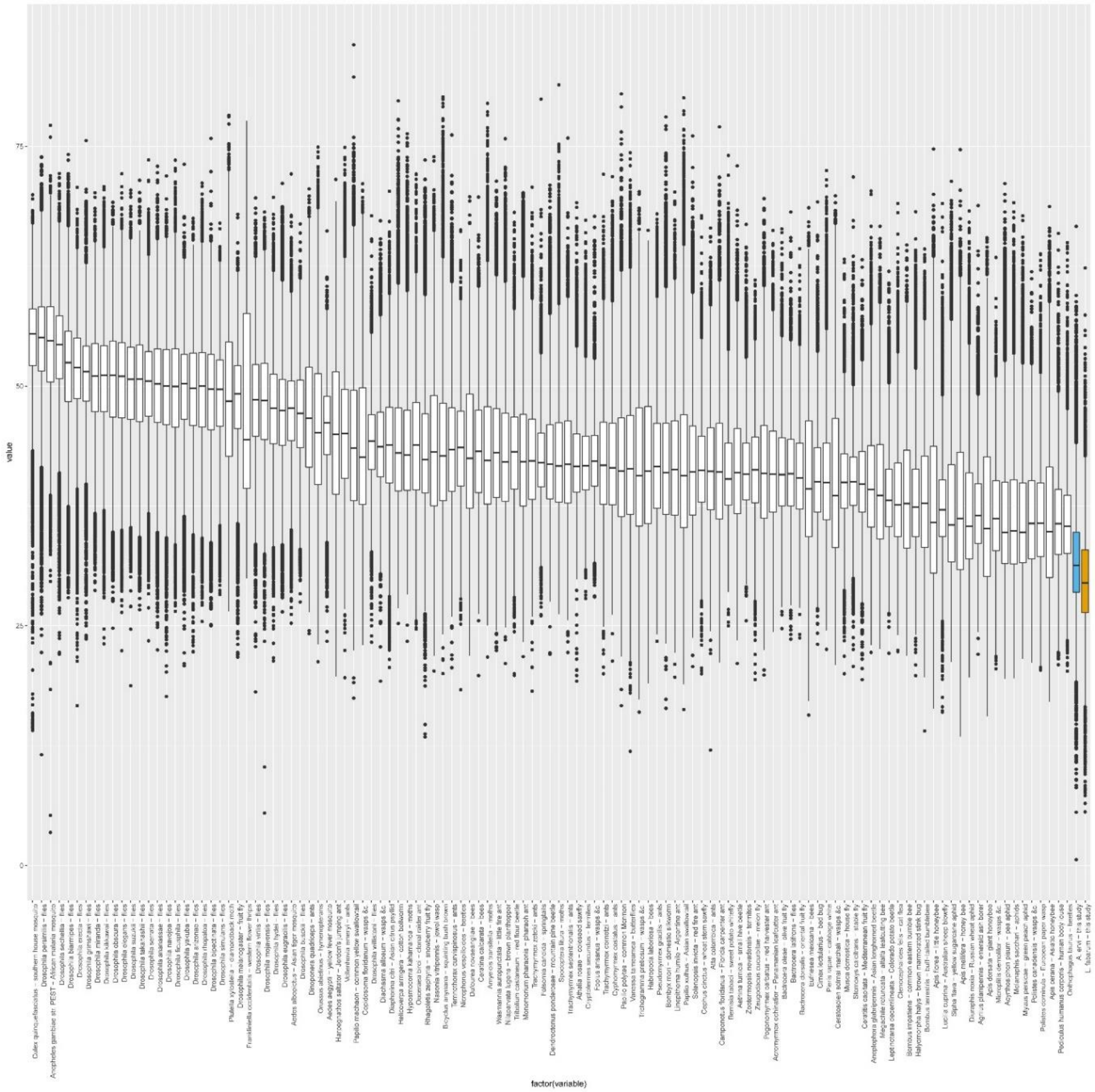

### GC content – in relation to the environment

To examine potential impacts of nutrient availability in the environment (namely nitrogen and carbon), we calculated several aspects of the chemical composition of expressed genes in both genomes. We compared the most highly (5%) and lowly (5%) expressed genes in both genomes, based on the idea that highly expressed genes would have higher material costs (Bragg & Wagner 2009).

We measured:

1) GC content of the expressed genes

The GC content of the most highly expressed genes was significantly higher in both species (Supplementary figure 7, panels a and b). This pattern has been reported broadly in other taxa (Chaney & Clark 2015), and does not suggest that overall expression level is selecting for lower GC content in these two taxa.

2) N-atoms per gene

There is evidence that ecologically limiting nitrogen can select for lower nitrogen content in expressed genes (Acquisti *et al.* 2009). We calculated nitrogen content in the expressed genes by calculating the total N-atoms in the expressed genes. This calculation (i.e. the RNA from our gene predictions) was performed using the weighting used in Acquisti et al (2009), and was conducted as follows: $n_A$= 5, $n_T$ = 2, $n_C$= 3, $n_G$= 5. These comparisons showed that the most highly expressed genes had significantly higher N-content (Supplemental Figure 7, panels c and d). There is a strong relationship between nitrogen content and GC content, because higher GC genes implicitly have higher N-content. Thus, we believe that this pattern is the result of the GC- nitrogen relationship, rather than evidence of selection based on nitrogen limitation.

3) Carbon: nitrogen ratios

As another comparison of nitrogen content, we scaled the nitrogen content by the carbon content. This was intended to correct for overall size of the amino acid and aid visualization (Elser *et al.* 2006). As with the total N-atoms per gene, we see no evidence for selection on lower-nitrogen content. Instead, the most highly expressed genes have the a lower ratio of carbon:nitrogen (Supplemental Figure 7, panels e and f), meaning that they have higher nitrogen content. Therefore, this does not support the idea that more highly expressed genes are under selection to conserve nitrogen.

A note about our assembly and GC content:

We do not believe the low GC content in these two genome assemblies is a byproduct of an incomplete assembly or some other bias (as has been previously observed in other hymenoptera, see: Elsik *et al.* 2014) for several reasons. Principal among these: (1) we have predicted a high presence of core BUSCO genes, (2) the numbers of genes are similar to those predicted in hymenoptera with higher GC content (Table 3), (3) we have incorporated long-read technology (PacBio), which should not have the same GC biases as Illumina-reads (Gan *et al.* 2019; Rhoads & Au 2015), and (4) separate transcriptomic studies in both species have produced transcripts with similar GC content (Ballesteros *et al.* 2017; Dennis *et al.* in revision; Dennis *et al.* 2017)

*Supplementary Figure 7: Content of top 5% of expressed genes in* L. fabarum *(n=1,374) and* A. ervi *(n=2,035)*

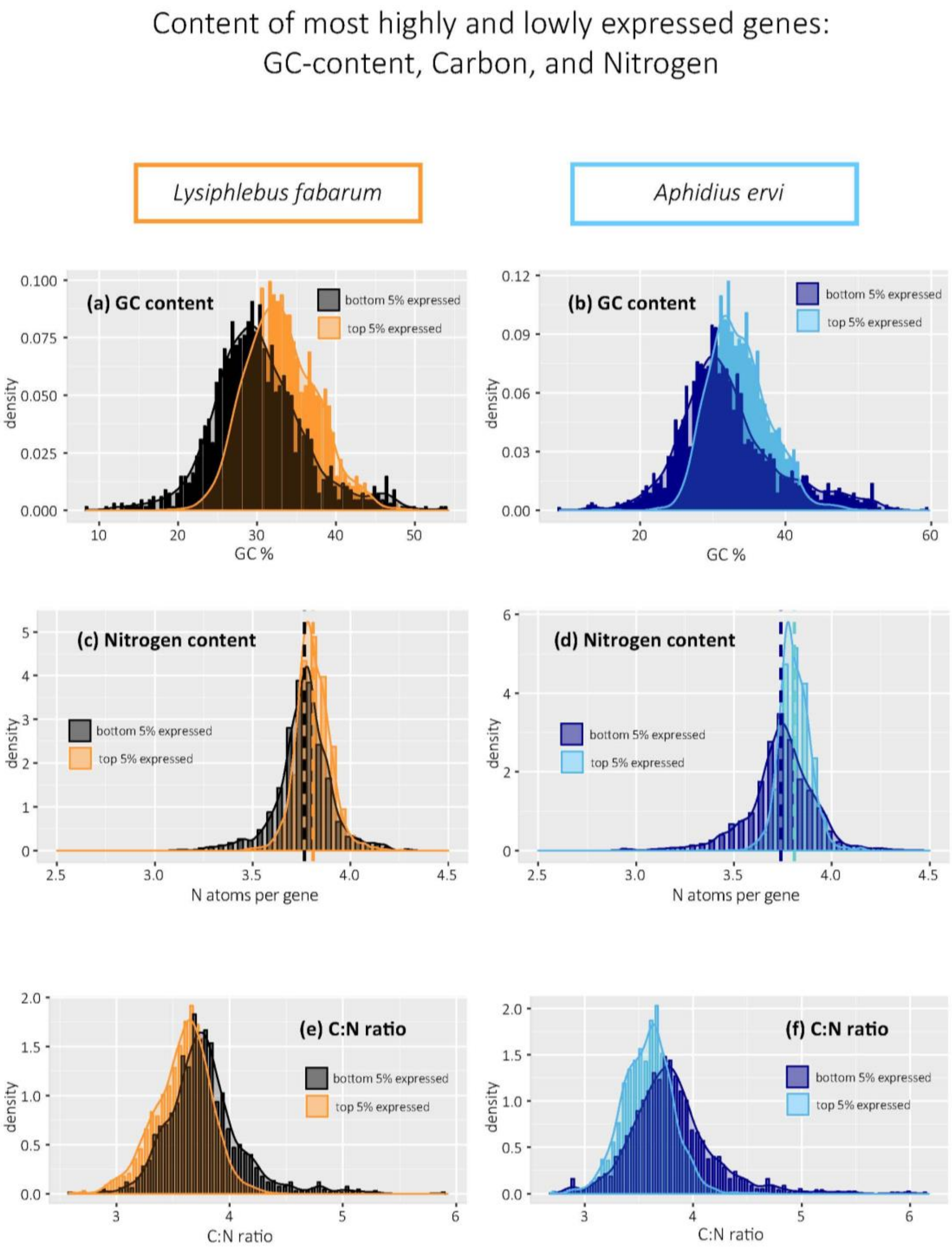

 

### GC content: Comparisons of genomic content in relation to other taxa

To examine the possibility that codon usage in this system is related to their environment and/or host, we examined RSCU, nitrogen content, and nitrogen:carbon ratios in *A. ervi* and *L. fabarum*, in comparison to other taxa from this system: namely four different aphid hosts and their endosymbionts, as well as the model parasitoid *N. vitripennis*. All calculations used only the predicted coding genes. Calculations of RSCU, nitrogen (N) and carbon (C) content, and PCA were performed as detailed above for the two parasitoid genomes.

*Supplementary Table 8: Summary of taxa used in comparative analysis of codon usage, GC-content, and C:N ratios*

| *Group* | *Species* | *Source* |
|---|---|---|
| *Host aphid* | *Aphis glycines* | Wenger *et al.* (2017) |
| | *Acyrthosiphon pisum* | Thorpe *et al.* (2018) |
| | *Myzus persicae* – clone G006 | Legeai *et al.* (2010) |
| | *Myzus persicae* – clone O | Legeai *et al.* (2010) |
| *1° endosymbiont* | *Buchnera aphidicola* from *A. glycines* | Cassone *et al.* (2015) |
| | *B. aphidicola* from *A. pisum* | Shigenobu *et al.* (2000) |
| | *B. aphidicola* from *M. persicae* strain W106 | Jiang *et al.* (2013) |
| *2° endosymbiont* | *Hamiltonella defensa* from *A. pisum* | Degnan *et al.* (2009) |
| *Parasitoid wasp* | *Nasonia vitripennis v2* | Rago *et al.* (2016) |
| | *Lysiphlebus fabarum* | *this study* |
| | *Aphidius ervi* | *this study* |

*Supplementary Figure 8: Relative Synonymous Codon Usage (RSCU) is not similar to other taxa in this system, nor to the parasitoid wasp* Nasonia vitripennis. *Note: two of the date points for* Buchnera *overlap completely.*

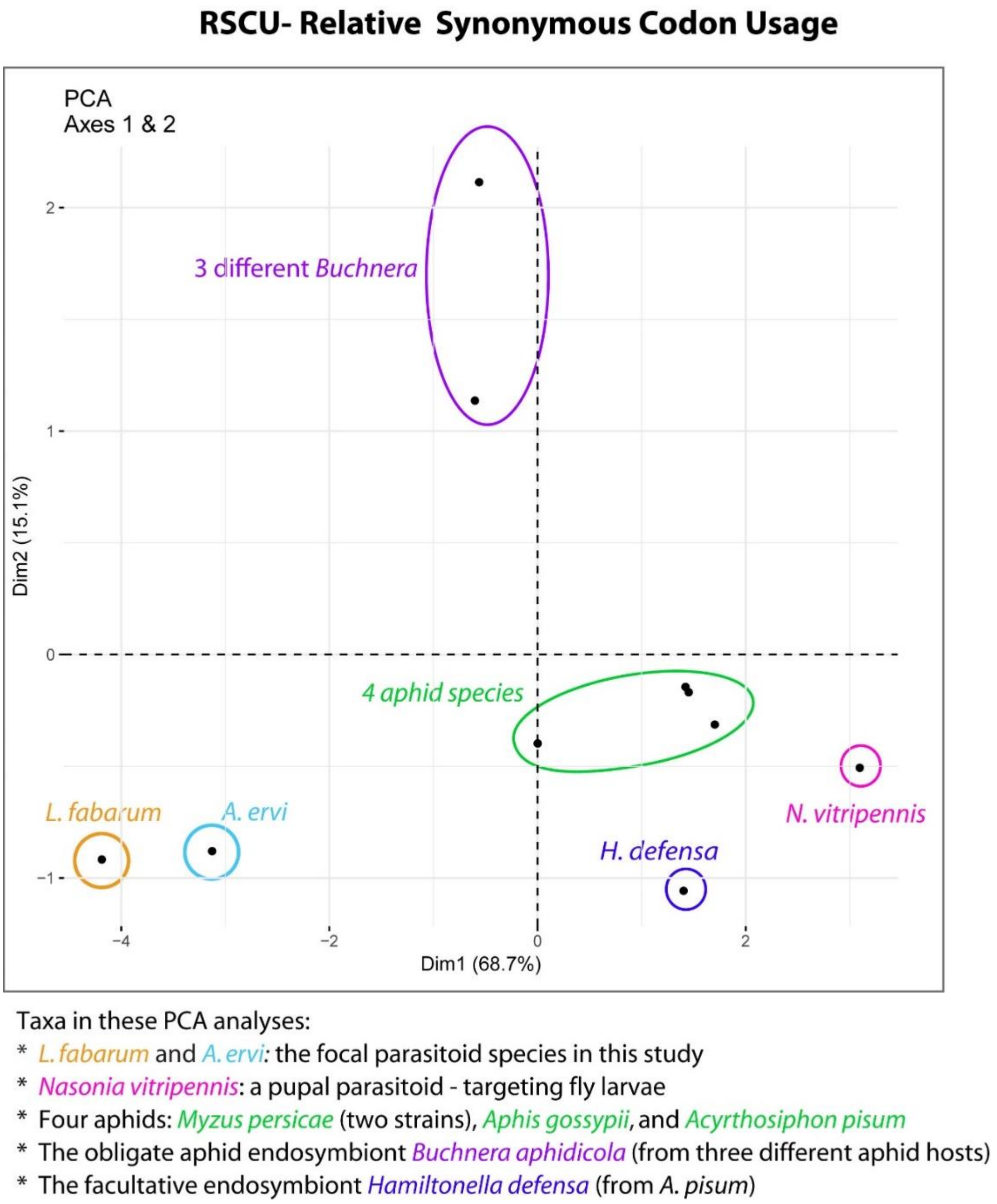

*Supplementary Figure 9: Ratio of carbon: nitrogen in the CDS of all predicted genes, shows little relationship among taxa in this system, nor to the parasitoid wasp* Nasonia vitripennis*.*

**C:N ratio in all codons**

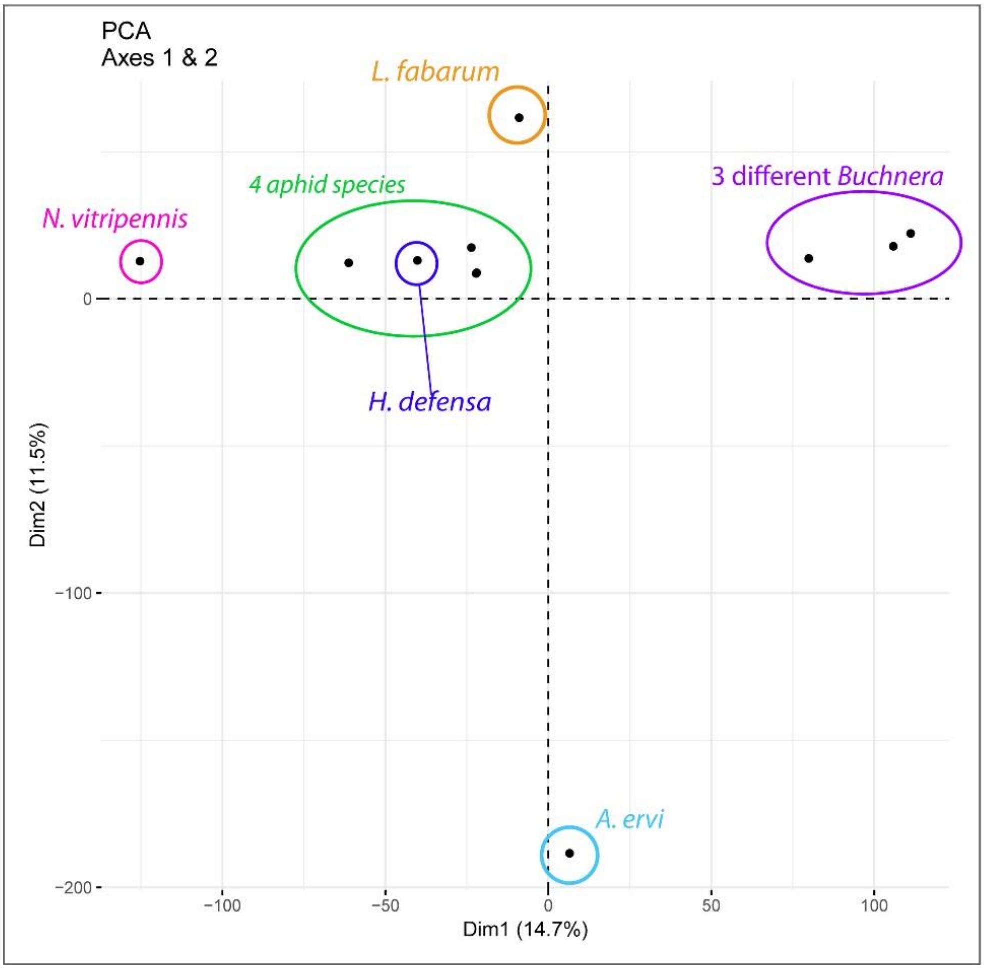

Taxa in these PCA analyses:

* *L. fabarum* and *A. ervi*: the focal parasitoid species in this study
* *Nasonia vitripennis*: a pupal parasitoid - targeting fly larvae
* Four aphids: *Myzus persicae* (two strains), *Aphis gossypii*, and *Acyrthosiphon pisum*
* The obligate aphid endosymbiont *Buchnera aphidicola* (from three different aphid hosts)
* The facultative endosymbiont *Hamiltonella defensa* (from *A. pisum*)

*Supplementary Figure 10: Count of total nitrogen, scaled to the number of amino acids (AA) in each predicted gene, for available genomes in this system, and the parasitoid wasp* Nasonia vitripennis*.*

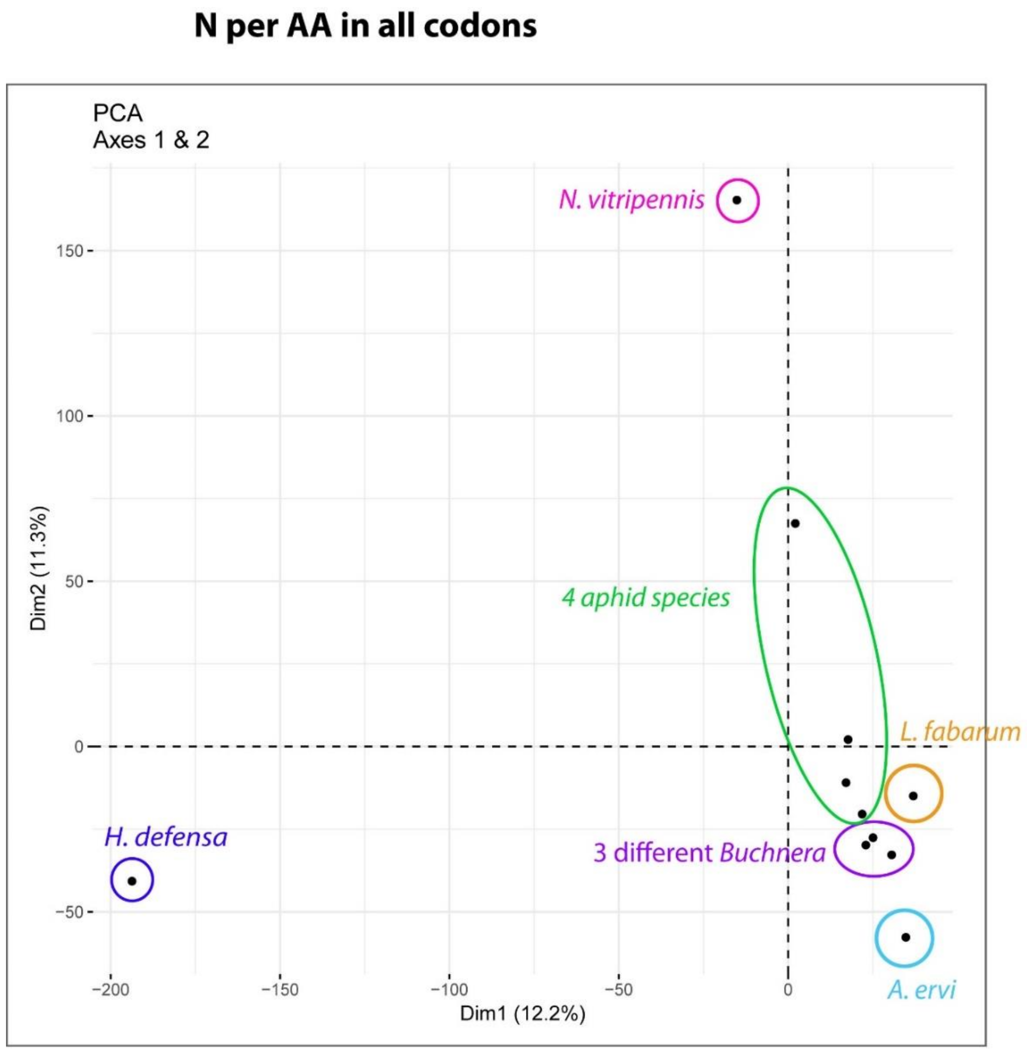

Taxa in these PCA analyses:

* *L. fabarum* and *A. ervi*: the focal parasitoid species in this study
* *Nasonia vitripennis*: a pupal parasitoid - targeting fly larvae
* Four aphids: *Myzus persicae* (two strains), *Aphis gossypii*, and *Acyrthosiphon pisum*
* The obligate aphid endosymbiont *Buchnera aphidicola* (from three different aphid hosts)
* The facultative endosymbiont *Hamiltonella defensa* (from *A. pisum*)

### Gene family evolution

Results from orphan gene analysis.

*Supplementary Table 9: Summary of orphan gene analysis*

| | *A. ervi* | *L. fabarum* |
|---|---|---|
| *Unassigned genes from Orthofinder* | 8,717 | 4,016 |
| *Unassigned genes with no hits in the* nr/nt/swissprot *databases (June 2019)* | 4,306 | 1,280 |
| *Orphan genes with transcriptomic support* | 2,568 | 968 |
| *Maximum length* | 5,844 bp | 2,514 bp |

### Comparative orthology with OMA

Results of OMA analysis, which identified putative expansions in gene families in *L. fabarum* and *A. ervi*, relative to another. To examine only the groups with putative expansions or contractions, we looked at the Hierarchical Ortholog Groups (HOGs) with unequal sizes between the two taxa. There were 865 of these groups in *A. ervi* and 223 in *L. fabarum* (Supplemental Figure 11). Among these only a few (ten groups) had more than 20 genes. We further examined these as putatively interesting in association with adaptive evolution in the two groups (Supplemental Figure 12). Among these, the four largest groups were identified as F-box homologs / Leucine-rich-repeats (Hereafter: F-box/*LRR*), and each of these groups was larger in *A. ervi* than in *L. fabarum*.

Among these, there is some spatial clustering across the genome. The greatest number of these genes appear on Chromosome 1 of *L. fabarum* (26 genes), with 12 of gene less than 10kb apart. In *A. ervi* there are 166 *LRR*s in the scaffolds that are syntenic to Chromosome 1, and these are often highly clustered (59 are less than 100kbp apart, ten are less than 1,000bp apart, Additional File 6). Additional members of this functional group were identified on 125 scaffolds in *A. ervi* and 43 scaffolds in *L. fabarum* (Additional File 6).

Because the *A. ervi* assembly contains more small pieces than *L. fabarum*, we wanted to exclude the possibility that this explains the differences in the number of predicted F-box/LRR genes in these two genomes. To explore this, we examined the size of scaffolds holding the genes predicted to encode F-box/LRR proteins. We conclude that the most abundant F-box/LRRs in *A. ervi* do not appear to be the product of its more fractured assembly. In fact, scaffolds containing LRRs in *A. ervi* are significantly larger than those in *L. fabarum* (Welch two-sampled t-test, p=0.001018)

*Supplementary Figure 11: Size of HOGs and identity of those containing >20 genes*

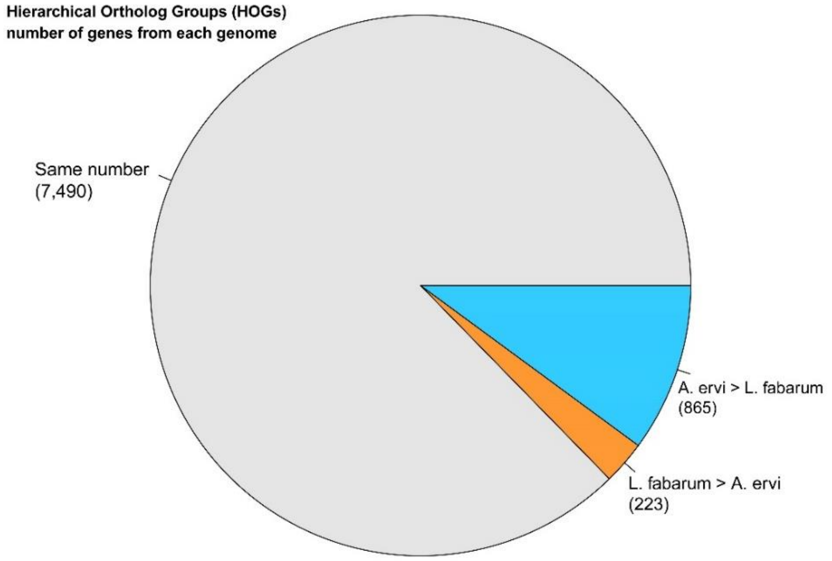

*Supplementary Figure 12: Summary of size of HOGs, and annotation information for the ten largest groups (i.e. all groups with more than 20 genes)*

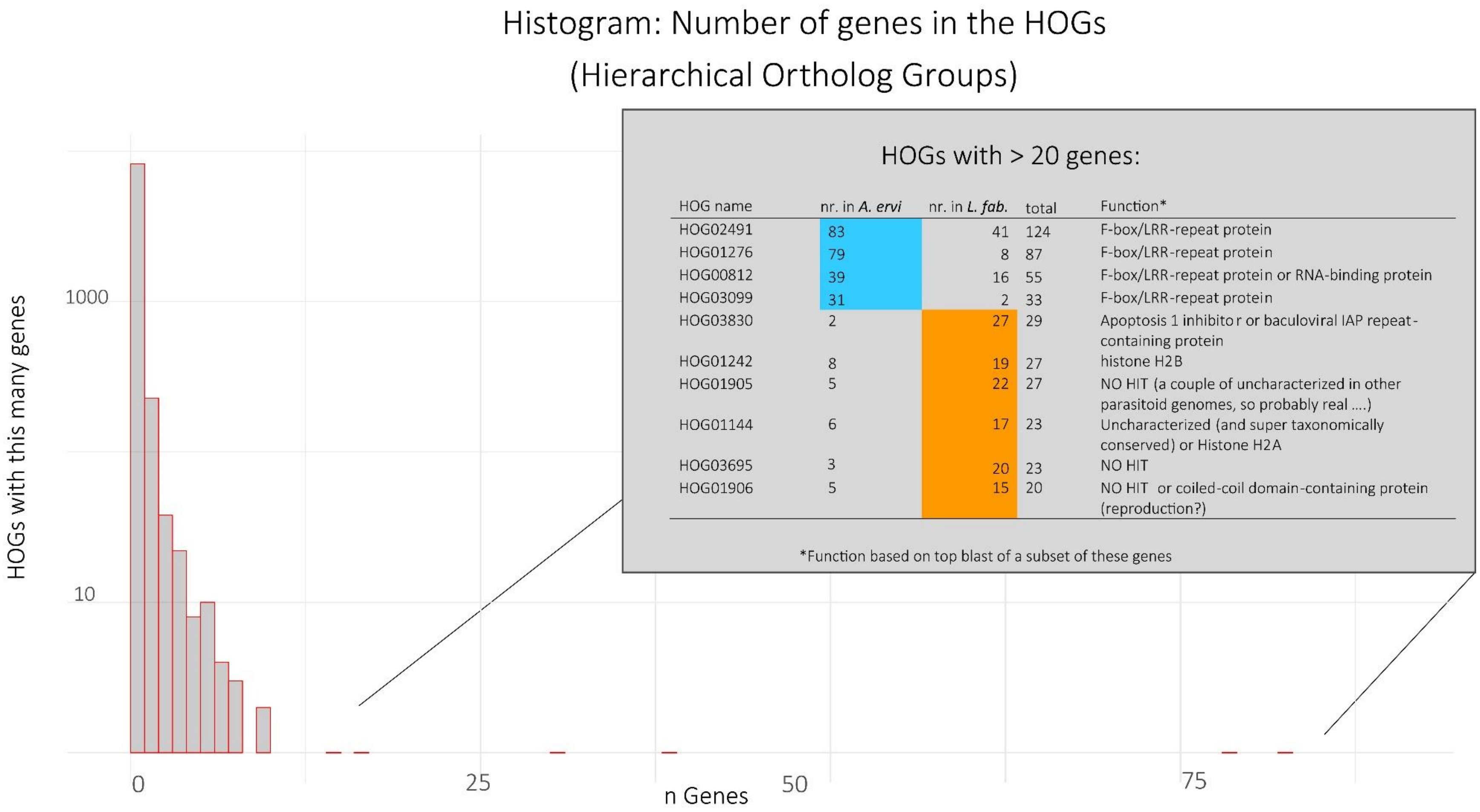

| HOG name | nr. in *A. ervi* | nr. in *L. fab.* | total | Function* |
|---|---|---|---|---|
| HOG02491 | 83 | 41 | 124 | F-box/LRR-repeat protein |
| HOG01276 | 79 | 8 | 87 | F-box/LRR-repeat protein |
| HOG00812 | 39 | 16 | 55 | F-box/LRR-repeat protein or RNA-binding protein |
| HOG03099 | 31 | 2 | 33 | F-box/LRR-repeat protein |
| HOG03830 | 2 | 27 | 29 | Apoptosis 1 inhibitor or baculoviral IAP repeat-containing protein |
| HOG01242 | 8 | 19 | 27 | histone H2B |
| HOG01905 | 5 | 22 | 27 | NO HIT (a couple of uncharacterized in other parasitoid genomes, so probably real ....) |
| HOG01144 | 6 | 17 | 23 | Uncharacterized (and super taxonomically conserved) or Histone H2A |
| HOG03695 | 3 | 20 | 23 | NO HIT |
| HOG01906 | 5 | 15 | 20 | NO HIT or coiled-coil domain-containing protein (reproduction?) |

*Supplementary Figure 13: Size distribution of scaffolds containing F-box/LRR proteins*

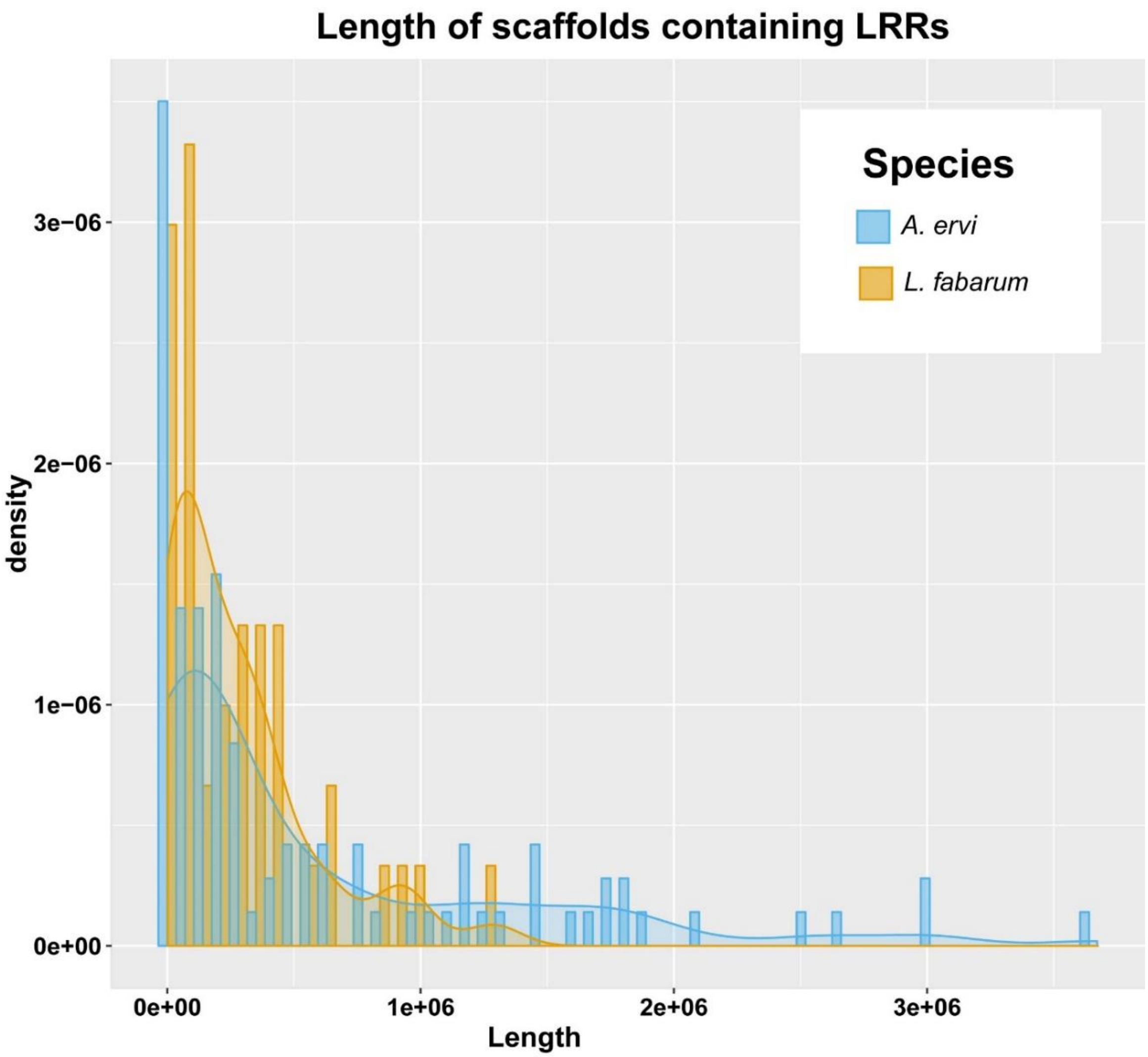

### Venom apparatus

We analyzed the venom proteome in *L. fabarum* using the complete venom glands, including tissues (Supplementary Figure 14). After 1D gel electrophoresis, the 16 major bands (Supplementary Figure 15) were excised and analyzed by mass spectrometry. A combined analysis of the genomic/transcriptomic and proteomic data resulted in 35 putative venom proteins (Supplementary Data 6). Since a number of typical cellular proteins (e.g. actin or myosin) that were identified probably came from venom gland tissues, we only considered putative venom proteins to be the sequences that were: (1) found in proteomics of the venom glands, and (2) either predicted to be secreted or for which the presence of a signal peptide was not tested due to the incompleteness of the sequence. Only 16 putative venom proteins were found in previous work that analyzed of venom proteins for *A. ervi*, but the former approach was more conservative since a protein was considered as venom only if it was found in proteomics not only of the venom glands but also of the reservoir (Colinet et al. 2014). In this work, we applied the less conservative criteria described above to the *A. ervi* previous data. This resulted in 32 putative venom proteins, a similar number to that found for *L. fabarum*. Therefore, we have retained these putative venom proteins for both species (32 in *A. ervi* and 35 in *L. fabarum*).

Comparison of *L. fabarum* venomics data with that of *A. ervi* (Colinet *et al.* 2014) revealed that more than 50% of putative venom proteins are shared between the two species when taking into account these less conservative criteria (Figure 4, Supplementary Data 6). Interestingly, a gamma glutamyl transpeptidase (GGT1) is the most abundant protein in venom of both *A. ervi* (Colinet *et al.* 2014) and *L. fabarum* (Supplementary Data 6, Supplementary Figure 15). As in *A. ervi*, a second GGT venom protein (GGT2) containing mutations in the active site was found in *L. fabarum* (Supplementary Figure 16). Phylogenetic analysis revealed that GGT1 and GGT2 venom proteins of the *A. ervi* and *L. fabarum* GGT1 and GGT2 venom proteins grouped together, suggesting that they originated from two successive duplication events that occurred prior to speciation of both species (Figure 5). As previously shown for *A. ervi* only, the GGT venom proteins of *A. ervi* and *L. fabarum* grouped with clade A among the three distinct phylogenetic clades formed by the non-venomous hymenopteran GGT (Supplementary Figure 5, and Colinet *et al.* 2014). In agreement with this, a similar exon structure was observed between the venomous and non-venomous GGT proteins in clade A, with the exception of exon 1 corresponding to the signal peptide of the venomous GGT proteins (Supplementary Figure 16).

*Supplementary Figure 14: Venom apparatus of* L. fabarum

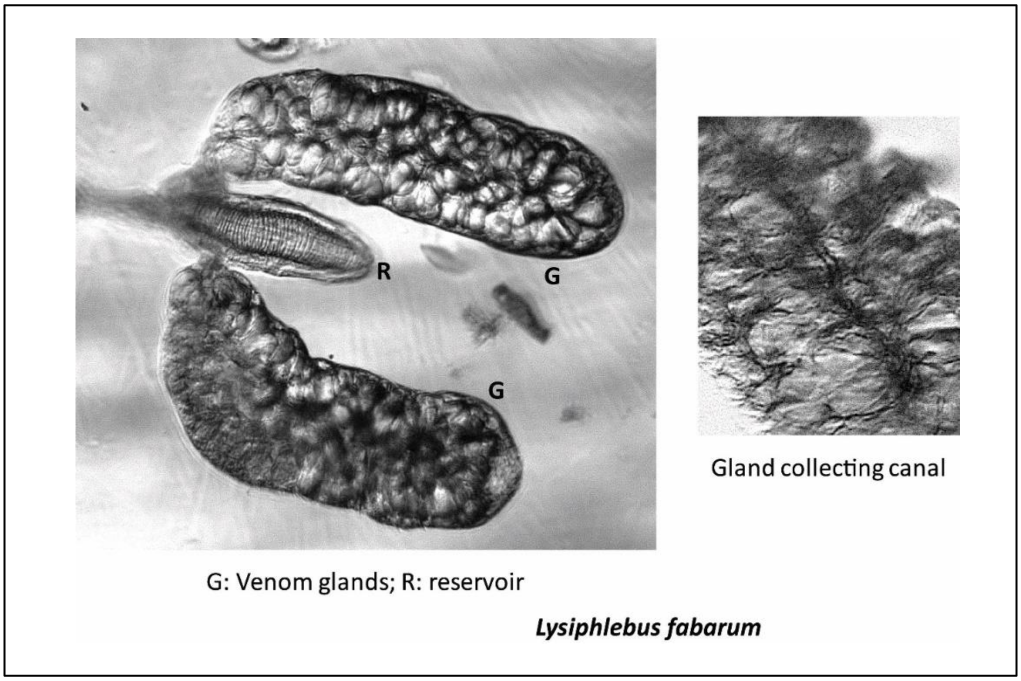

*Supplementary Figure 15: SDS PAGE of the* L. fabarum *venom gland (G) extract. SDS PAGE 12 %, MW, molecular weight standard in kDa. Numbers 1-16 (right side) indicate the 16 most visible bands, which were analyzed by mass spectrometry.*

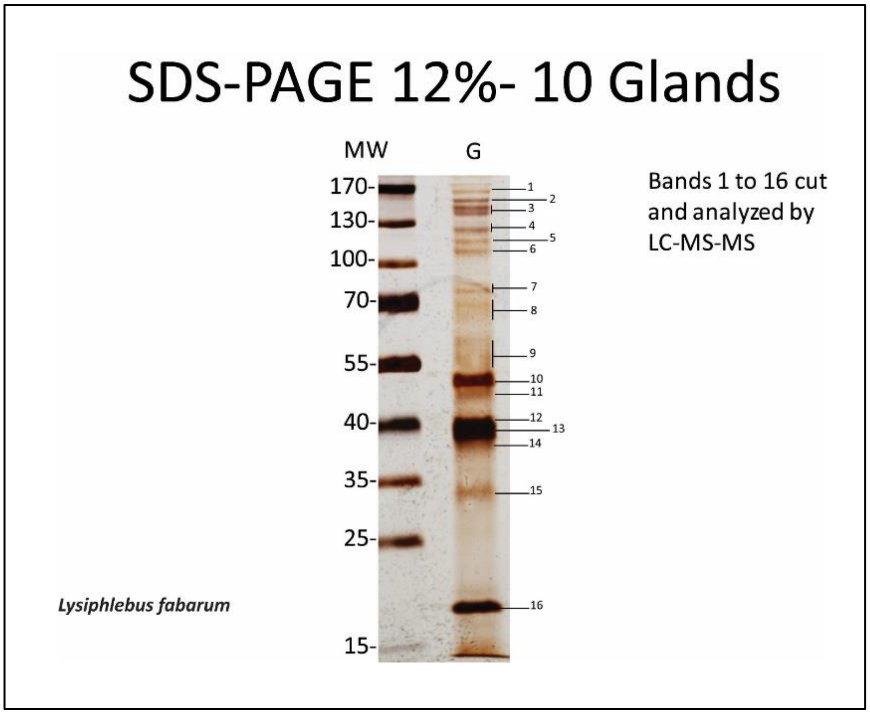

*Supplementary Figure 16: Exon structure of gamma glutamyl transferase (GGT) genes (venom components)*

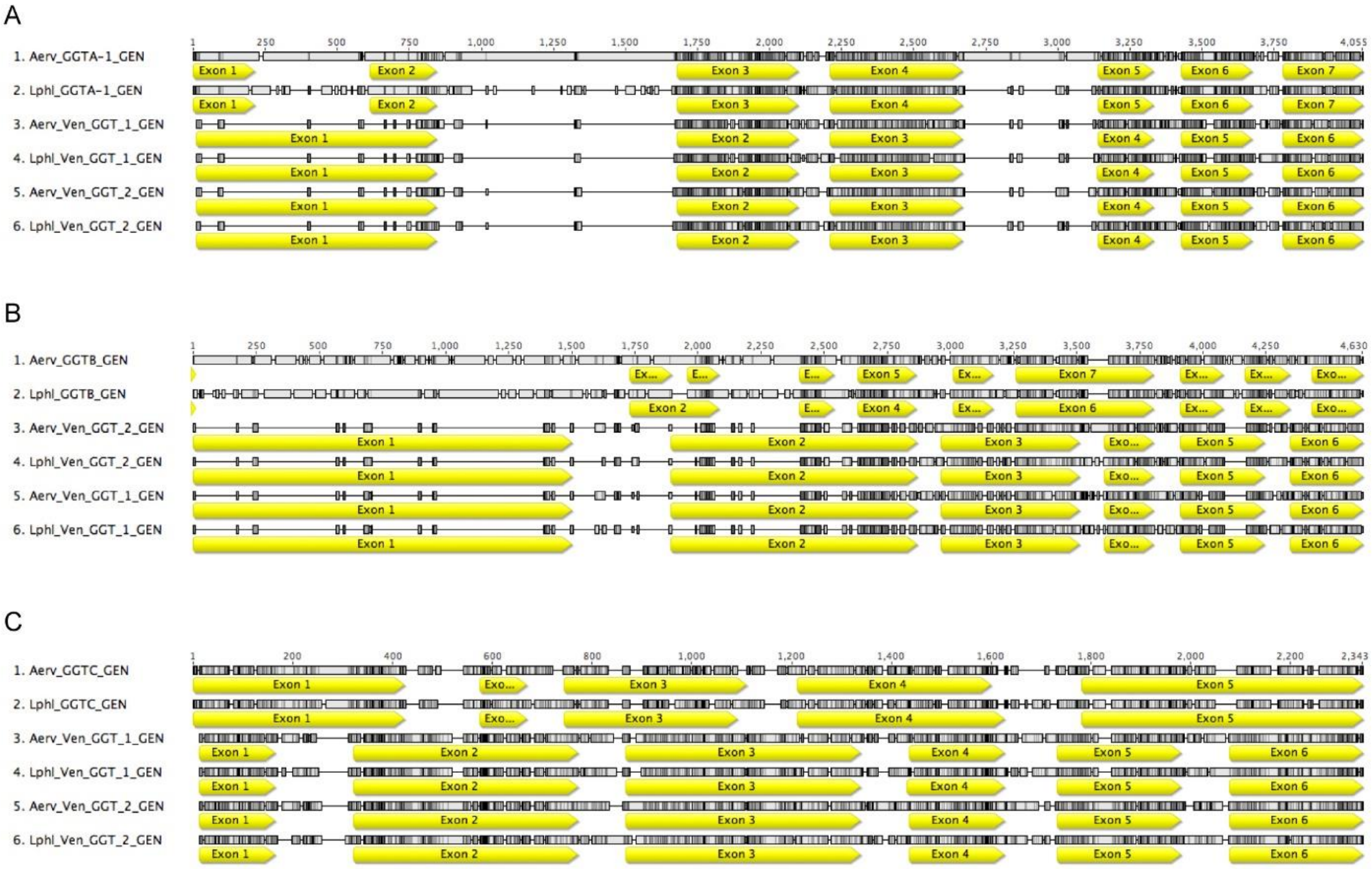

*Supplementary Figure 17: Multiple sequence alignment of venom GGT sequences with the human GGT1 sequence. Stars indicate mutations in the venom GGT_2 sequences described to affect the enzymatic activity of human GGT1. The red rectangle indicates the location of the predicted signal peptide in the venom GGT sequence.*

```
Aerv_Ven_GGT_1    1 MFSKYILTTTVALVLLKYQCY------SADP-SVSAQFKKSAVCAGANKCAEIGSSTLNN
Lphl_Ven_GGT_1    1 MFSKYILTTTVALVLLKYQCY------SADP-SVSGQFKKSAVCAGANKCAEIGSSTLNN
Aerv_Ven_GGT_2    1 MITKFLLS-MIALIFLKYQCN------CETPESVSRNFKKAAVCAGVDECAEIGLSILKK
Lphl_Ven_GGT_2    1 MIRKLILL-IITLMFLKHQCY------CETTSSISRNFKKAAVCAGADQCADIGLLILKK
HSAP_GGT1         1 MKKKLVVLGLLAVVLVLVIVGLCLWLPSASKEPDNHVYTRAAVAADAKQCSKIGRDALRD

Aerv_Ven_GGT_1   54 GGSAVDAAIATMICNNLVHPHLAGYGGGFFMTVYDRANKNVDFLNAREKAP--------G
Lphl_Ven_GGT_1   54 GGSAVDAAIATMICNNLVHPHLAGYGGGFFMTIYDRSSKNVDFLNAREKAP--------S
Aerv_Ven_GGT_2   54 GGTAVDSAIATMICNGLIHMHIAGYGGGFFMTIYQSDTKKVSFLNAKEKSPKETSEYSYK
Lphl_Ven_GGT_2   54 GGTAVDTAIATMICNGLIHMHMAGYGGGFFMTIYQSDTKKVSFLNAKEISPNDTDEISYK
HSAP_GGT1        61 GGSAVDAAIAALLCVGLMNAHSMGIGGGLFLTIYNSTTRKAEVINAREVAPRLAFATMFN
                                                                  *

Aerv_Ven_GGT_1  106 DISKASKTGVNSIAVPGEIAGYGVAHKKYGKLSWEKLFEPTIELCESGYQISKALAKAIA
Lphl_Ven_GGT_1  106 DVSKATKSGVNSIAVPGEIAGYELAHKKYGKLSWEKLFEPTIQLCENGFQVSKALKKAID
Aerv_Ven_GGT_2  114 QLPLDEKTGL-RIAVPGEIAGYAKAHEKFGKIPWSDLFQPTIDLCKKGFKITKTLHDALE
Lphl_Ven_GGT_2  114 DIEPNERNGL-KIAVPGEIAGYAEAHKKFGKLLWSELFQPTIDLCKKGYTITKTLSDALK
HSAP_GGT1       121 SSEQSQKGGL-SVAVPGEIRGYELAHQRHGRLPWARLFQPSIQLARQGFPVGKGLAAALE

Aerv_Ven_GGT_1  166 SSADTIKQDDTLTALYT----NGGALKKEGDTVNPGKLCDTLKTIASGGADEFYKGDLAS
Lphl_Ven_GGT_1  166 ESKDIINANENLKSLYA--------SKNEGDLIKPGKLCETLKKIASGGANAFYKGEVGT
Aerv_Ven_GGT_2  173 KSQSDIENDETLRELFMDKETNSQSLKREGDLVKPTALCETLVTIASKGAGEFYNGTLAK
Lphl_Ven_GGT_2  173 KSQEVIMKDETLRELFMDK--NTELLKKQGDTVNPTNICDTLNTIAKEGANAFYNGTLSK
HSAP_GGT1       180 NKRTVIEQQPVLCEVFC----RDRKVLREGERLTLPQLADTYETLAIEGAQAFYNGSLTA

Aerv_Ven_GGT_1  222 SIIGDLSKKSSALTKDDLSSYTAKWAS-PLKTTLLNDLTLYTANAPGGGAPLALALNIID
Lphl_Ven_GGT_1  218 SIAGDLRKKSGALTKDDLSSYTAEWST-SLKTTLLNDLKLHTANAPGGGATLALCLNIID
Aerv_Ven_GGT_2  233 TIVEDLQSQSSIITEDDFSDYEAEWLE-PLSIKLSNDLTLHTSSVPSSGGLLTLIMNILN
Lphl_Ven_GGT_2  231 TIIEDLKNENSIITEDDLSNYNAEWMK-PLSISLSNNLTLHTSNVPSSGGLLGLMMNVLD
HSAP_GGT1       236 QIVKDIQAAGGIVTAEDLNNYRAELIEHPLNISL-GDVVLYMPSAPLSGPVLALILNILK

Aerv_Ven_GGT_1  281 EYGNDLVSSDAA----LKLHRLSEIWKYSVAAKSKLGDPDVSS--LDELLKTITSDAYAK
Lphl_Ven_GGT_1  277 ELKLNPSSLNLP----STLNSLAEALKHCVARKLSLGDPKSNAQ-VEKEVKTMISDALAK
Aerv_Ven_GGT_2  292 EFNFTPNSLNGTDNTSLTYHKIMESFKWSFPQKYKLGDSKFSKKILDLLVNEFTSKDYAK
Lphl_Ven_GGT_2  290 DINFNSSSLNGTKNTALTYHLIMETWKWTSAQLYKLGDPKFSDKMFDLLAQEFTSKDFAK
HSAP_GGT1       295 GYNFSRESVESPEQKGLTYHRIVEAFRFAYAKRTLLGDPKFVD--VTEVVRNMTSEFFAA

Aerv_Ven_GGT_1  335 EIKSKINDKETSHEASHYGVDVDVQDE-GTAQVSIIDADGNAVSATSSLNGVFGSGVVSE
Lphl_Ven_GGT_1  332 EIRSKINDEKTSNDPKHYGADVDVQDD-GTAQVSIIDSNGNAVSATSSLNGPFGSGVVSE
Aerv_Ven_GGT_2  352 SIKKKINEKKTSNKESDYGGKESVEDQ-GTAQVSVIDSNGNAVSATSSINAPFGSGIVSK
Lphl_Ven_GGT_2  350 SIKNKINYEKTSNTAKTYGGKESVQDH-GTAQVSVIDSDGNAVSATSSINSPFGSGIVSK
HSAP_GGT1       353 QLRAQISDD-TTHPISYYKPEFYTPDDGGTAHLSVVAEDGSAVSATSTINLYFGSKVRSP

Aerv_Ven_GGT_1  394 STGIILNSALSDF-----------SPKSKANEIAPNKRPLSSMAPSIIVDSNDNVKLVIG
Lphl_Ven_GGT_1  391 STGIILNSALSDF-----------TPKSSANSLAPGKRPLSSMAPSIIIDSSDNVRLVIG
Aerv_Ven_GGT_2  411 RTGLIFNNAMDGFWTPTVTGPTGGEETQDGNRIDAQKKPLSSMLPTIITDSNGDVKVVIG
Lphl_Ven_GGT_2  409 RTGLIFNNAMDGFWIPTITGTTDKLTEIDGNRIDANKRPLSSMVPSIITDSNGDVKVVIG
HSAP_GGT1       412 VSGILFNNEMDDFSSPSITNEFG-VPPSPANFIQPGKQPLSSMCPTIMVGQDGQVRMVVG
                               *

Aerv_Ven_GGT_1  443 ATGGAKITTAVSSVLARYLWSKQGLKEAVDAPRLHQEIFPMELSYEE-SSDVITQLKEVY
Lphl_Ven_GGT_1  440 ATGGPKITTAVSLVLARYLWLKQGLQEAVDAPRIHVSLFPMELSYEK-SDDIISKLKTVY
Aerv_Ven_GGT_2  471 GTGGTKIITSVSFVLARYLWMGEDMKTAIDADRIHYQDIIMRVGAEK-MDDVVQNLKRKY
Lphl_Ven_GGT_2  469 GTGGTRILTSVSFVLARYLWMGEDMKTAIDANRIHYRPNVMILRTET-IDDIIKYMKS-Y
HSAP_GGT1       471 AAGGTQITTATALAIIYNLWFGYDVKRAVEEPRLHNQLLPNVTTVERNIDQAVTAALETR

Aerv_Ven_GGT_1  502 KHNTVEMKGITSAICALSREGDSILGVADGRRGGSVQGSN-
Lphl_Ven_GGT_1  499 GHNTVEMKGITSAVCALAREGDHILGVADGRRGGSVKGSN-
Aerv_Ven_GGT_2  530 GHRVVLLKEPHSAICSLSKENGVINGVADGRRGGNVAGIDE
Lphl_Ven_GGT_2  527 GHRTALLKDPDSSICSLTKENNIINGVADGRRGGNIAGIDE
HSAP_GGT1       531 HHHTQIASTFIAVVQAIVRTAGGWAAASDSRKGGEPAGY--
```

### Community annotation of individual gene families

Desaturases: Measurements of cuticular hydrocarbons

*Supplementary Figure 18: Characteristic chemical profile based on a pool of 135 individual* Lysiphlebus fabarum *(in 60ul) and table summarizing average composition of CHC compounds.*

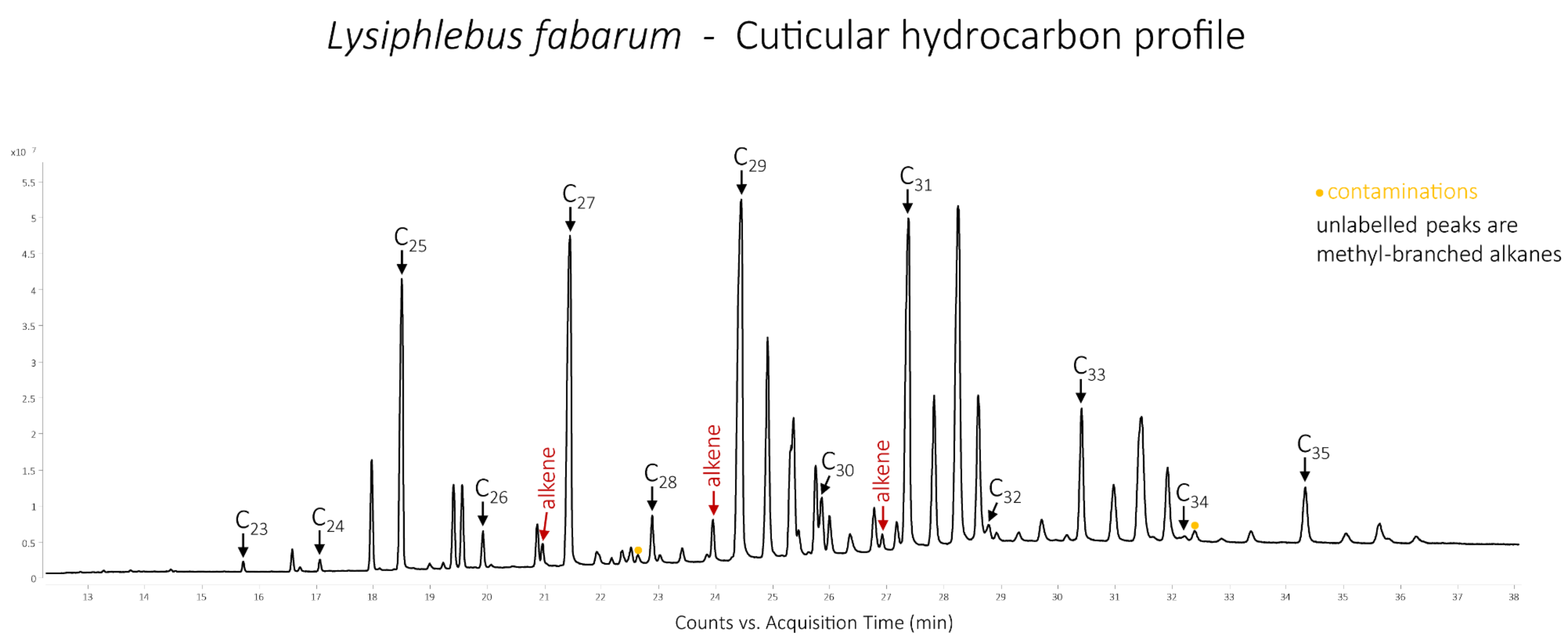

Average CHC compound class ratios (%)

| Alkanes | Methyl-branched alkanes | Alkenes | Dienes |
|---|---|---|---|
| 54.17 | 44.32 | 1.5 | 0 |

*Supplementary Figure 18: Characteristic chemical profile of an individual* Aphidius ervi *female with identifications of the most abundant compounds, and table summarizing average composition of CHC compounds.*

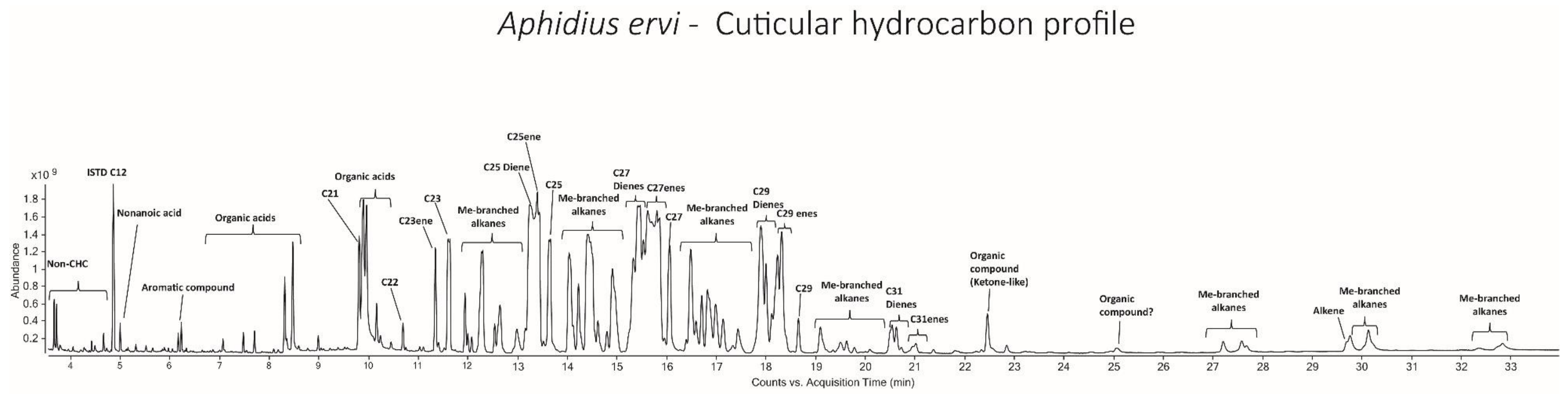

Average CHC compound class ratios (%)

| Alkanes | Methyl-branched alkanes | Alkenes | Dienes |
|---|---|---|---|
| 9.77 | 18.23 | 29.73 | 38.48 |

*Supplementary Figure 19: Comparison of the above CHC profiles from* A. ervi *and* L. fabarum. *Note, that these are aligned for comparative purposes only and as such, the abundance axis has been removed. The traces were produced on different machines and using pooled individuals for* L. fabarum *and a single individual for* A. ervi.

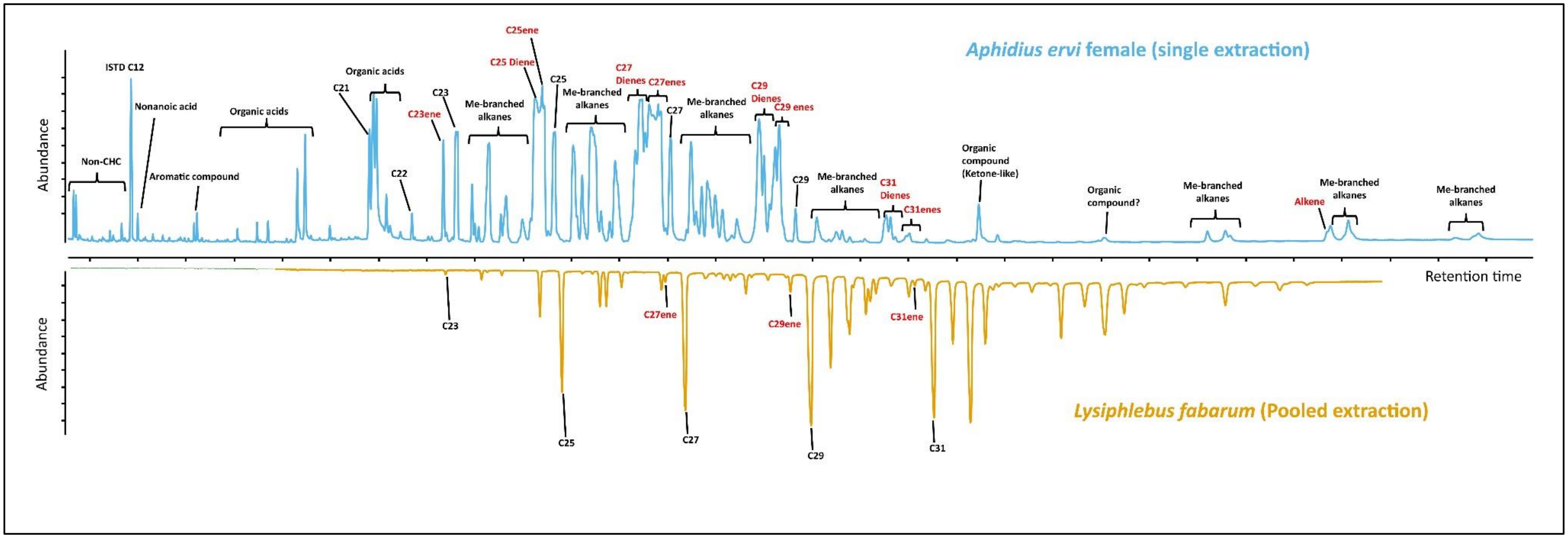

#### Immune genes

The Toll pathway is essential for the response to fungi and Gram-positive bacteria, and it acts via the nuclear factor kappa B (NF-κB, Supplementary Figure 18). Toll-like receptors (TLRs) in this pathway are activated by a cleaved form of an extracellular cytokine-like polypeptide, *spätzle*. Proteolytic cleavage of *spätzle* is the result of a cascade that is initiated by recognition of micro-organisms via pattern-recognition receptors (PRRs, e.g. peptidoglycan recognition proteins, or PGRPs and β-glucan recognition proteins, or βGRPs). The Imd/NF-kappa-B pathway (named for the immune deficiency *imd* gene) is pivotal in the humoral and epithelial immune response to Gram-negative bacteria. It is activated when the receptors' peptidoglycan recognition proteins *PGRP-LC* and *PGRP-LE* bind bacterial peptidoglycan. This initiates signaling to the NF-κB transcription factor *relish*, via the Fas-associated protein with death domain (FADD), the death-related ced-3/Nedd2-like protein (*DREDD*), and the transforming growth factor beta (TGF-β)-activated kinase 1 (*TAK1*), inhibitor of κB kinase (*IKK*) pathways (Buchon *et al.* 2014; Charroux & Royet 2010; Lemaitre & Hoffman 2007). DREiDD cleaves *imd*, which enables its association and activation by the ubiquitin E3-ligase *DIAP2*. *DREDD* also cleaves the Relish precursor, allowing its phosphorylation and translocation into the nucleus where it activates the transcription of specific antimicrobial peptides (AMP). The JAK-STAT pathway is activated after binding of cytokine-like proteins called unpaired (*upd*, *upd2*, and *upd3*) to the receptor *Domeless* (Dome). Activated JAKs phosphorylate each other and provide binding sites for the Src homology 2 (*SH2*) domains of *STAT* molecules. The *STATs* are then phosphorylated by *JAKs* and translocate into the nucleus, where they bind the promoters of target genes. Supplementary Table 10 compares the actors of these major pathways between *A. ervi*, *L. fabarum* and well-studied insect models.

*Supplementary Table 10: Comparison of some immune associated genes in* A. ervi, L. fabarum *and selected insect models. Known copy number for each gene (adapted from: Arp et al. 2016; Evans et al. 2006; Gerardo et al. 2010).*

| | | *D. melanogaster* | *A. mellifera* | *A. pisum* | *A. ervi* | *L. fabarum* |
|---|---|---|---|---|---|---|
| Recognition | PGRP | 13 | 4 | 0 | 1 | 0 |
| | GNBP | 3 | 2 | 2 | 0 | 0 |
| | DSCAM | 1 | 1 | 1 | 1 | 1 |
| Toll | Toll | 9 | 5 | 7 | 3 | 4 |
| | MyD88 | 1 | 1 | 1 | 1 | 0 |
| | Tube | 1 | 1 | 1 | 0 | 1 |
| | Pelle | 1 | 1 | 1 | 1 | 1 |
| | Cactus | 1 | 3 | 1 | 1 | 1 |
| | Dif/Dorsal | 2 | 2 | 2 | 1 | 1 |
| | Spätzle | 6 | 2 | 10 | 1 | 1 |
| Imd | Imd | 1 | 1 | 0 | 0 | 0 |
| | Dredd | 1 | 1 | 0 | 1 | 1 |
| | Tak1 | 1 | 1 | 1 | 1 | 1 |
| | FADD | 1 | 1 | 1 | 1 | 1 |
| | Relish | 1 | 1 | 0 | 1 | 1 |
| JAK/STAT | Basket (JNK) | 1 | 1 | 1 | 1 | 1 |
| | Hopscotch (JAK) | 1 | 1 | 1 | 1 | 1 |
| | STAT | 1 | 1 | 2 | 1 | 1 |
| Effectors | PPO | 3 | 1 | 2 | 1 | 1 |
| | Defensin | 1 | 2 | 0 | 1 | 0 |
| | Cecropin | 5 | 0 | 0 | 0 | 0 |
| | Lysozyme | 13 | 3 | 3 | 4 | 4 |
| | TEP | 6 | 4 | 2 | 3 | 3 |

*Supplementary Figure 20: Comparison with the* D. melanogaster *IMD and Toll pathways components. Red cross, gene absent in both species; Blue cross, absent in* L. fabarum; *Yellow cross, absent in* A. ervi; *Green circle, present in both species. (redrawn from KEGG: www.genome.jp/kegg/pathway.html)*

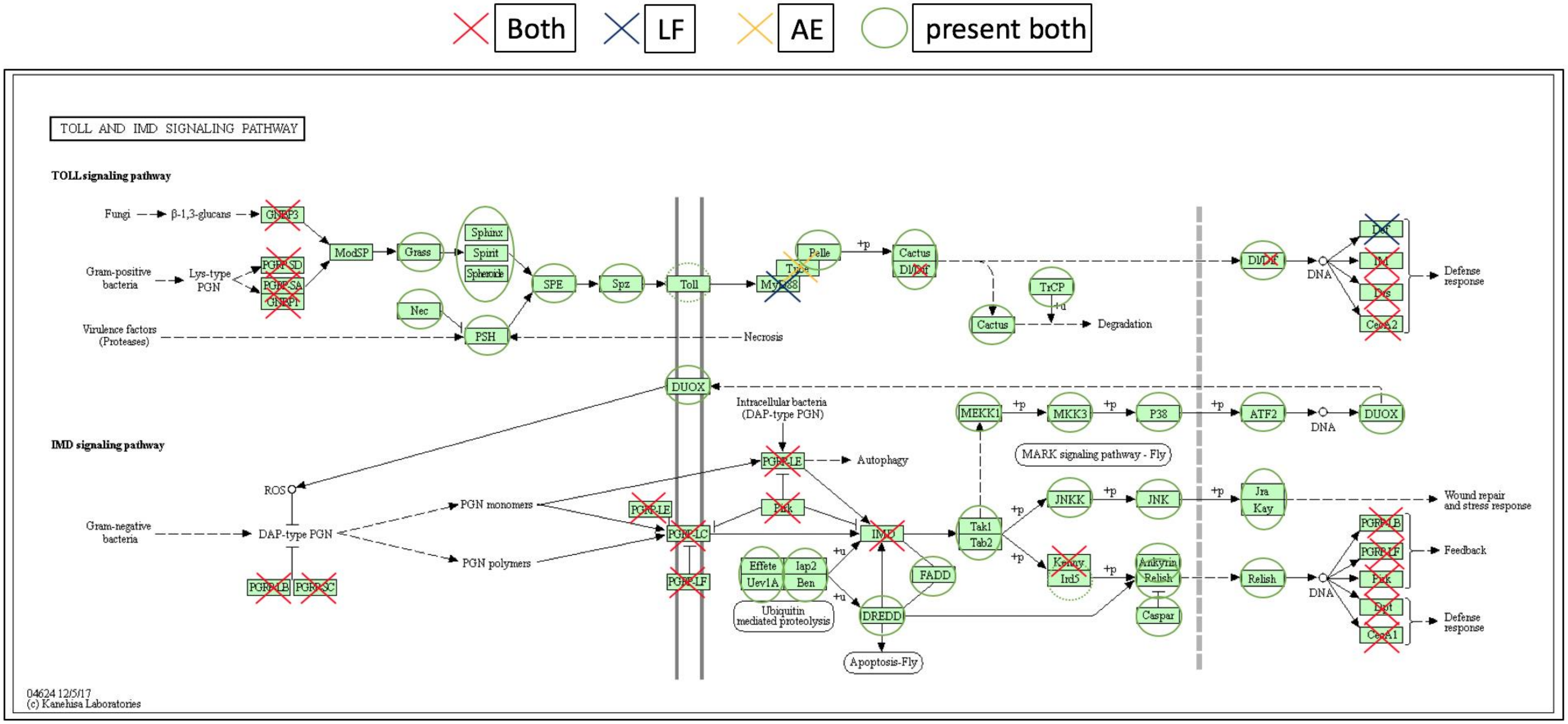

#### Osiris genes

The Osiris genes are an insect-specific gene family that underwent multiple tandem duplications early in insect evolution. All studied lineages of insects have approximately 20 Osiris genes, and most of these occur in a cluster, at least in well-assembled insect genomes (Shah *et al.* 2012; Smith *et al.* 2018). These genes are characterized by a domain of unknown function (DUF1676), a transmembrane domain, a signal peptide and a 3' AQXLAY motif (Shah *et al.* 2012). These genes are essential for proper embryogenesis (Smoyer *et al.* 2003) and for pupation (Andrade López *et al.* 2017; Schmitt-Engel *et al.* 2015). The genes are also tied to immune and toxin-related responses (e.g. Andrade López *et al.* 2017; Greenwood *et al.* 2017), as well as to developmental polyphenism (Smith *et al.* 2018; Vilcinskas & Vogel 2016). Developmental expression patterns are highly conserved for these genes, with major peaks during late embryogenesis and late pupal development (Smith *et al.* 2018), and these genes are co-expressed with suites of genes associated with the regulation of chitin, leading to the hypothesis that these genes are associated with chitin remodeling via the endolytic pathway (Smith *et al.* 2018). Expression levels for all Osiris genes are positively correlated, and correlations increase with both physical proximity and evolutionary distance. The three genes at the end of the cluster, in particular (Osi18-20) tend to be time-shifted in developmental expression relative to the other Osiris genes, suggesting that they are differently regulated (Smith *et al.* 2018); this set of three genes maintains microsynteny in the cluster despite the many inversions and translocations seen within the Osiris cluster across insect genomes (Smith *et al.* 2018).

We found 21 and 25 putative Osiris genes in each the *A. ervi* and *L. fabarum* genomes, respectively (Table 4). In *A. ervi,* these are spread across eight scaffolds, and in *L. fabarum*, seven scaffolds; the position and synteny of genes typically found in the main Osiris cluster are shown in Supplementary Figures 20 and 21. In insects with well assembled genomes, there is a consistent synteny of approximately 20 Osiris genes, and this cluster usually occurs in a ~150kbp stretch. These are, in order, Osi24, Osi1 – Osi20, though there have been some large inversions and translocations, such as in the beetle *T. castaneum* (Smith *et al.* 2018). Gene synteny is conserved in the known Hymenoptera genomes. The middle of this conserved cluster (Osi7 – Osi15) is present as a syntenic group on scaffold237 in *A. ervi*, with Osi16-17 on scaffold397, and Osi18-20 on scaffold759. These scaffolds are relatively short and maybe physically linked. The situation is similar in *L. fabarum*, but the majority of the genes, Osi2-17, are together on scaffold tig00001007 (Supplementary Figure 21). Interestingly, the three genes at the end of the cluster, Osi18-20, appear to have undergone a duplication. These three genes are ancient paralogs (Smith *et al.* 2018) and have a different timing of gene expression during development compared to other Osiris genes (Smith *et al.* 2018). It is hypothesized that these three genes are co-regulated because of their highly correlated levels of gene expression and because their synteny seems to always be preserved despite rearrangements in other parts of the cluster. The Osiris cluster is largely devoid of non-Osiris genes in most of the Hymenoptera, but the assemblies of *A. ervi* and *L. fabarum* suggest that, if the cluster is actually syntenic in these species, there are interspersed non-Osiris genes (those are black boxes in Supplementary Figures 20 and 21).

In support of their role in defense, these genes were much more highly expressed in larvae than in adults of *L. fabarum* (Supplemental Table 12) – larvae live within the adult and we hypothesize that Osiris genes may be part of an adaptive response to dealing with hostile host environment. In both species, transcription in adults was very low, with fewer than 10 raw reads across all cDNA libraries sequenced, and often fewer than one read per library. Across the six sequenced transcriptomic libraries in *A. ervi*, just seven of the 22 putative Osiris genes had non-zero read-counts mapped to them; the highest of these had only 121 mapped reads (Supplemental data 8). In *L. fabarum*, the pattern was similar in adults: across the 59 sequenced adult cDNA libraries, 21 of the 26 predicted Osiris genes had non-zero read-counts, but only six of these had > 10 reads (Supplemental data 8). Patterns of expression were different in developing larvae of *L. fabarum*. Here, across 51 cDNA libraries sequenced from larvae all putative Osiris genes had non-zero read-counts, and many had thousands of reads mapped. Among these, 19 of the 26 annotated Osiris genes were significantly differentially expressed in larvae over adults, with fold changes up to 15x (Supplemental data 6).

*Supplementary Table 11: Annotated Osiris genes (or outgroups) from* A. ervi *as used in phylogenetic reconstruction. The Osi FastTree group refers to Fig. S20. Raw reads mapped came from six cDNA libraries (total reads approximately 1.83 x $10^8$ pairs)*

| *A. ervi* Protein | Osi (FastTree Group) | Raw reads mapped* |
|---|---|---|
| AE3004881 | Out | 4 |
| AE3004882 | OsiC | 20 |
| AE3015979 | Osi15 | 0 |
| AE3015980 | Osi14 | 0 |
| AE3015981 | Osi11A | 0 |
| AE3015984 | Osi10+ | 0 |
| AE3015985 | Osi10+ | 0 |
| AE3015986 | OsiE1/10 | 0 |
| AE3015987 | Osi9 | 0 |
| AE3015989 | Osi7 | 0 |
| AE3017223 | Osi16a | 0 |
| AE3017224 | OsiL | 0 |
| AE3017226 | Osi17 | 0 |
| AE3017262 | Osi1 | 10 |
| AE3017642 | Osi6 | 70 |
| AE3017643 | Osi5 | 0 |
| AE3017820 | Osi18 | 0 |
| AE3017821 | Osi19 | 0 |
| AE3017822 | Osi20 | 0 |
| AE3018336 | Osi2 | 121 |
| AE3018337 | Osi3 | 10 |
| AE3019692 | Osi24 | 41 |

*Supplementary Table 12: Annotated Osiris genes (or outgroups) from* L. fabarum *as used in phylogenetic reconstruction. The Osi FastTree group refers to Supplementary Figure 22. Raw reads mapped came from six cDNA libraries (total reads approximately 2.18 x* $10^9$ *single end). Full results of expression analysis are in Supplemental Data 6.* ns, *no-significant difference.*

| *L. fabarum* Protein | Osi (FastTree Group) | Raw reads mapped larvae | Raw reads mapped adults | Adjusted p-value in expression significance (adults vs larvae) |
|---|---|---|---|---|
| LF003979 | Out | 11932 | 545 | $2.6 \times 10^{-29}$ |
| LF003980 | OsiC | 7921 | 81 | $1.2 \times 10^{-49}$ |
| LF010203 | Osi18 | 16 | 0 | *ns* |
| LF010204 | Osi19 | 25731 | 3 | $9.4 \times 10^{-24}$ |
| LF010205 | Osi20 | 11 | 0 | *ns* |
| LF011148 | Osi17 | 2872 | 5 | $3.2 \times 10^{-22}$ |
| LF011149 | OsiL | 16 | 1 | *ns* |
| LF011150 | Osi16a | 3590 | 9 | $2.1 \times 10^{-17}$ |
| LF011151 | Osi14 | 842134 | 7 | $5.8 \times 10^{-98}$ |
| LF011152 | Osi11A | 9 | 0 | *ns* |
| LF011153 | Osi10+ | 2966 | 3 | $6.8 \times 10^{-21}$ |
| LF011154 | Osi11B | 25 | 4 | *ns* |
| LF011155 | Osi10+ | 1090 | 1 | $4.2 \times 10^{-8}$ |
| LF011156 | Osi9 | 367397 | 44 | $8.7 \times 10^{-102}$ |
| LF011157 | Osi8 | 70487 | 361 | $2.1 \times 10^{-39}$ |
| LF011158 | Osi7 | 36599 | 1 | $7.0 \times 10^{-25}$ |
| LF011159 | Osi6 | 311721 | 45 | $2.5 \times 10^{-95}$ |
| LF011160 | Osi5 | 677 | 7 | $1.4 \times 10^{-10}$ |
| LF011161 | Osi3 | 12190 | 45 | $3.3 \times 10^{-25}$ |
| LF011162 | Osi2 | 5364 | 3 | $1.3 \times 10^{-62}$ |
| LF011915 | Osi23 | 38 | 6 | *ns* |
| LF012925 | Osi1 | 8 | 0 | *ns* |
| LF013026 | Osi20 | 16631 | 5 | $5.8 \times 10^{-22}$ |
| LF013027 | Osi19 | 11321 | 1 | $4.2 \times 10^{-13}$ |
| LF013028 | Osi18 | 8381 | 5 | $1.9 \times 10^{-16}$ |
| LF015016 | Osi17 | 1406 | 0 | $2.8 \times 10^{-17}$ |

*Supplementary Figure 21: Approximate maximum likelihood phylogeny for holometabolan Osiris genes. All colored groupings have local support values > 0.9 except for Group A, Group C, and Osi22. Group names are from Smith et al. (in review) and Shah et al. (2012).*

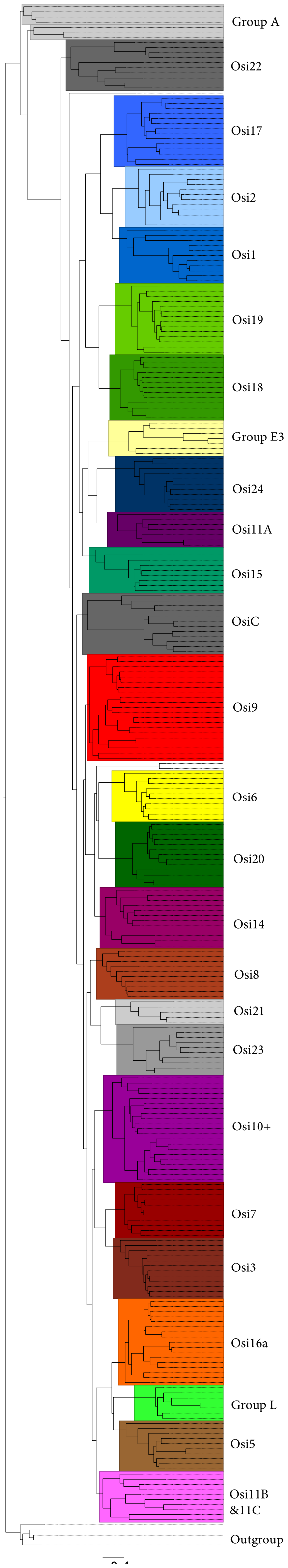

*Supplementary Figure 22: Scaffolds from the* A. ervi *genome assembly containing Osiris genes that are normally part of the conserved syntenic Osiris cluster. Most of these scaffolds are short and contain relatively few genes. The conserved synteny of this cluster in most insects suggests that these scaffolds may be contiguous. Color coding as in Supplementary Figure 22; non-black and numbered boxes are Osiris genes. Note difference in scaling of scaffolds.*

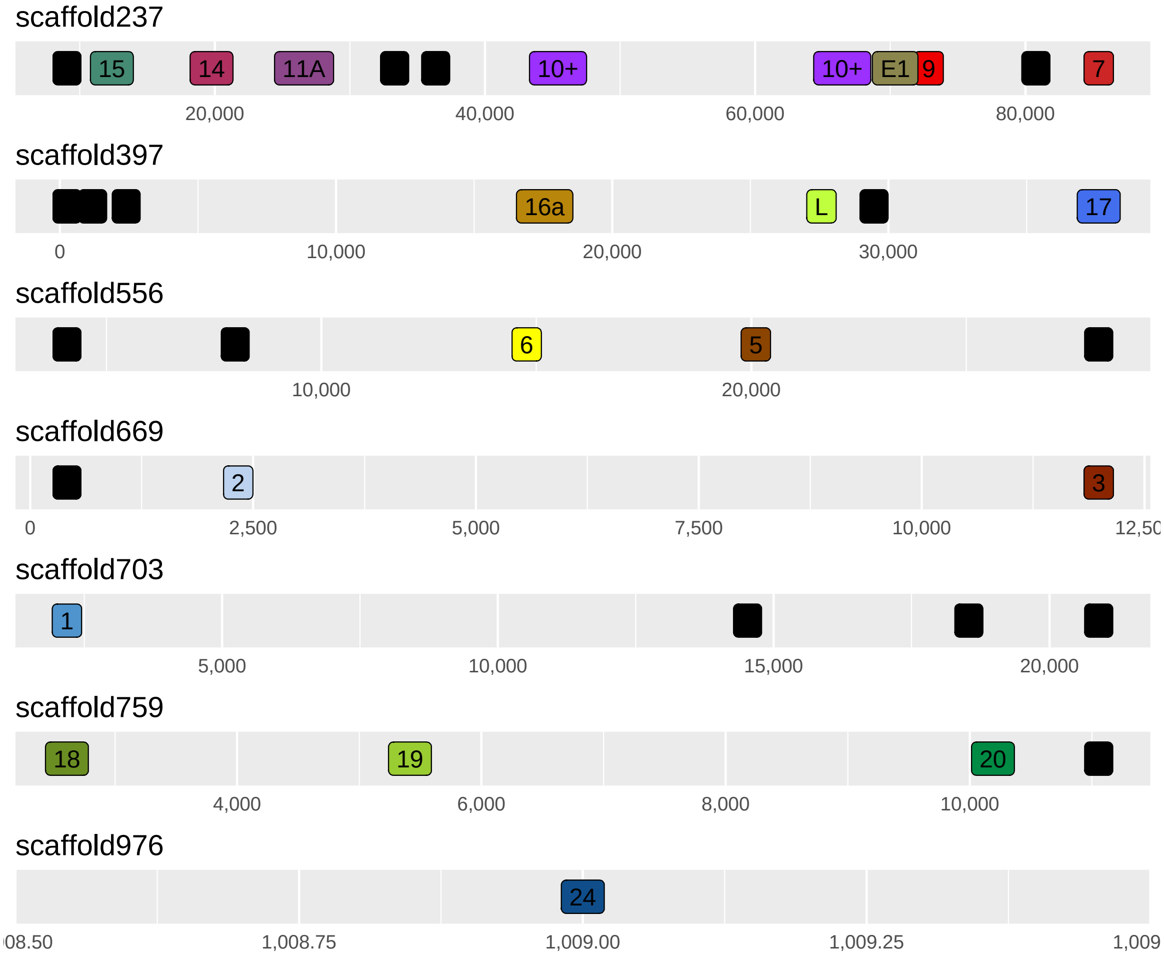

*Supplementary Figure 23: Scaffolds from the* L. fabarum *genome assembly, showing the Osiris genes that are normally part of the conserved syntenic Osiris cluster. As with* A. ervi*, the conserved synteny of these genes suggests that these relatively short scaffolds may be contiguous. Vertical separation of genes is to avoid overlap and has no biological meaning. Color coding is as in Supplementary Figure 22 and 23; non-black and numbered boxes are Osiris genes. Note differences in scaling of scaffolds.*

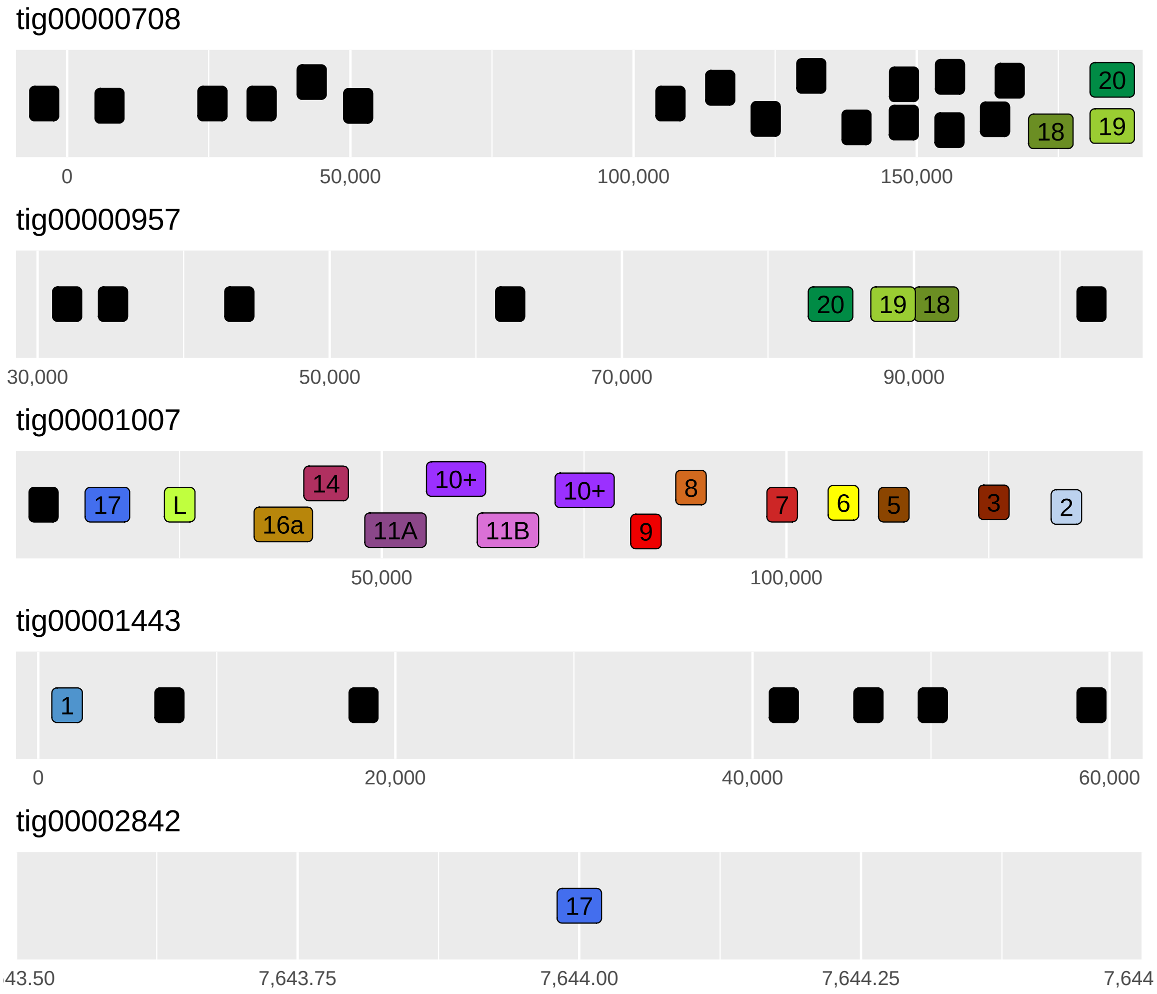

#### Oxidative phosphorylation (OXPHOS) genes

In most eukaryotes, mitochondria provide the majority of cellular energy (in the form of adenosine triphosphate, ATP) through the oxidative phosphorylation (OXPHOS) pathway. This pathway is composed of five protein complexes that utilize high energy electrons to produce a proton gradient across the inner mitochondrial membrane and the potential energy of this gradient is then used to phosphorylate ADP (adenosine diphosphate) into ATP. The OXPHOS pathway is conserved across eukaryotes and is unique because the complexes are comprised of both nuclear- and mitochondrial-encoded proteins. There are 13 protein-coding genes in nearly all eukaryotic mitochondrial genomes, all of which are used in the OXPHOS complexes, and there are ~67 "core" nuclear-encoded proteins that are used in these complexes (Porcelli *et al.* 2007). Unlike the mitochondrial-encoded genes, the nuclear-encoded genes vary in copy number across many organisms.

There were a total of 91 protein sequences of the nuclear-encoded oxidative phosphorylation genes from *Drosophila melanogaster* in the MitoDrome dataset (D'Elia *et al.* 2006). Of these 91 total genes, 20 are *Drosophia* duplications of some of the remaining 71 "core" genes. We found 69 of these 71 core genes in the *A. ervi* genome, with four of them duplicated and one with three copies, making a total of 75 nuclear-encoded oxidative phosphorylation genes (Supplemental Table 13). In the *L. fabarum* genome, we found 69 of these 71 genes, with five of them duplicated, for a total of 74 nuclear-encoded oxidative phosphorylation genes (Supplemental Table 13). The gene sets of *A. ervi* and *L. fabarum* contained the same genes, and the same genes were duplicated in each, implying duplication events that occurred prior to the split from their most recent common ancestor. One of these duplicated genes appears to be duplicated again in *A. ervi*, or the other copy has gone missing in *L. fabarum*.

In addition to the duplications inferred above, additional duplications were found that are likely errors in the assembly. The *L. fabarum* genome also appears to have many scaffolds with portions of nearly identical sequence. The gene models in these regions encode identical proteins and these may represent sequence assembly errors. There were 16 models that showed this pattern, 14 had one extra identical copy and two had two extra identical copies. *A. ervi* also had two gene models with this same pattern with one extra identical copy. Future work should resolve these duplications in the assembly.

*Supplementary Table 13: Summary of annotated OXPHOS genes in both genomes*

| | *A. ervi* | *L. fabarum* |
|---|---|---|
| Total genes (including duplicates) | 75 | 74 |
| Core OXPHOS genes | 69 | 69 |
| Missing "core genes" | 2 | 2 |
| Duplicated in *A. ervi* and *L. fabarum* | 5 | 5 |
| Unique duplicates | 1 | 0 |
| Models with identical copies (possible assembly errors) | 2 | 16 |

#### Chemosensory Genes- Ionotropic receptors (IRs):

*Supplementary Figure 24: Bootstrap tree built from predicted Ionotropic receptors (IRs) from* A. ervi *and* L. fabarum *(this study),* Apis melifera, Nasonia vitripennis*, and Diachasma alloeum. Rooted using the well-conserved IR co-receptors (including Ir8a and IR25a). Branches with 90% bootstrap support are indicated by grey dots. Naming follows the convention described by Croset* et al. *(2010) for those with unclear orthology (e.g AerviIR101).*

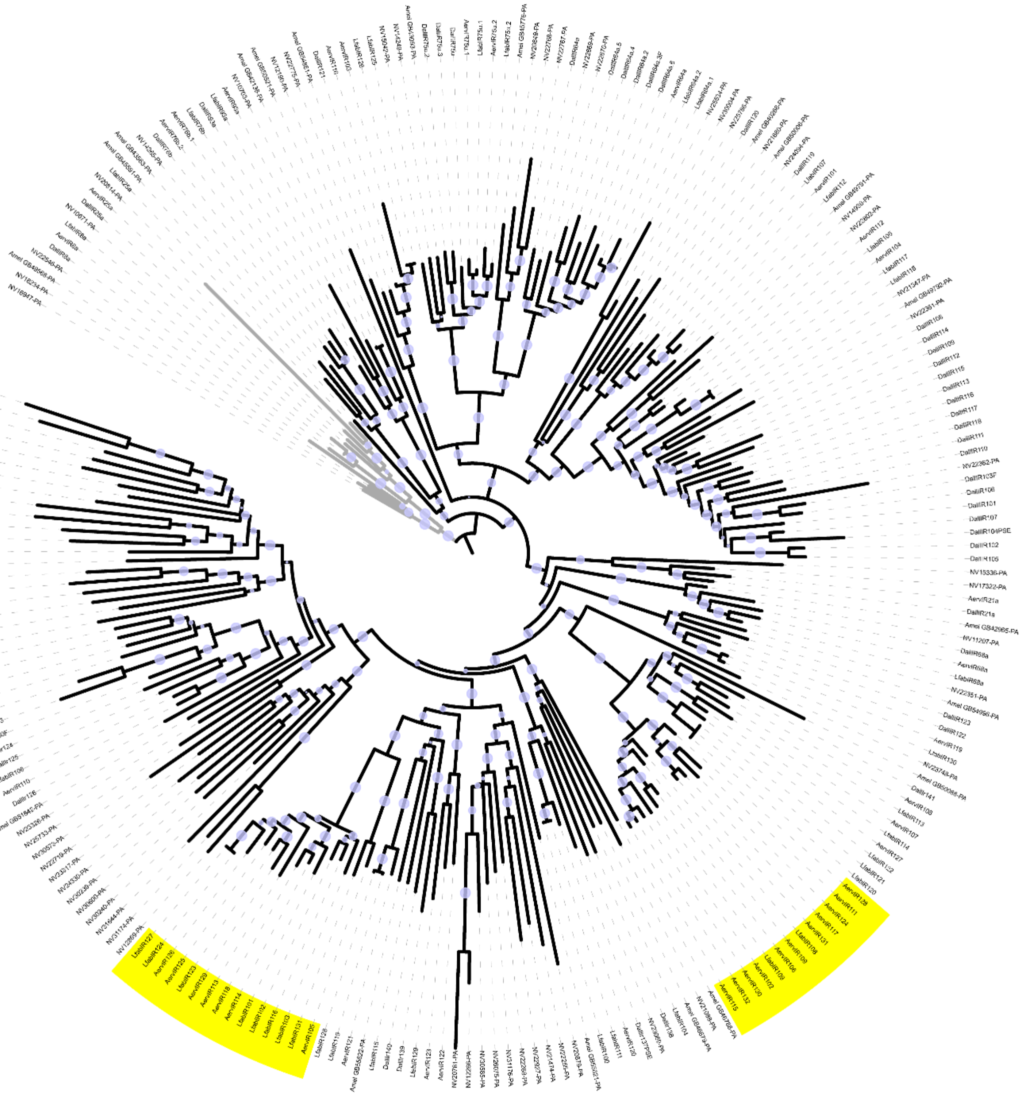

Sex Determination

*Supplementary Table 14: Core sex determination genes*

| Core sex determination genes | | | | | |
|---|---|---|---|---|---|
| *Species* | *Gene* | *Scaffold* | *plus/minus* | *Region (approx)* | *E-value of hit with Asobara homolog* |
| *A. ervi* | transformer | scaffold17 | plus | 10000-15000 | $4.0 \times 10^{-5}$ |
| *A. ervi* | transformerB | scaffold2824 | minus | whole scaffold* | 1.7 |
| *A. ervi* | transformer-2 | scaffold259 | plus | 33000-35000 | $3.0 \times 10^{-27}$ |
| *A. ervi* | doublesex | scaffold43 | minus | 236000-225000 | $2.0 \times 10^{-31}$ |
| *L. fabarum* | transformer | tig00000389 | plus | 105000-110000 | 0.008 |
| *L. fabarum* | transformer-2 | tig00001999 | plus | 60000-62000 | $4.0 \times 10^{-27}$ |
| *L. fabarum* | doublesex | tig00000015 | plus | 14000-25000 | $2.0 \times 10^{-30}$ |

* Incomplete, only CAM-domain plus RS/P-rich regions, first (more conserved half) of gene not on scaffold

*Supplementary Table 15: Annotation of genes related to sex determination. *This second homolog of RBP1 on the same scaffold mirrors what is seen in the* Microplitis *and* Fopius *genomes.*

| *Genes related to sex determination* | | | | | |
|---|---|---|---|---|---|
| *Species* | ***Gene*** | ***Scaffold*** | ***plus/minus*** | ***Region (approximate)*** | ***e-value (to* Nasonia *homolog)*** |
| ***A. ervi*** | *fruitless* | scaffold24 | plus | 589500-613000 | $1.0 \times 10^{-91}$ |
| ***A. ervi*** | sex-lethal homolog | scaffold3 | plus | 151000-152500 | $4.0 \times 10^{-63}$ |
| ***A. ervi*** | *CWC22 (aka nucampholin)* | scaffold18 | minus | 122000-119500 | 0 |
| ***A. ervi*** | *CWC22* paralog | scaffold18 | minus | 421000-418000 | 0 |
| ***A. ervi*** | *RBP1* homolog | scaffold39 | minus | 212500-213500 | $5.0 \times 10^{-39}$ |
| ***A. ervi*** | *RBP1* homolog* | scaffold39 | minus | 204700-205300 | $3.00 \times 10^{-31}$ |
| ***L. fabarum*** | *fruitless* | tig00000057 | minus | 321900-307000 | $4.00 \times 10^{-88}$ |
| ***L. fabarum*** | sex-lethal homolog | tig00001640 | minus | 6500-5000 | $5.00 \times 10^{-62}$ |
| ***L. fabarum*** | *CWC22 (aka nucampholin)* | tig00000545 | plus | 153000-155400 | 0 |
| ***L. fabarum*** | *RBP1* homolog | tig00000173 | minus | 429000-429300 | $4.00 \times 10^{-39}$ |
| ***L. fabarum*** | *RBP1* homolog* | tig00000173 | minus | 420000-420500 | $6.00 \times 10^{-38}$ |

#### Validation of DNA methylation genes

We confirmed these low levels of methylation in *A. ervi* by mapping this previously generated bisulfite sequencing data (Bewick *et al.* 2017) to our genome assembly. Since this data originated from a strain that is genetically differentiated from that which we have sequenced, we allowed for up to 13 alignment mismatches per read between their sequences and ours (Supplementary Figure 26).

Mapping the whole genome bisulfite sequencing data to the reference genome while allowing for 13 alignment mismatches retained 625,765 reads (80.94% of all raw reads, 94.5 Mbp). Sequence coverage was low: 63,554 of all methylation-available cytosines in the reference genome were covered by at least one read, and only 1,216 sites by more than one read (see figure 2). Of all methylation-available cytosines, 63,409 sites were never methylated, 143 sites were always methylated, and two were variably methylated. These methylated sites were roughly equally distributed among methylation-available cytosine classes: CG, CHG and CHH sites were methylated at a rate of 0.154%, 0.179%, and 0.210%, respectively. Because very few sites were methylated >2 times, we did not have the power to estimate variation in per-site methylation.

*Supplementary Table 16: Genes related to methylation. No homolog was detected for DNMT1 in either genome.*

| *DNA methylation genes* | | | | | |
|---|---|---|---|---|---|
| *Species* | ***Gene*** | ***Scaffold*** | ***plus/minus*** | ***Region (approx)*** | ***e-value hit (to Nasonia homolog)*** |
| ***A. ervi*** | *EEF1AKMT1* homolog | scaffold94 | minus | 144000-145500 | 1.00E-66 |
| ***A. ervi*** | *DNMT3* | scaffold45 | plus | 581000-585100 | 5.00E-138 |
| ***L. fabarum*** | *EEF1AKMT1* homolog | tig00000449 | plus | 13300-14100 | 5.00E-63 |
| ***L. fabarum*** | *DNMT3* | tig00002022 | plus | 68000-70600 | 9.00E-117 |

### Extended methods

### Assemblies

We made whole-genome alignments between the *L. fabarum* and *A. ervi* genomes using NUCmer, which is part of the Mummer 2.0 software package. Alignments were made using the default settings (Kurtz *et al.* 2004), with the six *L. fabarum c*hromosomes and unincorporated scaffolds (1,407 pieces in all) used as the reference genome. To remove potentially erroneous matches and better view the alignment, we filtered to retain only the *A. ervi* scaffolds >1Mbp, and only including instances with three or more consecutive matches. These filtered matches were then visualized using the program Circos (Krzywinski *et al.* 2009).

### GC content

##### Nitrogen and carbon content

Carbon and nitrogen content were quantified separately for each gene in the CDS using the PROTPARAM online platform (Gasteiger *et al.* 2005), invoked via a perl script (Hussain 2016). These outputs were further manipulated in R (R-Core-Team 2012), and visualized by graphing the first two components of a Principal component analysis, or PCA (R packages factoextra, reshape, and ggplot2, Kassambara & Mundt 2016; Wickham 2007, 2009).

##### Differential expression analysis: larvae vs. adult *L. fabarum*

To examine the GC content of transcripts that are expressed at different life-history stages, we compared expression in adult and larval *L. fabarum*. This previously generated RNA-seq data was also utilized in the gene predictions for *L. fabarum*, and is available to view in the online genome server, hosted by bipaa (bipaa.genouest.org/is/parwaspdb). This RNA-seq data (all 100-cycle, single end sequencing on an Illumina HiSeq2500) was generated from twelve lineages of parasitoid that had undergone experimental evolution on aphid hosts possessing different strains of *H. defensa* (as described in: Dennis *et al.* 2017). In both adults and larvae, the exact same lineages and conditions were sampled, collected in the course of an experimental evolution project at generations 11 (adult) and 14 (larvae). A total of 67 transcriptomic libraries were compared. Of these, 43 libraries came from pools of adults, sampled as 12-24h old virgin females (NCBI SRA PRJNA290156, Dennis *et al.* 2017). Larvae were part of a dual transcriptomic study, and were sampled 3-4 days after oviposition; we only used the 24 samples from the study that were deemed successful infections (NCBI SRA Accessions: SAMN10024115- SAMN10024165. Larval success described in: Dennis *et al.* in revision). In all cases, raw data was quality filtered and trimmed of potential Illumina primers as in the publications, using Trimmomatic (Bolger *et al.* 2014), and was mapped to the *L. fabarum* draft genome using STAR in the “quantMode” (Dobin *et al.* 2013). Differentially expressed genes between adults and larvae were identified using DESeq2 (Love *et al.* 2014). To achieve this, we built a model in which samples were blocked by replicate population (4 per treatment), aphid host, and sampling environment (*H. defensa* free, or not, see: Dennis *et al.* 2017), taking into account the replicate cages from the experiment. The full model was ~ age + age:tmt + tmt+ host + cage. Within this, we used a pairwise contrast between the two ages (larvae vs adult) to identify genes with significantly higher expression at either stage (FDR < 0.05). The most highly expressed genes from these subsets were determined by ranking the assembled transcripts according to the total normalized counts for all libraries. We compared the GC content of all subsets of genes (differentially expressed or highly expressed) using a two-sided t-test, implemented in R.

### Gene family evolution

#### Orphan gene ID

We identified orphan genes as those for which we could not find orthologs in any other sequenced parasitoid genomes. To do this, we generated clusters of orthologous and paralogous genes by comparing the predicted genes (CDS) from the genomes of *A. ervi* and *L. fabarum* to the predicted genes from parasitoids in both the *Braconidae* and *Ichneumonidae* (*Diachasma alloeum*, *Fopius arisanus*, *Macrocentrus cingulum*, *Microplitis demolitor* and *Nasonia vitripennis*). This was done using OrthoFinder (Emms & Kelly 2015) which is based on the analysis of pairwise sequence similarity scores obtained from an all-vs-all BLAST alignment among gene sequences between different species to identify orthogroups of genes. OrthoFinder produces a set of "Unassigned" genes that were not assigned to any orthogroup. We further examined these in two steps. First, we identified the genes that were present in the other genome assembled here using blastp (e-value < 0.001). Second, we identified species-specific genes, which we are calling orphan genes, by removing all genes that had hits to any other genes in the NCBI database. For this we identified matches at the protein level to the *nr* database, Swissprot database (using blastx), and the *nt* database (blastn), and at the DNA level against the *nt* database. In all cases, the e-value cut-off was 1e-10, and the databases were updated in June 2019. Within these putative orphans, we only retained those with transcriptomic support, based on the read counts generated for the GC analysis. We retained predicted genes of all lengths, as long as they had transcriptomic support, because manual inspection of some of these predicted genes suggested that they belonged to a region that actually contained a longer gene. The fasta file of the putative orphan genes for each species is available as Supplemental data (2 and 3).

### Venom proteins

#### Identification of *L. fabarum* venom proteins

For proteomic analysis, ten venom glands (see Supplemental Figure 14) were obtained by traction of the female ovipositor, isolated and dilacerated in 20 µl of Insect Ringer (KCl 182 mM; NaCl 46 mM; CaCl2 3 mM; Tris-HCl 10 mM) supplemented with a protease inhibitor cocktail (S8830; Sigma). The extract was centrifuged at 15,000 x g at 4°C for 10 min and the supernatant mixed with 4x Laemmli reducing buffer (Laemmli 1970) and loaded on a 12.5% SDS-PAGE (Supplemental Figure 15). The 16 most visible bands (numbered from the heavier to the lighter, Supplemental Figure 15) after silver staining (Morrissey 1981), were cut and sent for mass spectrometry. The 1D bands were treated with trypsin (Sequencing grade, Sigma) and alkylated reduced as previously described (Colinet *et al.* 2013). Samples were analyzed by mass spectrometry using a hybrid Q-Orbitrap mass spectrometer (Q-Exactive, Thermo Fisher Scientific, United States) coupled to a nanoliquid chromatography (LC) Dionex RSLC Ultimate 3000 system (Thermo Fisher Scientific, United States). Samples were trapped with a C18 PepMap 300 trap column (300 µm × 5 mm, C18, 5 µm, 300 Å) and desalted with solvent A (water with 0.1% formic acid) for 3 min at a flow rate of 20 µL/min. Peptide separation was performed on an Acclaim PepMap RSLC capillary column (75 µm × 15 cm, nanoViper C18, 2 µm, 100 Å) at a flow rate of 300 nl/min. The analytical gradient was run with various percentages of solvent B (acetonitrile with 0.1% formic acid) in the following manner: (1) 2.5–25% for 57 min, (2) 25–50% for 6 min, (3) 50-90% for 1 min, and (4) 90% for 10 min. Mass spectrometry (MS) spectra were acquired at a resolution of 35,000 within a mass range of 400–1,800 m/z. Ion accumulation was set at a maximum injection time of 100 ms. Fragmentation spectra of the 10 most abundant peaks (Top10 method) were acquired with high-energy collision dissociation (HCD) at a normalized collision energy of 27%.

All raw data files generated by MS were processed to generate mgf files and searched against (i) *L. fabarum* proteome predicted from genome (*Lysiphlebus fabarum* annotation v1.0 proteins) as well as (ii) *L. fabarum* transcriptome (Dennis *et al.* in revision; Dennis *et al.* 2017) using the MASCOT software (Perkins *et al.* 1999). Search parameters were as follow: variable modifications: methionine oxidation and carbamidomethyl-cysteine, mass tolerance: 10 ppm on parent ion and 0.02 Da on fragment ions, and a maximum of two tryptic missed cleavages.

#### Sequence annotation and analysis

To identify similarities with known proteins, comparisons with NCBI non-redundant protein sequence database were performed using blastp with a cut-off e-value of 1e-7 and a cut-off identity of 30%. Signal peptide prediction was performed with SignalP (Emanuelsson *et al.* 2007; Nielsen 2017). Identification of homologous venom proteins between *A. ervi* and *L. fabarum* was performed using blastp with a cut-off e-value of 1e-7 and a cut-off identity of 30%. Search for protein domains was performed using PfamScan (Finn *et al.* 2013). Identification of venom protein genes was performed using BLAST tools in Apollo.

Identification of *A. ervi* and *L. fabarum* non-venomous γ-GT proteins was performed using *N. vitripennis* γ-GT sequences as queries in blastp and tblastn searches against *A. ervi* and *L. fabarum* proteomes predicted from the genomes (*Aphidius ervi* annotation v3.0 proteins and *Lysiphlebus fabarum* annotation v1.0 proteins) and *A. ervi* (Ballesteros *et al.* 2017) and *L. fabarum* transcriptomes respectively. Multiple amino acid sequence alignments of GGT sequences were performed using MUSCLE (Edgar 2004). Phylogenetic analysis of GGT amino acid sequences was performed using maximum likelihood (ML) with PhyML 3.0 (Guindon & Gascuel 2003). SMS was used to select the best-fit model of amino acid substitution for ML phylogeny (Lefort *et al.* 2017).

### Community annotation

#### Osiris genes

Osiris gene orthologs in *A. ervi* and *L. fabarum* were determined with a two-part approach, candidate gene categorization followed by phylogenetic clustering. Candidate Osiris genes were generated using multiple, complementary methods, hidden Markov model searching, (HMMER3.1b2, Wheeler & Eddy 2013), and local alignment searching (BLAST, Altschul *et al.* 1990). A custom HMM was derived using all 24 well annotated and curated Osiris genes of *Drosophila melanogaster*. Next, an HMM search was performed on the *A. ervi* and *L. fabarum* proteomes, extracting all protein models with $P < 0.05$. Similarly, all *D. melanogaster* Osiris orthologs were searched in the annotated proteomes of *A. ervi* and *L. fabarum* using protein BLAST ($e < 0.05$). The top BLAST hit for each ortholog was then searched within each parasitoid genome for additional paralogs ($e < 0.001$). All unique candidates from the above approaches were then aligned using MAFFT (Katoh & Standley 2013), and an approximate maximum-likelihood phylogeny was constructed using FastTree (Price *et al.* 2009) via the CIPRES science gateway of Xsede (Miller *et al.* 2015). For simplicity, only Osiris orthologs from selected Holometabola were used, mostly other species of Hymenoptera. The species used were: the fruit fly (*D. melanogaste*r), the tobacco hornworm moth (*Manduca sexta*), the silkworm moth (*Bombyx mori*), the flour beetle (*Tribolium castaneum*), the jewel wasp (*Nasonia vitripennis*), the honeybee (*Apis mellifera*), the buff tail bumble bee (*Bombus terrestris*), the red harvester ant (*Pogonomyrmex barbatus*), the Florida carpenter ant (*Camponotus floridanus*), and Jerdon's jumping ant (*Harpegnathos saltator*).

To examine expression in Osiris genes, we used the read-counts that were generated using STAR mapping against the whole genome, as part of the GC analysis (see above). To get an understanding of general level of expression, we compared raw reads that mapped to the putative Osiris genes in both species. To further explore differences between adults and larvae, we looked at differential expression between the 59 previously generated adult RNA-seq libraries and 51 previously generated larval libraries (4-5 day old larvae). As with the read-mapping, this DE analysis was part of the same analysis used to compare larvae and adults for the GC analysis.

### OXPHOS

Annotation of genes involved in the oxidative phosphorylation pathway (OXPHOS) was performed in several steps. Initial blasts were performed on the protein level, and matched predicted genes from the two genomes to a set of nuclear-encoded OXPHOS proteins from *Nasonia vitripennis* (Gibson *et al.* 2010; J. Gibson unpublished) and a similar set of proteins from *Drosophila melanogaster*, downloaded from the MitoComp website (www.mitocomp.uniba.it, Porcelli *et al.* 2007). Matches to *N. vitripennis* were taken preferentially and mismatches between the *N. vitripennis* and *Drosophila* matches were manually investigated. After this, we searched for OXPHOS genes that were not identified in the predicted proteins for *A. ervi* and *L. fabarum* by blast-searching the *N. vitripennis*/*Drosophila* protein against the entire genome sequences. This was used as evidence to build new gene models.

With all possible OXPHOS genes identified in the two genomes, gene models in *A. ervi* and *L. fabarum* were used to improve one another. This was done using available expression evidence (typically more for *A. ervi*). The protein models from both species were then aligned to one another and to *N. vitripennis* to find missing or extraneous sections. These were also compared to other hymenopteran proteins. Lastly, annotated proteins were blast-matched back to the *N. vitripennis* genome, to ensure they were reciprocal-best-blast hits.

Genes were named according to the existing *N. vitripennis* nomenclature (which has been extensively curated with NCBI to ensure the *Nasonia* naming was as consistent as possible, J. Gibson pers comm.) This is important to note, because the OXPHOS genes have many different names in different organisms (sometimes 5+ names). These were all added as synonyms.

To detect duplications and assembly errors, the gene models in the blast results to *N. vitripennis* were inspected beyond the top match, with additional inspection to ensure that it was an OXPHOS gene. True gene duplications were identified when there were two similar copies in one (or both) genome(s). These genes were given the same gene name, with A or B at the end, and the same gene symbol with A or B appended. Duplicates that were more similar to the *N. vitripennis* gene were given the A designation. There were many cases in the *L. fabarum* genome where two models had the exact same blast results to a given OXPHOS gene. Further investigation of the underlying genomic sequence showed that these were either extremely similar, or identical. These are likely assembly errors and differ by just a few SNPs. These were given identical gene names and gene symbols, are annotated as “Alleles” A and B, and are listed as “duplications” in the Supplemental Data 11. Further investigation should resolve true duplications from probable sequencing errors, and this suggests further refinement could benefit other gene models as well.

### Validation of DNA methylation genes

To verify the low DNA methylation level of 0.5% reported by Bewick *et al*. (2017) and suggested by the absence of DNMT1, we mapped their low-coverage whole genome bisulfite sequencing data from *A. ervi* (NCBI Short Read Archive Accession GSE83497) against the *A. ervi* reference genome. This data set contains 773,136 unpaired short reads of 151bp length (116.7 Mbp in total). There was no information available on the wasp strain used in this study, however, most likely it was not the same as the strain from which the genome was sequenced. In other words, some genetic divergence should be expected between these two data sets.

Reads were mapped to the *A. erv*i genome with BS-Seeker2 (Guo *et al.* 2013), a full pipeline explicitly developed for mapping bisulfite sequencing data. Initially we ran BS-Seeker2 with default settings, which allows for a sequence mismatch of 4bp between reads and reference genome during mapping. The resulting ‘mappability’ (the percentage of all reads that map to a unique location on the genome) of the data was lower than what would be expected for data of the same species (62.79%). In order to improve the mappability of the data, we repeated the analysis for a range of settings for allowed mapping mismatches (0 to 8 inclusive). Very low mappability is expected at zero mismatches since sequencing error introduces per read a few mismatches between read sequence and reference genome. A preliminary analysis of the output showed that allowing for up to 8 mismatches per read still improved the mappability considerably, and hence the parameter range of allowed mismatches was expanded up to 20 mismatches and the analyses repeated accordingly. At higher number of allowed mismatches the number of uniquely aligned reads asymptotically approached the total number of reads that have a single hit on the genome (Supplemental Figure 23).

Interestingly, the number of reads that have multiple hits on the genome does not change with varying numbers of allowed alignment mismatches (33,793 reads, i.e. 4.37%, Supplemental Figure 23). Note that reads having multiple matches are not informative on DNA methylation levels and hence these are automatically excluded from the analysis. Allowing for a higher number of mapping mismatches might introduce mapping hits on the reference genome that are false positives. Instead, we chose to aim for 95% inclusion of the reads that have a unique hit to the genome, and not to further increase this parameter for the number of allowed mismatches. This percentage was reached when allowing for 13 mismatches, and therefore this number was chosen for downstream analysis of DNA methylation levels.

*Supplementary Figure 25: Summary of bisulfite sequencing against the* A. ervi *genome. Mapping success was evaluated across a range of allowed mismatches, and 13 was chosen to obtain at least 95% inclusion of the data.*

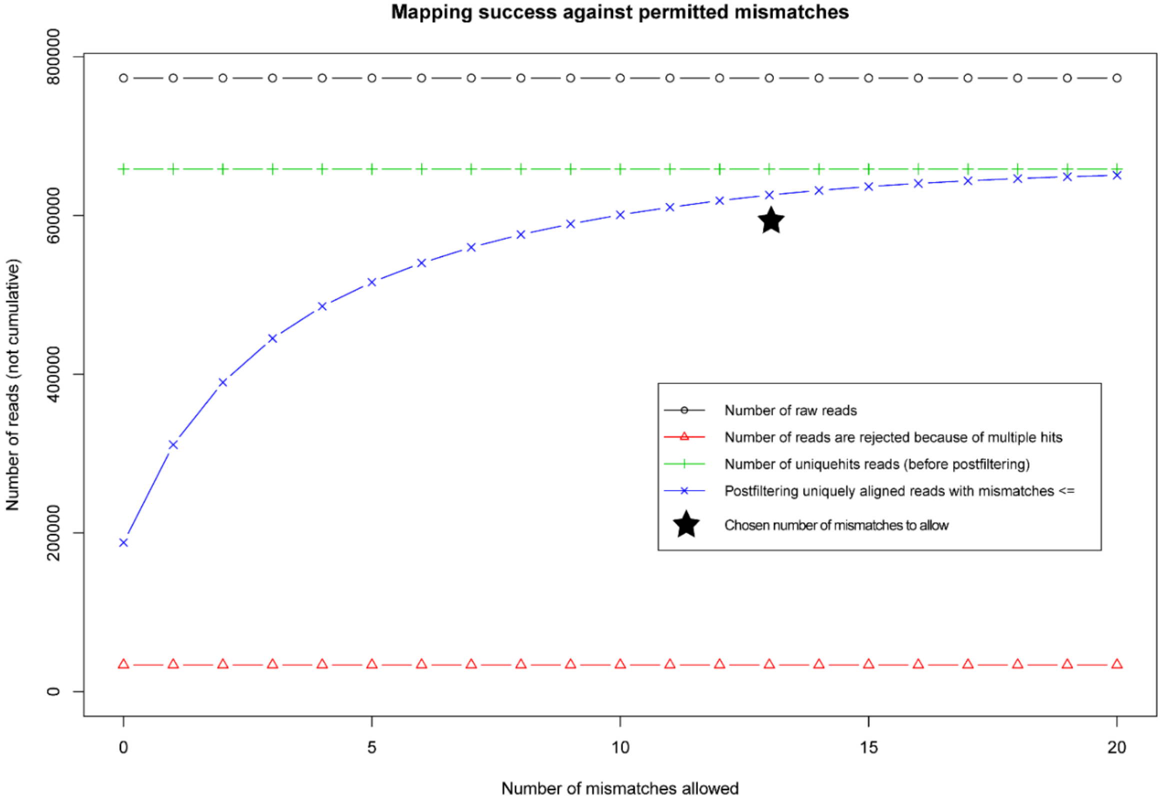
